# Two antagonistic gene regulatory networks drive Arabidopsis root hair growth at low temperature

**DOI:** 10.1101/2024.04.09.588718

**Authors:** Tomás Urzúa Lehuedé, Victoria Berdion Gabarain, Miguel Angel Ibeas, Hernan Salinas-Grenet, Romina Acha, Tomas Moyano, Lucia Ferrero, Gerardo Núñez-Lillo, Jorge Perez, Florencia Perotti, Virginia Natali Miguel, Fiorella Paola Spies, Miguel A. Rosas, Ayako Kawamura, Diana R. Rodríguez-García, Ah-Ram Kim, Trevor Nolan, Adrian A. Moreno, Keiko Sugimoto, Norbert Perrimon, Karen A. Sanguinet, Claudio Meneses, Raquel L. Chan, Federico Ariel, Jose M. Alvarez, José M. Estevez

## Abstract

The root hair (RH) cells can elongate to several hundred times their initial size, and are an ideal model system for investigating cell size control. Their development is influenced by both endogenous and external signals, which are combined to form an integrative response. Surprisingly, a low temperature condition of 10°C causes an increased RH growth in *Arabidopsis* and in several monocots, even when the development of the rest of the plant are halted. Previously, we demonstrated a strong correlation between the RH growth response and a significant decrease in nutrient availability in the medium under low temperature conditions. However, the molecular basis responsible for receiving and transmitting signals related to the availability of nutrients in the soil, and their relation to plant development, remain largely unknown. In this study, we have discovered two antagonic gene regulatory networks (GRNs) controlling RH early transcriptome responses to low temperature. One GNR enhances RH growth and it is commanded by the transcription factors (FTs) *ROOT HAIR DEFECTIVE 6* (RHD6), *HAIR DEFECTIVE 6-LIKE 2 and 4* (RSL2-RSL4) and a member of the homeodomain leucine zipper (HD-Zip I) group I 16 (AtHB16). On the other hand, a second GRN was identified as a negative regulator of RH growth at low temperature and it is composed by the trihelix TF *GT2-LIKE1* (GTL1) and the associated DF1, a previously unidentified MYB-like TF (AT2G01060) and several members of HD-Zip I group (*AtHB3, AtHB13, AtHB20, AtHB23*). Functional analysis of both GRNs highlights a complex regulation of RH growth response to low temperature, and more importantly, these discoveries enhance our comprehension of how plants synchronize the RH growth in response to variations in temperature at the cellular level.

**Significance Statement:** Root hair (RH) cells may expand hundreds of times, making them a useful cell size model. Integrated endogenous and exogenous cues affect their development. Even when plant development ceases, Arabidopsis and other monocots grow RH at 10°C. We previously established a strong correlation between growth response and a substantial medium nutrition reduction at low temperature. Receiving and transmitting soil nutrient signals and their impact on plant development are unknown molecularly. Our study identified two opposing gene regulatory networks (GRN) that govern early transcriptome responses linked to RH growth at low temperature. Functional analysis shows a complex regulation of the transcriptional cascade to influence the low-temperature RH growth. These results explain how cells coordinate RH formation in response to temperature.

## Introduction

Root hairs (RHs) are single cells that develop as a tubular protrusion from the root epidermis in a mixed polar and non-polar manner up to several hundred times their original size (Yi et al. 2010: Mangano et al. 2016; Herburger et al. 2022; Jia et al. 2023). RH differentiation is a complex process and consists of 3 stages: differentiation, initiation and elongation. The first is controlled by a developmental program involving transcription factor complexes (TFs) allowing or repressing the expression of the GL2 (GLABRA2) protein. This protein blocks RH development (Lin *et al*., 2015) by inhibiting the transcription of the RHD6 (*ROOT HAIR DEFECTIVE 6*) regulator, which together with RSL1 can induce the expression of other factors such as RSL2 (Root Hair Defective 6 like 2) and RSL4 (*ROOT HAIR DEFECTIVE 6 LIKE 4*), thus allowing RH growth (Menand *et al*., 2007; Yi *et al*., 2010; Datta *et al*., 2015; Proust *et al*., 2016; Vijayakumar *et al*., 2016; Lopez *et al*., 2023). RHs are able to grow within hours in order to reach water-soluble nutrients in the soil, to promote interactions with the local microbiome, and to support the anchoring of the root. The overall fitness of plants is reduced in various loss-of-function RH mutants that lack RHs when grown in challenging soil conditions (Wu *et al*., 2007, Haling *et al*., 2013, Miguel *et al*., 2015). On the other hand, genotypes with longer RHs, achieved through the overexpression of RSL transcription factors, have shown beneficial effects in species such as wheat (Han *et al*., 2016, 2017), rice (Moon *et al*., 2019), and Brachypodium (Kim & Dolan, 2016). This highlights the deep importance of RHs in relation to plant development and physiology, despite not always receiving full recognition. Intrinsic factors regulate RH elongation at the cellular level and external signals, among which we find morphogenetic programming, nutrient availability, and the action of different hormones (Casal & Estevez, 2021; Lopez *et al*., 2023). Several studies previously showed that RH growth is highly responsive to increasing nutrient concentrations in the media (Yi *et al*., 2010; Bhosale *et al*., 2018; Moison *et al*., 2021; Jia *et al*., 2023). While high nutrient concentrations impair RH development and growth, low concentrations of nutrients such as inorganic phosphate (Pi) and/or nitrates trigger a strong growth response mediated by the phytohormone auxin (Bhosale *et al*., 2018; Mangano *et al*., 2017; Jia *et al*., 2023; Lopez *et al*., 2023).

The activation of RH expansion, caused by nutrient unavailability, is known to occur via a process of transcriptional reprogramming controlled by RHD6 and other downstream transcription factors like RSL4 (Bhosale *et al*., 2018; Shibata *et al*., 2018; Shibata *et al*. 2022; Jia et al. 2023). The RH growth in wild-type (WT) plants shows that higher concentrations of nutrients (ranging from 0.5X to 2.0X of the Murashige and Skoog medium) hinder RH growth independently of the low temperature (Moison et al. 2021). In summary, low temperature (10°C) is able to enhance RH growth possibly as a result of the restriction of nutrient mobility and accessibility by the root (Moison *et al*., 2021; Martínez-Pacheco *et al*., 2021). Conversely, an increase in agar concentration in the medium that restrains nutrient mobility resulted in the reversion of low temperature-induced RH elongation (Moison *et al*., 2021). Altogether, these observations suggested that low temperatures restrict nutrient mobility and availability in the culture medium, leading to the promotion of polar RH growth. Despite these previous findings, the signals that trigger the RH cell elongation under these conditions are still unknown, but it is presumed that RHs could be highly susceptible to environmental stresses such as temperature changes, triggering differences at the transcriptomic and proteomic levels. Specifically, a novel ribonucleoprotein complex composed by the lncRNA AUXIN-REGULATED PROMOTER LOOP (APOLO) and the TF WRKY42 forms a regulatory hub to activate RHD6 by shaping its epigenetic environment and integrating low temperature signals governing RH growth (Moison *et al*., 2021; Martínez-Pacheco *et al*., 2021). Recently, new molecular components related to cell wall-apoplastic related peroxidases, PRX62 and PRX69, were identified as important factors for low temperature triggered RH growth by inducing changes in ROS homeostasis and cell wall glycoprotein EXTENSIN insolubilization (Martinez-Pacheco *et al*., 2022). An autocrine regulation pathway between Rapid Alkalinization Factor 1 (RALF1) and the *Catharanthus roseus* RLK1-like (CrRLK1L) FERONIA (FER) receptor phosphorylates the early translation factor eIF4E1 and produces RH-related proteins including RSL4 to promote RH formation (Zhu *et al*., 2020a,b). Finally, at low-temperature, RHs showed a physical link between FER and TARGET OF RAPAMYCIN COMPLEX (TORC). Mutants for *fer-4* and TORC components responded differently to low nitrate than Col-0, and RHs remained stunted throughout the bulge stage (Martinez-Pacheco *et al*., 2023a). This FER-TORC pathway connects nutrients perception at the cell surface with downstream responses in RH growth at low temperatures (Martinez-Pacheco *et al*., 2022). Overall, these results highlighted here summarizes several of the molecular components recently discovered from the plant cell surface perception to the transcriptional regulation involved in low temperature triggered RH growth.

Although advances were achieved in our understanding of how RH growth occurs at low temperature, the dynamic transcriptional cascade controlling RH growth remains largely unknown, providing us with the opportunity to find new key regulators in the process of RH elongation. Here, we provide a deep characterization of how two antagonistic gene regulatory networks (GRNs) are able to direct gene expression at early time points that allows RH growth to respond to the low temperature. We functionally validated these transcriptional nodes by gene expression assay, phenotypic characterization of mutant/overexpressor and evaluation of genetic interactions. These findings improve our understanding of how plants coordinate the development of their RHs in response to changes in temperature and nutrient availability at the cellular level.

## Results

### Low temperature effect on RH growth depends on nutrient availability

Root hair (RH) growth is a well conserved adaptive response to external low nutrients in the media-soil. Previously, it was shown that at low temperature growth conditions (10°C), nutrients in the media are reduced, which triggers a strong response in RH growth (Moison *et al*., 2021; Martinez-Pacheco *et al*., 2021). Then, we assessed the effect of the low temperature on RH growth, through a detailed temporal characterization of root development (**Figure 1A**). Starting at 2 days of 10°C, a significant effect on RH elongation compared to control conditions was detected. This induction of growth reached its peak at 3 days of low temperature treatment. Then, we decided to test if small temperature changes around 10°C also trigger a similar RH growth effect. We performed a temperature gradient from 22°C to 4°C in 2°C steps and maximum growth was detected at 8-10°C (**Figure 1B**). From 8°C down to 4°C, the low temperature had a negative effect on RH growth suggesting that the low temperature itself is more deleterious than the linked low-nutrient effect. Based on this, we tested if a medium with no nutrient content is able to repress the low temperature induced growt2s. To this end, we used agarose that lacks any detectable nutrients and salts and tested the low temperature growth effect (**Figure 1C**). In agreement with previous evidence, when no nutrient is in the medium no low temperature effect is observed on the RH growth, confirming that the signal is nutritional and not related to the temperature itself as suggested before (Moison *et al*., 2021; Martinez-Pacheco *et al*., 2021; Martinez-Pacheco *et al*., 2023). Our data allowed us to use the low temperature effect on RHs as a proxy of complex nutritional signals changes coming from the medium. When we evaluated the effect of different pretreatment growth times, it was observed that the final RH lengths in plants exposed to low temperatures were equivalent whether plants were pre-grown for 7 or 14 days at 22°C, which was also the case for their basal RH lengths (**Figure S1**). Since root light exposure generates a stress that affects growth, hormonal signaling, abiotic responses, or nutrient starvation adaptation compared with roots grown in the dark (Silva-Navas *et al*., 2016), we decided to test if direct light exposure might affect the RH growth intensity recorded before. For this purpose, we used the system called D-Roots originally developed to cultivate roots in darkness, including dark agar plugs and black-colored vertical plates (Silva-Navas *et al*., 2015). Using this D-root system, we have confirmed the previously measured RH phenotypes at low temperature although with slightly higher magnitude (**Figure 1C**). These results together showed that 8-10°C were the main temperature trigger of RH growth affecting nutrient accessibility to the root surfaces and light exposure does not significantly affect this growth process in RHs. Finally, we tested if this enhancement of RH growth at low temperature is also present in monocots (**Figure 1D**) including *Brachypodium distachyon* (Bd21), rice (Kitaake), and wheat (Chinese Spring). In these three cases we observed the same effect on RH growth at low temperature as in Arabidopsis, although in these cases we tested them using 16 h:8 h day:dark cycle (see methods). This last result highlighted that low temperature effect on RH growth might be a broader response in plant roots, also applicable to these monocots as important crops.

**Figure 1.**
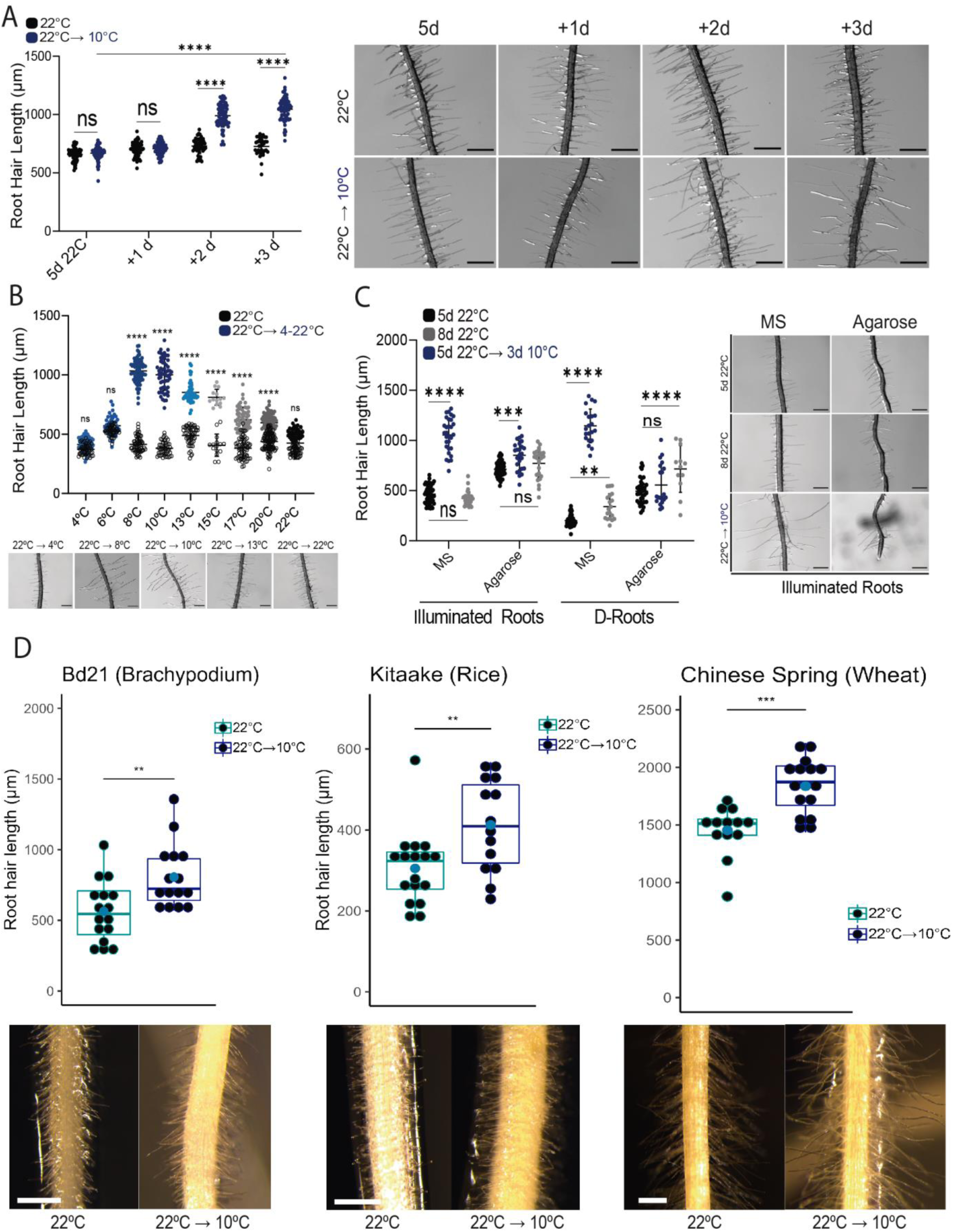
Low temperature effect on RH growth in Arabidopsis, Brachypodium, rice and wheat. (**A**) Low-temperature RH growth up to 3 days of treatment. (**B**) Temperature fluctuations impact on RH growth and growth maxima was detected at 8-10°C. (**C**) A nutrient-free medium can suppress the low-temperature growth impact on RHs. The previously recorded RH traits at low temperature were verified with somewhat larger magnitude using this D-root method. (A-C) Each point is the mean of the length of the 10 longest RHs identified in the maturation zone of a single root. Data are the mean ± SD (N= 30 roots), two-way ANOVA followed by a Tukey–Kramer test; (****) p <0.001, NS= non-significant. Results are representative of three independent experiments. Asterisks indicate significant differences between Col-0 and the corresponding treatment at the same temperature or between the same treatment at different temperatures. Representative images of each genotype are shown on the right. Scale bars= 500 µm. (**D**) Low-temperature triggers an enhancement on RH growth in the monocots *Brachypodium distachyon* (Bd21), rice (Kitaake), and wheat (Chinese Spring). Representative pictures are shown at the bottom. Scale bars= 500 µm. Data are the mean ± SD (N= 30 roots), two-way ANOVA followed by a Tukey–Kramer test; (**) p <0.01, (***) p <0.001, NS= non-significant.

### Genome wide transcriptional changes in roots exposed to low temperature

RSL4 and RSL2 have been identified as important regulators of RH growth in response both at 22°C and at low temperature, although RSL4 has been shown to be of higher relevance than RSL2 in this role (Yi *et al*., 2010; Moison *et al*., 2021, this study **Figure 3B**). Therefore, the loss of function of RSL4, and RSL2 to a lower extent, would lead to changes in the transcript abundance of direct and indirect RH gene targets. To identify all genes regulated by the transcription factor (FT) RSL4 in the context of low temperature response, and to find molecular factors involved in RH development that are dependent on RSL4, RSL2, or both, we analyzed the transcriptome of *rsl2*, *rsl4*, and *rsl2 rsl4* genotypes in response to low temperature treatments (2h and 6h) and compared it with the transcriptome of Wt ecotype Columbia-0 (Col-0) plants (**Figure 2A**). Differential expression analysis was performed and the number of genes was filtered considering an FC>|0.5| in at least one condition. From a total of 4,167 differentially expressed genes (DEG) identified between Col-0 and any of the mutant genotypes, only in *rsl4* there are 2,768 DEG (comprising 66.4% of the total change) while RSL2 controls only 102 genes (**Figure S2**). The heatmap in **Figure S3** shows the total number of DEGs identified, which were clustered into 5 groups (see also **Supplementary Table S1**). Clusters 2 and 3 only discriminate against low temperature effects independently of the genotypes while clusters 1 and 4 change based only on the RSL4 genotype but are independent of the low temperature. Cluster 5 was composed of 1,157 genes, which are positively regulated by RSL2 and RSL4, and is enriched with several GO terms related to RH development (**Figure S4**). Unfortunately, any of these 5 clusters respond to genotype and treatment (low temperature) at the same time. To identify genes whose expression depends on the interaction between low temperature and the genotypes, we performed a two-way analysis of variance (ANOVA) analysis as described previously (Alvarez *et al*., 2014), considering temperature (T) and each genotype (G). This analysis allows us to uncover genes for the interaction of TxG factors whose response to temperature is affected in the mutants analyzed. Our analysis identified 2,365 genes with a significant TxG factor for the *rsl4* mutant (**Figure 2B**). These results were consistent with RSL4 being more influential in the RH response to low temperature as compared to RSL2 (Moison *et al*., 2021) in agreement with previous evidence that show that RSL4 levels define the final RH length (Datta *et al*., 2016). By analyzing the heat map of these 2,365 DEG, we identified three main clusters regulated by RSL4 and also affected by low temperature (**Figure 2B**). GO analysis revealed several processes related to RNA, virus replication and cold. The expression of RSL4 (and the related RHD6) were increased at early time points of low temperature treatment while RSL2 was decreased (**Figure S4**). Taken together, these data indicate that the TF RSL4 is a major regulator of the early RH transcriptome at low temperature.

**Figure 2.**
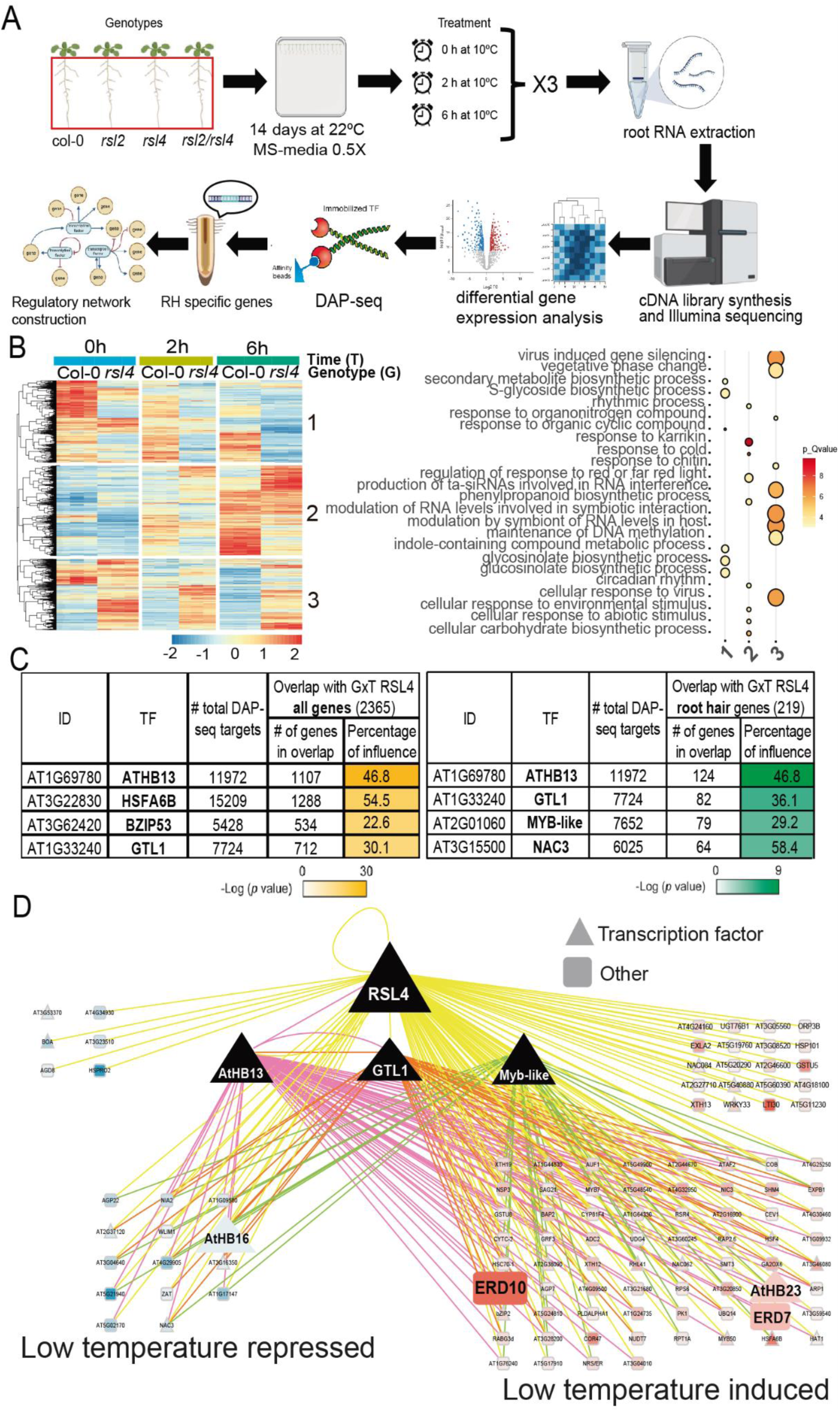
Identification of the RH specific RSL4-mediated gene regulatory network acting at low temperature on an early temporal window. (**A**) Dual bioinformatic-experimental strategies using RNA-seq plus DAP-seq prediction followed to identify the RH specific RSL4-transcriptional network at early times at low temperature. (**B**) Differential expression analysis by a heat map of the transcriptome of Wt Col-0 and *rsl4* genotypes in response to low temperature treatments (2h and 6h). A two-way analysis of variance (ANOVA) analysis was carried out, considering temperature (T) and each genotype (G) as factors and controlling type I error using the false discovery rate. A total of 2,365 DEG showed an altered response to low temperature in the *rsl4* mutant (significant TxG factor) were grouped into 3 clusters. (**C**) Best 4-ranked list of TFs based on the overall overlap of 2,365 genes controlled by RSL4 with DAP-seq common targets (on the left) and 219 genes are RSL4-specifically expressed in RHs (on the right) using the ConnecTF database. (**D**) Gene regulatory RSL4-network acting on RH growth at low temperature after 6h of exposure. Genes are drawn as triangles (TF, transcription factors) and squares (target genes). Lines indicate predicted transcriptional activation or predicted transcriptional repression. Yellow lines indicate RSL4, red line AtHB13 and green lines MYB-like putative TFs binding site by DAP-seq in the upstream region of the corresponding genes.

**Figure 3.**
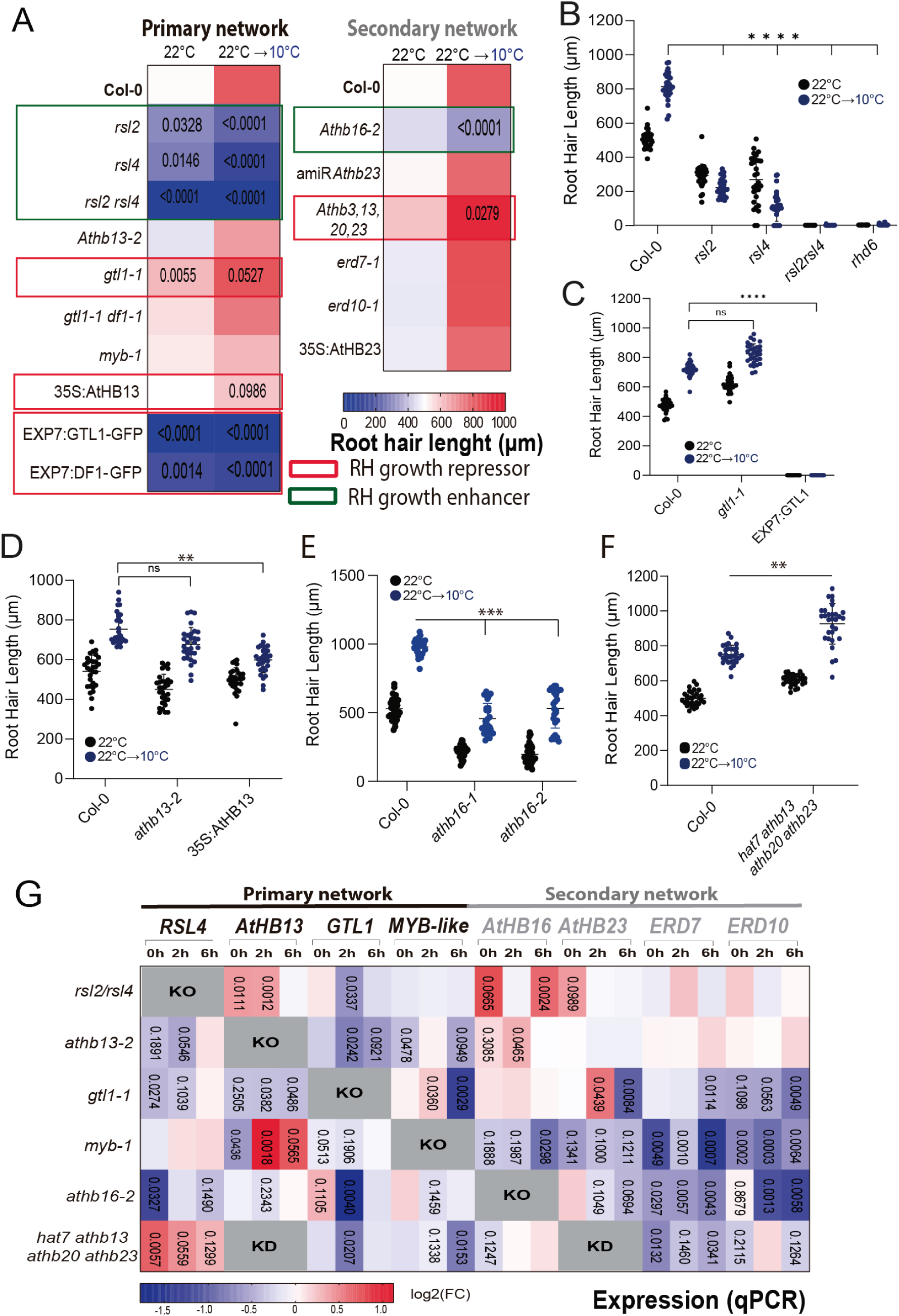
Characterization of major nodes of the RSL4-transcriptional primary and secondary network. (**A**) Heatmap of the RH phenotype at 22°C and 10°C of the different mutants and overexpression lines of the main network nodes RSL4 (and RSL2), AtHB13, GTL1/DF1 and MYB-like as well as the downstream selected components AtHB16/AtHB23 and ERD7/ERD10. Blue-Red color scale represents the length of RH in µm (0-+1000). In green boxes RSL4/RSL2 and AtHB13 are indicated as positive effectors of RH growth while in red boxes GTL1, DF1 and multiples AtHBs are shown as repressors of RH growth. Pair comparisons are made between Col-0 and each line (see **Supplementary Table S3** and **Figures S4-S9**). Data are the mean ± SD (N=30 roots), two-way ANOVA followed by a Tukey–Kramer test; only p-values below 0.20 are shown. (**B-F**) Selected phenotypes are shown. Each point is the mean of the length of the 10 longest RHs identified in a single root. Data are the mean ± SD (N=30 roots), two-way ANOVA followed by a Tukey–Kramer test; (*) p <0.05, (***) p <0.001, NS=non-significant. Results are representative of three independent experiments. Asterisks indicate significant differences between Col-0 and the corresponding genotype at the same temperature. (**B**) Scatter-plot of RH length of Col-0, *gtl1-1* mutant, and the overexpressor EXP7:GTL1-GFP grown at 22°C or at 10°C. (**C**) Scatter-plot of RH length of Col-0, *rsl2-1*, *rsl4-1* and *rhd6* grown at 22°C or at 10°C. (**D**) Scatter-plot of RH length of Col-0, *athb13-2* mutant and overexpressor 35S:AtHB13 grown at 22°C or at 10°C. (**E**) Scatter-plot of RH length of Col-0 and *athb16-1* and *athb16-2* mutants grown at 22°C or at 10°C. (**F**) Scatter-plot of RH length of Col-0 and the quadruple *athb3 (hat7) athb13 athb20 athb23* grown at 22°C or at 10°C. (**G**) Heatmap of the gene expression analysis of the main gene network nodes composed by the TFs RSL4, AtHB13 and GTL1/DF1 and as well as selected components ERD7/ERD10 and AtHB16/AtHB23 in each mutant line previously isolated and characterized (**Figures S10-S13**). Data are the mean of three biological replicates, two-way ANOVA followed by a Tukey–Kramer test; only p-values below 0.20 are shown.

### RSL4 controls low temperature response

To get insight into the regulatory cascade triggered by RSL4, we focused on the 2,365 genes that showed an altered response to low temperature in the *rsl4* mutant (significant TxG factor). To investigate which TF might mediate the RSL4 effect of gene expression in response to low temperature, we exploited the TF-target data allocated in the ConnecTF database (Brooks *et al*., 2021). The ConnecTF platform provides access to a library of experimentally confirmed TF–target gene interaction datasets. To this end, we used the list of differentially expressed TFs that depend on RSL4 and checked which TF has available validated *in vitro* TF-binding data obtained by DNA Affinity Purification sequencing (DAP-seq) (O’Malley *et al*., 2016). We then reduced the list of possible target genes to those 1,644 genes highly or uniquely expressed in RH cells previously identified by single cell RNA-seq or GFP-enriched protoplast studies (Brady *et al*., 2007; Denyer *et al*., 2019; Jean-Baptiste *et al*., 2019; Ryu *et al*., 2019; Shulse *et al*., 2019; Zhang *et al*., 2019) (**Supplementary Table S2**; **Figure 2C**). The global overlap between GxT controlled by RSL4 on the top ten TFs and at the same time with high expression on RHs gives 334 genes regulated by low temperature at 6h (**Supplementary Table S3-S4**) while the genes controlled only by RSL2 are 164 genes and by both RSL2 and RSL4 are 106 genes in the same conditions. Then, we selected the top three TFs of the best five-ranked list: (i) AtHB13, a class I HD-Zip TF previously described as involved in low temperature response but not related to RH growth (Cabello *et al*., 2012), (ii) a MYB-like (AT2G01060) and (iii) GTL1, a previously characterized negative regulator of RH growth under high nutrient condition, and its homologous DF1 (Shibata *et al*., 2018, 2022). Based on this analysis, a core RH gene regulatory network composed of 112 genes controlled by RSL4 specifically at low temperature was identified (**Figure 2D**; **Supplementary Table S5**). Downstream of these main regulators, we selected several members of the Arabidopsis HD-Zip I subfamily, including AtHB16/AtHB23 and two TFs *EARLY-RESPONSIVE TO DEHYDRATION* ERD7/ERD10 for further characterization. Notably, none of these TFs (unless RSL4) were previously linked either to RH growth or to low temperature. This emphasizes the prognostic capability of these integrated methodologies.

### Two antagonistic gene regulatory networks regulate RH growth at low temperature

To validate the function of these RSL4-regulated gene regulatory networks in RH growth at low temperature, we performed detailed RH phenotype screenings at 22°C and at 10°C in loss of function mutants and overexpression lines for each component of the identified gene regulatory regulatory network nodes (**Figures S5-S13**; **Figure 3A; Supplementary Table S6**). These include the TFs RSL4, AtHB13, GTL1/DF1 and MYB-like as well as selected components AtHB16/AtHB23 and ERD7/ERD10 (**Supplementary Table S6**) (**Figure 3A-B**). Not surprisingly, several of the TFs analyzed exhibited an altered RH growth at 10°C (**Figures S5-S9** and **Figures 3A-F**) providing strong evidence of the predictive power of this combined bioinformatics-experimental approach to identify key roles of specific TFs in certain environmental conditions. According to our functional screening, the main nodes RSL4 (and RSL2) and the downstream gene AtHB16 are positive regulators of RH growth at low temperature while, on the other hand, GTL1-DF1 and AtHB3,13,20,23 emerged as negative RH growth regulators at low temperature (**Figures S5-S9** and **Figures 3A-F**). A higher order of mutants for multiple HBs as the quadruple mutant *athb3 (hat7) athb13 athb20 athb23* showed an enhanced RH long phenotype at low temperature while single mutants *athb13* and *athb23* alone did not. In concordance with that, two triple mutant combinations (*athb3 (hat7) athb13 athb23* and *athb13 athb20 athb23*) also showed no differences with Wt Col-0 (**Figure S7**). This result suggests a high level of functional redundancy of these four AtHBs in RH growth at low temperature. Surprisingly, *athb16* has a very short RH phenotype at low temperature. This may indicate at least two contrasting roles for these HBs (AtHB16 vs AtHB3,13,20,23) in RH growth at low temperature. In addition, we tested if most of these mutants also have defects on RH cell elongation under a low nutrient environment (MS 0.1X) at 22°C condition (**Figure S10**). In a similar trend to 10°C RH growth response, *rsl2*, *rsl4* and the double *rsl2rsl4* showed reduced cell elongation as well as *hb16-2*. On the contrary, the quadruple AtHB mutant, *athb3 (hat7) athb13 athb20 athb23* and 35S:AtHB13 behaved as Wt Col-0 (**Figure S10**). This suggests that 10°C condition reduce nutrients availabilities in similar but not exactly in the same manner as low nutrient media (MS 0.1X) condition since in the MS 0.1X nutrient reduction is proportional for all macro and micronutrients while at 10°C the effect depends on the intrinsic mobility over the temperature of each nutrient. To support the role of each TF in the primary network on RH growth, we have quantified the expression levels of the gene regulatory primary network’s nodes (*RSL4, AtHB13, GTL1/DF1* and *MYB-like*) and on specific genes of the secondary network (*AtHB16/AtHB23* and *ERD7/ERD10*) at early temporal points (0h (Basal, B), 2h, and 6h at 10°C) in selected mutant lines previously isolated and characterized (**Figure S11-S14** and **Figure 3G**). In this manner, we are able to measure the impact and influence of each of the main regulatory nodes identified in our network over the expression of the selected gene from the proposed GRN. We found that RSL4-GTL1 and GTL1-AtHB13 TFs form positive transcriptional feedback loops while RSL4-AtHB13 forms a negative feedback loop. Regarding the secondary network, GTL1 and MYB-like both enhance ERD7/ERD10 expression while RSL4 and AtHB13 both repressed AtHB16 (**Figure S13-S14**) (**Figure 3G**). This highlights a complex transcriptional landscape behind a coherent phenotypic response, in this case at low temperature even for single plant cells.

To determine whether RSL4 controls the expression of these main gene regulatory network nodes by direct binding to their promoter regions, we searched for putative RSL4-binding sites (Hwang *et al*., 2017) in open chromatin regions, according to publicly available ATAC-seq datasets (Maher *et al*., 2018). According to ChIP-qPCR using RSL4:RSL4-GFP plants and anti-GFP antibody, GTL1, AtHB13 and MYB-like are direct RSL4 targets in the primary gene network and AtHB16/AtHB23-ERD10 are their targets in the secondary network (**Figure 4A**), as revealed in comparison to the previously identified direct target EXPANSIN 7 (EXP7) (Hwang *et al*., 2017). Overall, these results confirm that RSL4 positively controls the expression of GTL1 and represses the expression of AtHB13 thus impacting RH growth. It was also tested that presence of the GTL1 binding sites in the selected gene network by ChIP-seq and positive peaks indicate direct regulation close to the starting transcriptional sites for AtHB16 and ERD7/ERD10 only at room temperature suggestion that the low temperature repressed its activity in these genes (**Figure 4B**). Based on the mutant phenotypic characterization linked to the mutants impact on selected gene expression (**Figure 3**) together with the direct regulation of RSL4-GTL1 on selected gene targets (**Figure 4A-B**), we have uncovered that two antagonistic GRNs, composed by RHD6-RSL2/RSL4-HB16 (GRN1) and by GTL1/DF1-multiple AtHBs-MYB-like (GRN2), both control RH growth at low temperature (**Figure 4C**). Then, we measured the expression levels using protein translational reporters for RSL4:RSL4-GFP (and the related RHD6:RHD6-GFP), as well as the node GTL1:GTL1-GFP (and the related DF1:DF1-GFP) (**Figure 5A-D**) both at room and at low temperature. All of them except RSL4 were induced at low temperature in epidermal cells close to the RH development and also in the elongation zone. RSL4 did not change over the treatment. In addition, GUS staining of AtHB13, AtHB16 and AtHB23 driven lines were also characterized (**Figure 5E**) and a strong signal was detected for AtHB16 in most of the root tissues including RHs, and to a lower extent, in AtHB23 also in RHs in both low temperature and room temperature. This result confirms that all the TFs identified and tested (unless AtHB13) are expressed in RHs at both room and low temperatures. It is plausible that the level of expression of AtHB13 is too low to be detected by the GUS staining or the promoter region that drives this line is missing some RH specific sequences.

**Figure 4.**
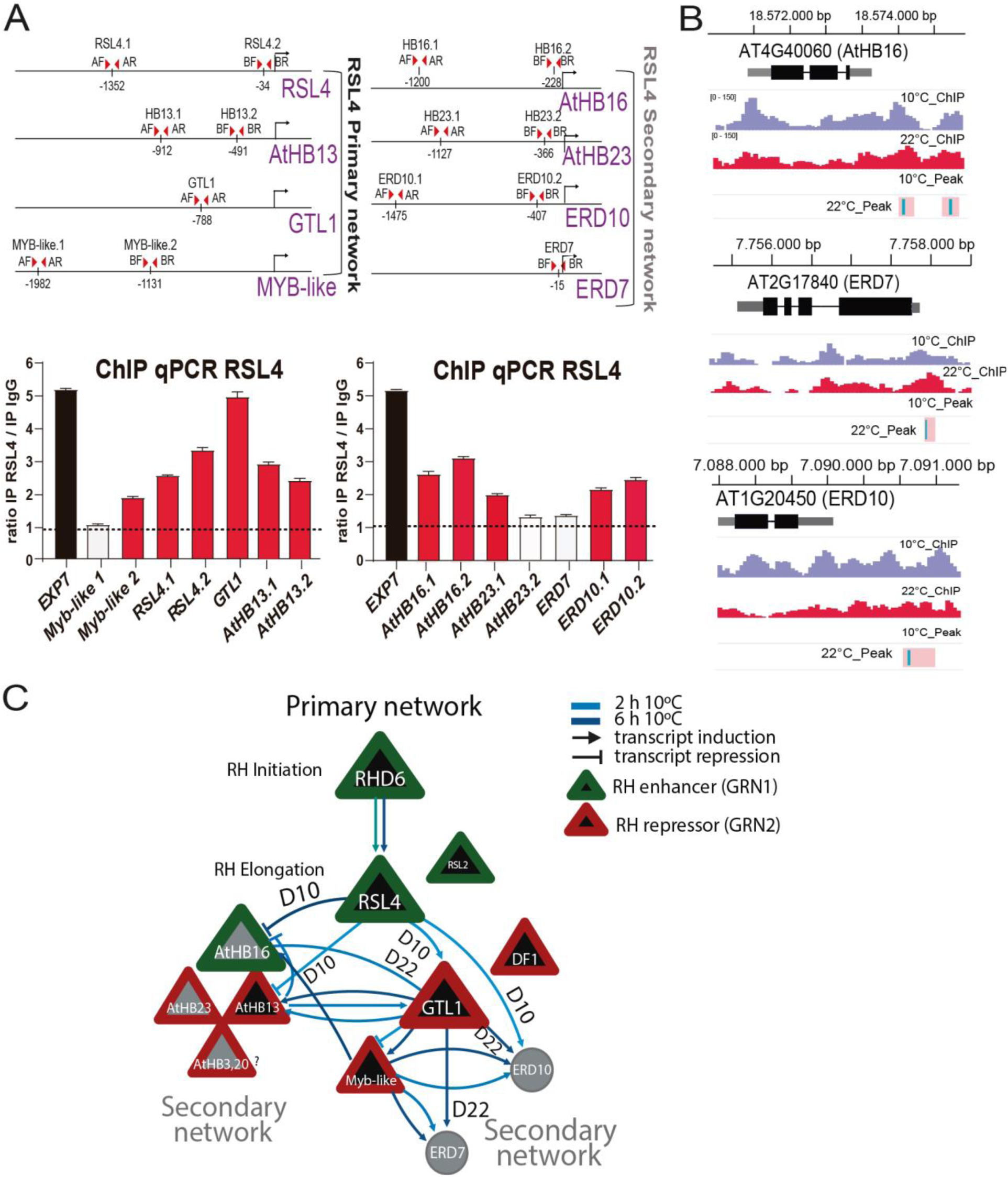
Direct regulation of RSL4 and GTL1 on selected members of the transcriptional primary and secondary network. (**A**) ChIP-qPCR analysis of RSL4 binding to the promoter regions of the gene regulatory networks at low temperature. Positive hits are indicated with red bars. Schemes of the loci showing the location of the fragments analyzed by ChIP-qPCR are shown in the upper part (A = Primer A; B = Primer B; F = Primer forward; R = Primer reverse). Primers were designed analyzing ATAC-seq experiments in regions where the chromatin is accessible (Maher *et al*., 2018). *EXP7* oligos were used as a positive control. The data are mean ± SD of 2 biological replicates. The experiment was performed twice with similar results. The enrichment was measured relative to the negative control IgG. (**B**) ChIP-seq analysis of GTL1 binding on HB16, ERD7/ERD10 as secondary RSL4 nodes gene targets promoter regions only at room temperature. Schemes of the three loci were included. ChIP-peaks at 22°C are indicated in pink-blue boxes. The data are representative of 2 biological replicates. (**C**) Proposed model of two antagonic Gene Regulatory Networks GRN1 and GRN2 acting at early times on RH growth at low temperature. Main transcriptional nodes of gene network with the detailed effect of each component at transcriptional and RH phenotypic levels. Direct regulation at 10°C/22°C (D10/D22) was assessed by ChIP results. Arrows indicate positive gene expression regulation and blunt arrows indicate transcriptional repression. (?) Indicates that needs to be experimentally validated.

**Figure 5.**
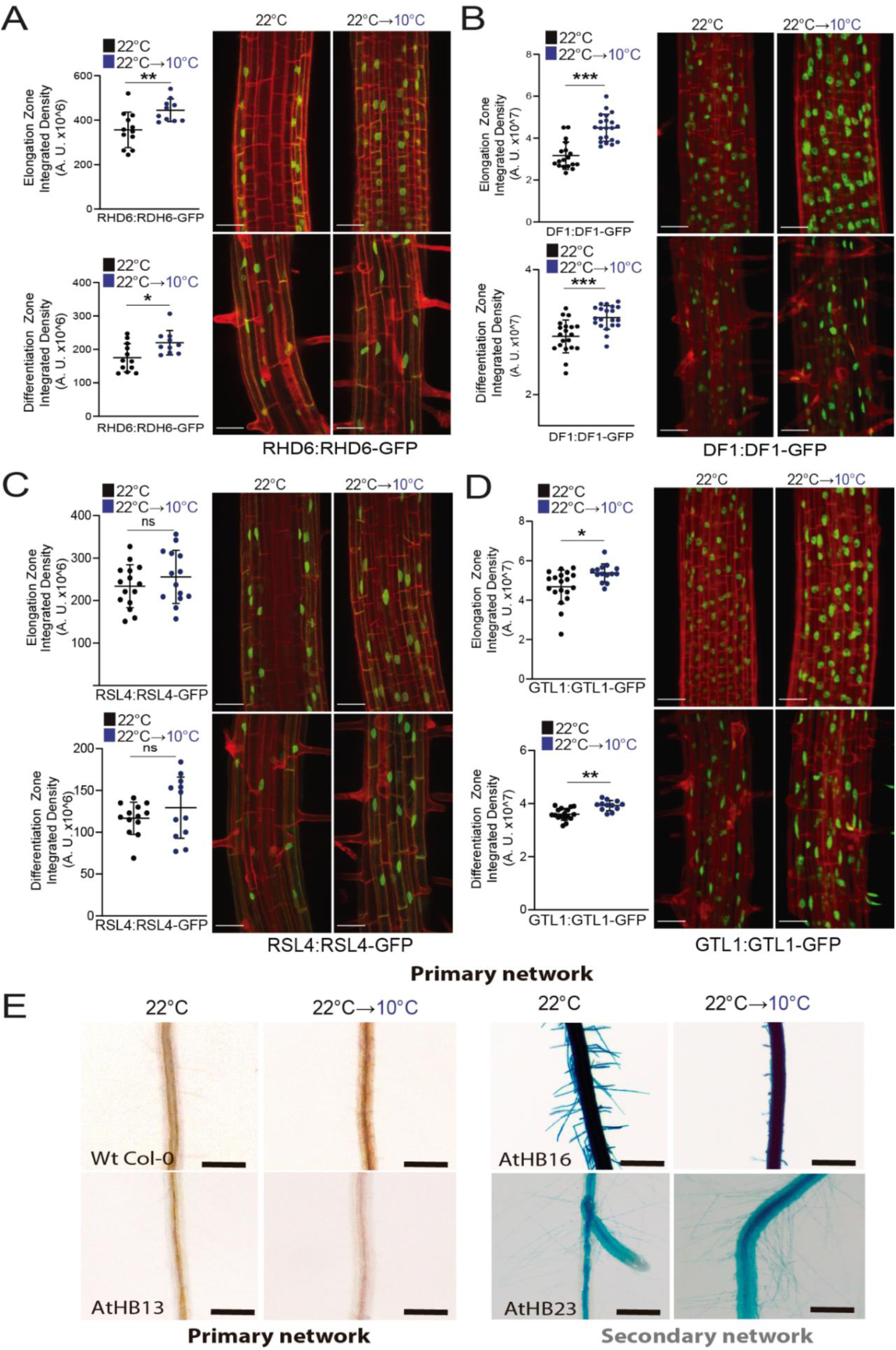
Low temperature expression of the main TF in the elongation and differentiation zones of the root. (**A**) Confocal images of RHD6:RHD6-GFP translational reporter line of at 22°C and after transfer from ambient to low temperature (22°C→10°C). Quantitative evaluation of the GFP fluorescence intensity across the root elongation zones at 22°C and after transfer from ambient to low temperature. Fluorescence intensity is expressed in arbitrary units (A.U.), N=8 roots. Results are representative of three independent experiments. Scale bars=50 μm. (**B**) Confocal images of DF1:DF1-GFP in *df1-1* translational reporter line of at 22°C and after transfer from ambient to low temperature (22°C→10°C). (**C**) Confocal images of RSL4:RSL4-GFP translational reporter line of at 22°C and after transfer from ambient to low temperature (22°C→10°C). Scale bars=50 μm. (**D**) Confocal images of GTL1:GTL1-GFP in *gtl-1* translational reporter line of at 22°C and after transfer from ambient to low temperature (22°C→10°C). Results are representative of three independent experiments. Scale bars=50 μm. (**E**) Expression in roots of GUS reporter lines for AtHB13, AtHB16 and AtHB23 at 22°C and after transfer from ambient to low temperature (22°C→10°C). Images are representative of three independent experiments. Scale bars=200 μm. All lines except AtHB23 were incubated 12hs and AtHB23 were incubated 3 days.

To further elucidate potential interactions between these two GRN identified here, we performed protein-protein interaction predictions via AlphaFold-Multimer (AFM). This approach allowed us for the prediction of protein complex structures (Evans et al., 2021). By analyzing predicted aligned error (PAE) data from AFM predictions, we calculated Local Interaction Score (LIS) and Local Interaction Area (LIA) to identify putative interacting pairs (Kim et al., 2024). The AFM analysis predicted several potential protein complexes, including one possibly composed by RHD6-RSL4 and another predicted complex included several AtHBs, such as interactions by pairs homodimers and heterodimers of AtHB3, AtHB13, AtHB16, and AtHB20 (**Figure 6A-B and Figure S15**). Interestingly, AtHB23 seems to have weaker interactions with others. Full prediction results for RSL4 regulatory network are detailed in **Supplementary Table S8**. These prediction results highlight the importance of RHD6-RSL4 as well as these 4 AtHBs in the regulation of RH growth response quantified before. Finally, we confirmed the protein-protein interactions for several of the top candidates predicted pairs in AFM (**Figure S15**) by using Bimolecular Fluorescence Complementation (BiFC) assay such as RHD6-RHD6, RSL4-RHD6, RSL4-AtHB13, RSL4-AtHB16 and AtHB16-AtHB20 in both possible orientations N-terminal GFP/C-terminal GFP (**Figure 6C**). On the contrary, we could not detect interactions between MYB-AtHB13, MYB-AtHB16, and MYB-AtHB20 that have much lower statistical probabilities to interact in the AFM prediction. This highlights the high predictive power of the *in silico* AFM approach. In sum, our systems biology approach identified new components and relevant TF-TF interactions, TF–target gene interactions that together control RH growth in responses to moderate cold. Our results uncovered that existence of two antagonistic GRNs, composed by RHD6-RSL2/RSL4-HB16 (GRN1) and by GTL1/DF1-multiple AtHBs-MYB-like (GRN2), both control RH growth at low temperature (**Figure 7**). Overall, we were able to experimentally confirm the function of these new regulatory components in low temperature-controlled RH growth in Arabidopsis, establishing a complex regulatory circuitry that integrates an environmental signal with RH development.

**Figure 6.**
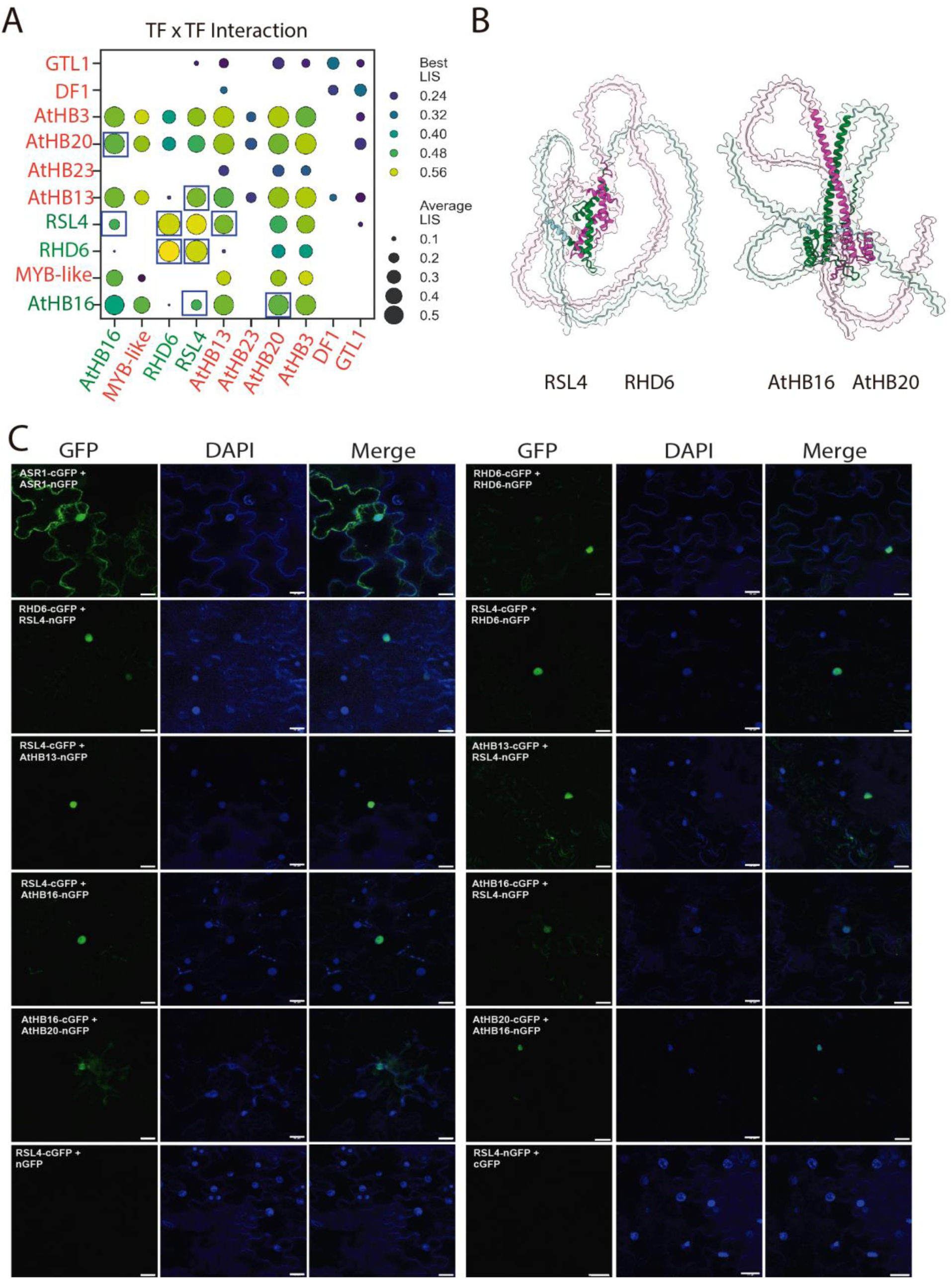
Protein-protein interactions in the regulatory networks that control RH growth at low temperature. (**A**) Dot-plot illustrating the interactions of the main RSL4-transcriptional network using the AlphaFold-Multimer (AFM) screening approach. In green are indicated those TFs that are positive regulators of RH growth at low temperature (GRN1) while in red are those TF that are negative regulators of RH growth at low temperature (GRN2). Positive interactions exceeding thresholds for best Local Interaction Score (LIS) and Local Interaction Area (LIA) and average LIS/LIA are displayed. Strongest interactions are highlighted as black boxes. In blue boxes are the pair-interactions confirmed by BiFC shown in (C). See also **Supplementary Table S8** for details. (**B**) Visualization of predicted domain structures for two selected interactions, RSL4/RHD6 and AtHB16/AtHB20. Interaction interface is highlighted by green (for RHD6 and AtHB16) and magenta (for RSL4 and AtHB20). (**C**) BiFC protein-protein interactions assessed on selected members of the GRN1 (RHD6, RSL4, and AtHB16) and GRN2 (AtHB13 and AtHB20). Positive control of self-interactive ASR1 (Ricardi et al. 2014) and negative controls (RSL4 + empty n/cGFP) were included. Scale bars=20 μm.

**Figure 7.**
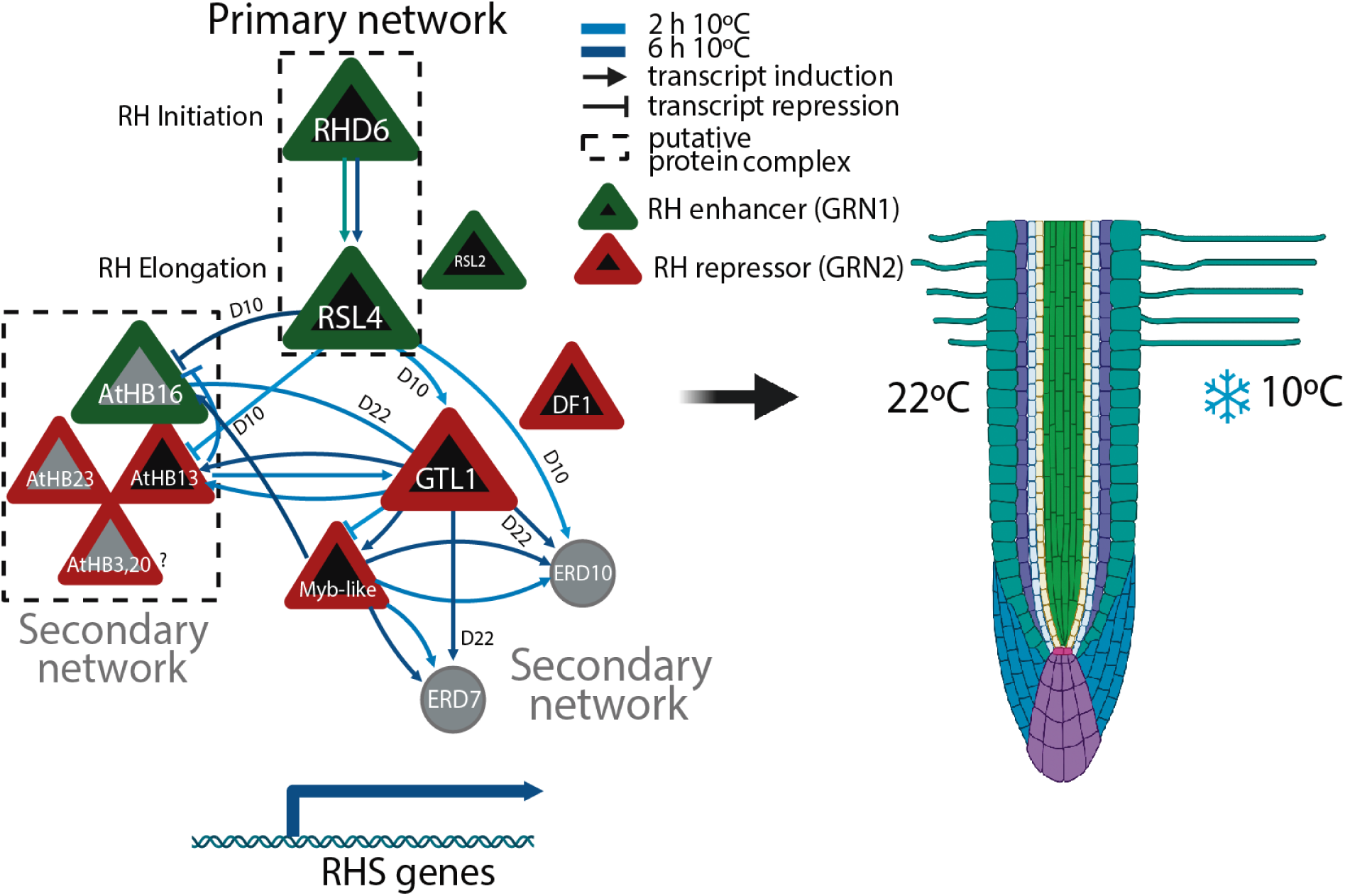
Proposed model of two antagonic GRN1 and GRN2 acting at early times on RH growth at low temperature. Main transcriptional nodes of gene network with the detailed effect of each component at transcriptional and RH phenotypic levels. Direct regulation at 10°C/22°C (D10/D22) was assessed by ChIP results. Arrows indicate positive gene expression regulation and blunt arrows indicate transcriptional repression. Protein-protein interactions proposal is based on Alpha-Fold Multimer (AFM) predictions and Bimolecular Fluorescence Complementation (BiFC) approaches. (?) Indicates that needs to be experimentally validated. RHS= Root Hair Specific genes.

## Discussion

In this work, we have extended the RH growth responses at low temperature to three monocots including Brachypodium, rice and wheat in addition to *Arabidopsis thaliana* suggesting a more general growth response than anticipated. We uncovered a regulatory map of RH growth under low temperature in Arabidopsis by focusing on early time points (hours) using an RNA-seq filtered by DAP-seq and validated expression profiling using mutant lines, coupled to ChIP approaches. This research has identified two opposing gene regulatory networks (GRN1 and GRN2) that govern the early transcriptome responses to low temperature (**Figure 7**). One gene regulatory network (GNR1) promotes the formation of root hairs. This network is controlled by certain transcription factors (FTs), including ROOT HAIR DEFECTIVE 6 (RHD6), HAIR DEFECTIVE 6-LIKE 2 and 4 (RSL2-RSL4), and a member of the homeodomain leucine zipper (HD-Zip I) group I 16 (AtHB16). Additionally, a second gene regulatory network (GRN2) was discovered to inhibit the growth of RH at low temperatures. This GRN consists of the trihelix transcription factor GT2-LIKE1 (GTL1), the previously unknown MYB-like transcription factor DF1 (AT2G01060), and several members of the HD-Zip I group (AtHB3, AtHB13, AtHB20, AtHB23). The study of both gene regulatory networks (GRNs) reveals an intricate control of root hair (RH) development in response to low temperature. None of the TFs were previously associated with low temperature (except AtHB13 and ERD7) and related to RH growth highlighting the predictive power of this approach. Previously, based on single-cell RNA sequencing, it was identified several TFs similar to our gene regulatory network (GRN) such as AtHB3 (HAT7) and GTL1 as responsive to brassinosteroids in the cortex during cell elongation (Nolan et al. 2023). This suggests that similar GRN might be sub-functionalized in different cellular contexts and under different environmental and hormonal cues. One of the negative regulators of RH growth identified here is the trihelix TF GT2-LIKE1 (GTL1) and its closest homolog, DF1, which halts RH cell growth by directly repressing RSL4 but also bind to RHD6 preventing the activation of RSL4 (Shibata *et al*., 2022) under excess nutrient conditions (Shibata *et al*., 2018; Shibata *et al*. 2022) but it was not linked to low temperature conditions. In our work, we found that GTL1 is positively controlled by RSL4 and *vice versa*, which negatively regulates RH growth at low temperature with a similar function as previously described for nutrient responses in growth media.

Several members of the Arabidopsis HD-Zip I subfamily, AtHB3, AtHB13, AtHB16, AtHB20, and AtHB23, were identified in this study as relevant TFs in the regulation of RH growth at moderate low temperature (**Figure 7**). They belong to clade V (Arce *et al*., 2011). They have a leucine zipper (Zip) downstream the homeodomain (HD), located in the middle of the protein (Viola *et al*., 2016). Arabidopsis HD-Zip I members were classified based on their exon-intron structure (Henriksson *et al*., 2005) and subsequently based on uncharacterized conserved motifs outside the HD-Zip (Arce *et al*., 2011). All these type I AtHBs identified in our study (with the exception of AtHB3) are expressed in RH cells based on single cell RNA-seq or GFP-enriched protoplast studies (Brady *et al*., 2007; Denyer *et al*., 2019; Jean-Baptiste *et al*., 2019; Ryu *et al*., 2019; Shulse *et al*., 2019; Zhang *et al*., 2019; **Supplementary Table S1**). HD-Zip I proteins are only found in plants and have been linked to environmental stresses (Perotti *et al*., 2017) but also to many developmental processes (Ribone *et al*., 2015; Capella *et al*., 2015; Perotti *et al*., 2021; Perotti et al. 2022). AtHB13 provides enhanced tolerance to both biotic and abiotic stresses, plays a role in pollen hydration and seed germination (Hanson *et al*., 2002; Cabello *et al*., 2012; Gao et al. 2014; Silva *et al*., 2016) while AtHB23 participates in the elongation of the hypocotyl and the expansion of cotyledons (Choi *et al*., 2014), with functions in lateral root development (Perotti *et al*., 2019; Perotti *et al*., 2020; Perotti et al. 2021), together with PHL1 promotes carbohydrate transport from pedicel-silique nodes to seeds (Spies *et al*., 2022), and is a developmental modulator of root growth in normal and salinity conditions, by interacting with PHL1 and MYB68 (Spies et al., 2023). Here, we found that AtHB13 is negatively regulated by RSL4-GTL1 and positively regulated by MYB-like at low temperature and participates in RH growth. It is plausible that AtHB13 do not act alone since in the RSL4-regulated RH network at low temperature were also identified several more HD-Zip I proteins. It remains to be tested if the contrasting growth roles of AtHB16 and AtHB3, AtHB13, AtHB20, AtHB23 in the RH response to low temperature are mediated via formation of protein complexes and/or by individual TFs. By AFM approach, we found that these 4 AtHBs form putative protein-protein complexes at least as homodimers and heterodimers to orchestrate the control of RH growth at low temperature (**Figure 6A-B**). In addition, by BiFC we validate the most important interactions in both of the GRN1 and GRN2. This is consistent with previous protein-proteins evidences for the AtHBs type-I TFs that are able to form homo- and heterodimers through LZ domain, interactions that are crucial for their DNA-binding capability (Palena et al. 2021; Li et al. 2022). For instance, AtHB13 shows interaction with AtHB20 among 27 other partners (Wanamaker et al. 2017) and AtHB16 interacts with AtHB5 (Johannesson et al. 2001). AtHB20 shows *in vitro* interactions with 49 other proteins, including AtHB13, AtHB21, AtHB30 (Wanamaker et al. 2017) and AtHB23 appears to interact with various other HD-ZIP proteins (Tan & Irish 2006).

In addition, we uncovered one uncharacterized MYB-like TF as part of the GRN that controls RH growth at low temperature. The MYB family, one of the biggest TF groups in seed plants, consists of 197 members and this group is characterized by the presence of a DNA-binding domain called the MYB DNA-binding domain (Riechmann *et al*., 2000) with a minimum of four repetitions of around 52 amino acids, which assemble into a helix-turn-helix shape. The MYB-CC family consists of 15 members, which are distinguished by the presence of both the MYB domain and a coiled-coil (CC) domain (Rubio *et al*., 2001). Here we identified a MYB-like TF (AT2G01060), currently not well characterized, that regulates RH growth at low temperature. Finally, downstream of RSL4-GTL1 regulators, we have identified ERD7 and ERD10 proteins that belong to a small family of highly conserved proteins that contain a PAM2 motif that may interact with poly(A)-binding protein (Aalto et al., 2012). ERD7 is strongly upregulated by both biotic and abiotic stresses such as cold, salt, excess light and Pi starvation (Cheng et al., 2013; Rasmussen et al., 2013) and, it was found to be associated with lipids droplets via its C-terminal senescence domain (SD) (Doner et al. 2021). Specifically, it binds to negatively charged phospholipids, such as phosphatidylinositol (PI) and phosphatidic acid (PA) that could induce structural and mechanical changes in the membrane that affect membrane fluidity (Barajas-Lopez et al. 2021). In particular and in agreement with our study, ERD7 is associated with low temperature stress. Overall, although our low temperature condition is moderate, ERD7 and, possibly ERD10, might play a protective role in the membrane stability during the RH cell expansion process.

Through our investigation of gene-expression patterns, we were able to uncover potential TFs that potentially control RH growth responses to low temperature. The incorporation of RH specific genes coupled to an early temporal data has a significant favorable effect on the accuracy of our network predictions. Furthermore, our findings served to identify novel TFs participating in the low temperature response triggering RH growth. This highlights the significance of using regulatory networks when investigating specific biological inquiries. Our study enabled us to ascertain that genes sensitive to low temperature and their associated biological processes are regulated in a coordinated manner in both spatial and temporal dimensions. Together with RHD6-RSL4, the TFs AtHB3, ATHB13, ATHB20, AtHB23, AtHB16 and GTL1/DF1, have significant functions in controlling the low temperature response in RH growth and the downstream changes in gene expression eventually result in the overall enhancement of RH growth. These discoveries enhance our comprehension of how plants synchronize the RH growth in response to variations in temperature and nutrient availability at the cellular level. Future challenges are related to the definition of the fine tune mechanisms of all these TFs acting in real time in the control of RH genes. As a long term goal, these new RH regulators identified here could be used to generate crops with enhanced RH growth to tailored nutrient uptake to specific soil and temperature conditions.

## Materials and methods

### Plant Material and Growth Conditions

All the *Arabidopsis thaliana* lines used were in the Columbia-0 (Col-0) background. All the plant lines sussed in this study are detailed (**Table Supplementary S6**). Seeds were surface sterilized and stratified in darkness at 4°C for 3 days before being germinated on ½ strength 0,5X MS-MES (Duchefa, Netherlands) on 0.8% Plant Agar™ (Duchefa, Netherlands) on 120 x 120 mm square petri dishes (Deltalab, Spain) in a plant growth chamber in continuous light (120 μmol s^−1^ m^−2^). For the RH phenotype characterization, seeds were surface sterilized and stratified in darkness for 3 days at 4°C. Then grown on ½ strength MS agar plates, in a plant growth chamber at 22 °C in continuous light (120 μmol s^−1^ m^−2^) for 7-14 days depending on the experiment at 22°C as a pretreatment and then at 10°C (moderate-low temperature treatment) for 12, 24, 36, 48, 60 and 72 hours. Measurements were made between 7-17 days. For quantitative analysis of RH phenotype, 10 fully elongated RHs were measured (using the ImageJ software) from each root, and 10 roots (n=10) were measured in each of the three biological replicates on vertical agar plates (total n= 30). After treatment only new RH was measured. Images were captured with a Leica EZ4 HD Stereo microscope (Leica, Germany) equipped with the LAZ ez software.Results were expressed as the mean ± SD using the GraphPad Prism 8.0.1 (USA) statistical analysis software.

### RH phenotype in monocots

Plant species and varieties used include *Brachypodium distachyon* (Bd21), rice (Kitaake), and wheat (Chinese Spring). All seeds were imbibed in water for 2 hours and then surface sterilized with 0.6% hypochlorite and 1% Triton X-100 with gentle shaking for 8 minutes. Seeds were then rinsed five times with sterile water and then placed uniformly across ½ Murashige and Skoog (MS; PhytoTechnology laboratories, USA), pH 5.7, 0.3% Gelzan (PhytoTechnology laboratories, USA) plates and stored at 4°C for 3 d in the dark. For Arabidopsis, seeds rinsed with 70% ethanol and then surface sterilized with 0.6% hypochlorite and 0.05% Triton X-100 with gentle shaking for 5 minutes. Seeds were then rinsed five times with sterile water and placed uniformly across ½ Murashige and Skoog (MS; PhytoTechnology laboratories, USA), 1% sucrose, pH 5.7, 3% Gelzan (PhytoTechnology laboratories, USA) plates and stored at 4°C for 3 days in the dark. Plants were grown in a Conviron Adaptis growth chamber model CMP6010 with a 16 h : 8 h, day : night cycle at a light intensity of 100 μmol m^−2^ s^−1^ at 22°C. 10 fully elongated root hairs from the maturation zone were measured per root after 8 d at 22°C and after 5 d at 22°C + 3d at 10°C. Images were captured using a M205 FA Leica Stereo microscope. The root length data is represented as a box blot with the solid color dot representing the mean. The root hair density data are the mean ± SE (n=10–20 roots). A t-test was used to determine significant differences between the means.

### RNA-seq analysis

For the RNA-seq analysis, seedlings were grown on ½ strength MS agar plates, in a plant growth chamber at 22 °C in continuous light (120 μmol s^−1^ m^−2^) for 14 days at 22°C as a pre-treatment and then at 10°C (moderate-low temperature treatment) for 2 and 6 hours. We analyzed a dataset with 12 factor groups (tree time points: B, 2h, 6h) and four genotypes (Col-0, *rsl2*, *rsl4* and *rsl2 rsl4*) each with three biological replicates giving 36 samples in total. Total RNA was extracted from 20–30 mg of frozen root tissue. Frozen root samples were ground in liquid nitrogen and total RNAs were extracted using E.Z.N.A Total RNA Kit I (Omega Bio-tek, Georgia, USA). RNA quantity and purity were evaluated with a Qubit®2.0 fluorometer (Invitrogen™, Carlsbad, CA, USA) using a Qubit™ RNA BR assay kit. RNA integrity and concentration were assessed by capillary electrophoresis using an automated CE Fragment Analyzer™ system (Agilent Technologies, Santa Clara, CA, USA) with the RNA kit DNF-471-0500 (15nt). Total RNA-seq libraries were prepared according to the TruSeq Stranded Total RNA Kit (Illumina, San Diego, CA, USA) following the manufacturer’s instructions. Finally, the constructed libraries were sequenced using Macrogen sequencing services (Seoul, Korea) in paired end mode on a HiSeq4000 sequencer. For total RNA differential expression analysis, a quality check was performed with FASTQC software (Andrews, 2010). Then, the adapter sequences were removed, reads with a quality score less than 30 and length less than 60 nucleotides were eliminated using Flexbar (Dodt *et al*., 2012). Resulting filtered reads were aligned against *Arabidopsis thaliana* Araport 11 genome with the STAR aligner software. A total of 36 RNA libraries were sequenced, obtaining an average of 84,741,219 reads for each one, with a minimum and maximum value of 72,658,864 and 102,182,242 reads, respectively. After filtering them by quality and removing adapters, an average of 98.6% of the reads remained and after aligning them against the *Arabidopsis thaliana* reference genome, between 88.1% (69,820,848) and 96.5% (93,740,042) of total reads were correctly aligned. For each library, the feature Counts software from the Rsubread package (Liao *et al*., 2019) was applied to assign expression values to each uniquely aligned fragment. Differential gene expression analysis was performed using the Bioconductor R edgeR package (Robinson *et al*., 2010). Differentially expressed genes (DEGs) were selected with an FDR < 0.05 and a Log2 FC > |0.5|. To search for genetic functions and pathways overrepresented in the DEG lists, genetic enrichment analysis was performed using the Genetic Ontology (GO) database with the R package ClusterProfiler v4.0.5 (Yu *et al*., 2012), using the compareCluster function. The parameters used for this analysis were: lists of differentially expressed genes for each comparison in ENTREZID, enrichGO sub-function, the universe from the total of differentially expressed genes that present annotation as genetic background, Benjamini-Hochberg statistical test and a filter of FDR less than 0.05. Subsequently, the semantics filter of GO terms was performed using the simplify function of the same package using a p-value and q-value cutoff less than 0.05.

### RT-qPCR analyses

Total root RNA was extracted from these samples using the E.Z.N.A. Total RNA Kit (Omega Bio-tek, Georgia, USA) according to the product’s protocol. One microgram of total RNA was reverse transcribed using an oligo(dT)_20_ primer (Macrogen, South Korea) and M-MLV Reverse Transcriptase™ (Invitrogen, USA) according to the manufacturer’s instructions. cDNA was then diluted 3-fold before qPCR analysis. qPCR was performed on an AriaMx™ Real-time PCR System (Agilent, USA) using 10 uL of Brilliant III Ultra-Fast SYBR Green Master-Mix (Stratagene, USA), 1 μL of cDNA, and 0.25 μM of each primer to a total volume of 20 μL per reaction. The *TAFII15* (AT4g31720) gene was used as reference for normalization of gene expression levels. Statistical analysis was performed using either parametric or non-parametric T-tests, after determining the normality of the populations via the Shapiro-Wilk test. Results were plotted showing mean and standard deviation. Primer used is detailed in **Supplementary Table S7**.

### Network construction

Direct and indirect targets of RSL4 were identified using the RSL4 cis-motif reported in Hwang *et al*., 2017 and the ConnecTF database. ConnecTF was used to connect RSL4 to its indirect targets via TF2s that are themselves regulated by RSL4 according to our RNA-seq analysis. We queried the ConnecTF database for all the DAP-seq *in vitro* binding datasets. We restricted the results returned using Target Genes filter with the list of differentially expressed genes according to our ANOVA results for *rsl4* mutants (GxT genes). For this query, the TF2s were restricted using the Filter TFs option with the list of TFs regulated by RSL4. Finally, the resulting network was exported in the Network tab, combined with the direct TF-target information based on RSL4 cis-motif, uploaded, and visualized using Cytoscape.

### TF-target list enrichment

Using the Target Enrichment tab in ConnecTF, all the TF2s that are enriched for RSL4 indirect targets were analyzed and visualized. The significance of the overlap between TF–targets captured by DAP-seq in the list of RSL4 regulated targets was calculated. The P-values are calculated using Fisher’s exact test adjusted with the Bonferroni correction. The background set of genes used for the calculation, which is by default all protein-coding genes for the Arabidopsis instances of ConnecTF, were manually set by using the Background genes option in the query page.

### Chromatin immunoprecipitation (ChIP) PCR assay

Chromatin immunoprecipitation (ChIP) assays were performed on RSL4:RSL4-GFP plants (Moison *et al*. 2021) mainly as described in (Ariel *et al*., 2020). Plants were grown for 10 d in plates containing MS 0.5× medium (pH 5.7; 0.8% agar) placed vertically in a culture chamber at 22 °C and continuous light (120 μmol s^−1^ m^−2^). After 10 d, the plates were incubated at low temperature for 24hs. The expression of RSL4 was checked by qPCR. Chromatin was crosslinked with formaldehyde 1% (v/v) for 10 min at room temperature. Crosslinking was stopped by adding glycine (125 mM final concentration) and incubating for 10 min at room temperature. Crosslinked chromatin was extracted by cell resuspension, centrifugation, cell membrane lysis, and sucrose gradient as previously described (Ariel *et al*., 2020). Nuclei were resuspended in Nuclei Lysis Buffer and chromatin was sonicated using a water bath Bioruptor Pico (Diagenode; 30 s on/30 s off pulses, at high intensity for 10 cycles). Chromatin samples were incubated for 12 h at 4 °C with Protein G Dynabeads (Invitrogen) precoated with the antibodies anti-GFP (Abcam ab290) or anti-IgG (Abcam ab6702) as a negative control. Immunoprecipitated DNA was recovered using Phenol:Chloroform: Iso-amilic Acid (25:24:1; Sigma) and analyzed by qPCR using the primers listed in **Supplementary Table S7.** One upstream region of the EXP7 gene was used as a positive control (Hwang *et al*., 2017). Untreated sonicated chromatin was processed in parallel and considered the input sample. The GraphPad Prism 6 software was used to analyze the data and produce the graphs.

### GTL1 ChIP-seq analysis

ChIP-seq analysis was performed using roots of GTL1::GTL1-GFP/ *gtl1-1* mutants expressing GTL1-GFP proteins under its native promoter in the mutant background. Plants were grown in continuous light at 22°C for 12 days, then transferred to 22°C or 10°C for 6 hrs. The chromatin immunoprecipitation was carried out using antibodies against GFP (ab290, Abcam). The experiment was repeated twice with different biological samples. Illumina sequencing reads derived from ChIP and Input DNA of GTL1 were mapped to the TAIR10 reference genome employing Bowtie2 for alignment. Subsequently, the elimination of duplicate reads was performed. For the identification of GTL1 ChIP peaks in relation to its corresponding Input DNA control, the MACS2 software was utilized with a significance threshold set at a q-value of 0.05. Post peak-calling, gene annotation of the identified peaks was conducted utilizing BEDtools, which incorporated a 2 kb region upstream from the transcription start site (TSS) for consideration. The preparation of the sequencing data for visualization involved filtering, ordering, and scaling of bam files, followed by conversion to bigwig format. This conversion leveraged the “bamCoverage” function within deepTools 2.0, configured with a bin size of 10 bp, RPKM normalization, and a smooth length of 60. Visual representations of the genomic data displayed in **Figure 4B** were generated using the Integrative Genomics Viewer (IGV)(reference: 10.1093/bib/bbs017).

### GUS Activity Assay

For the GUS activity assay, a working solution for the GUS stain was prepared, containing the following components: 10 mM EDTA (pH 8), 0.1% Triton X-100, 42.3 mM Na₂HPO₄, 57.7 mM NaH₂PO₄, 0.05 mM K₃Fe(CN)₆, 0.05 mM K₄Fe(CN)₆, 1 mM X-Gluc, and Milli-Q water. The plants were grown for 5 days under continuous light at 22 °C. After this growth period, the plates were transferred to 10 °C for 3 days. Following the cold treatment, the plants were fixed in the GUS stain solution and subjected to a vacuum procedure at 65 mMPa for 15 minutes in the absence of light. After vacuum treatment, the plants were incubated at 37 °C for 12 hours and 3 days in a light-free incubator. After this incubation period, the plants were analyzed and photographed using a stereoscopic microscope (Leica).

### Confocal Microscopy

Confocal laser scanning microscopy for the lines RSL4:RSL4-GFP, RSL2:RSL2-GFP, RHD6:RHD6-GFP, GTL1:GTL1-GFP, and DF1:DF1-GFP was performed using Zeiss LSM5 Pascal (Zeiss, Germany) (Excitation: 488 nm argon laser; Emission: 490–525 nm, Zeiss Plain Apochromat 10X/0.30 or 40X/1.2 WI objectives according to experiment purpose). Z stacks were done with an optical slice of 1 µm, and fluorescence intensity was measured at the RH tip. The scanning was performed using confocal laser scanning microscopy Zeiss LSM 710 (Carl Zeiss, Germany). For image acquisition, 20x/1.0 NA Plan-Apochromat objective for root tips were used. The GFP signal was excited with a 488 nm argon laser at 4% laser power intensity and emission band of 493-549 nm. Propidium Iodide signal was excited with a 488 nm argon laser at 4% laser power intensity and emission band 519-583 nm. GFP signal at the RH tip were quantified using the ImageJ software. Fluorescence AU was expressed as the mean ± SD using the GraphPad Prism 8.0.1 (USA) statistical analysis software. Results are representative of two independent experiments, each involving 15 roots.

### Protein complex prediction using AlphaFold-Multimer

For the prediction of protein complexes using AlphaFold-Multimer (AFM), we employed LocalColabFold version 1.5.2 (Mirdita et al., 2022). It integrates AFM version 2.3.1 (Evans et al., 2021) and utilizes MMseqs2 for generating multiple sequence alignments. Computations were performed on the Harvard O2 high-performance computing cluster. Our prediction involved five models for each complex, each undergoing five recycling iterations. After AFM analysis, Local Interaction Score (LIS) and Local Interaction Area (LIA) were calculated to identify potential positive interaction (Kim et al., 2024). The code for LIS/LIA analysis is available in https://github.com/flyark/AFM-LIS. For the visualization of the predicted structures, ChimeraX was utilized (Meng et al. 2023). Results are included in **Supplementary Figure 15** and **Supplementary Table S8.** ATG number, protein name and (Uniprot ID) are indicated for each TF analyzed: AT1G66470 RHD6 (A0A178W430); AT1G27740 RSL4 (A0A384L1Z0); AT1G69780 AtHB13 (A0A654EP87); AT2G01060 MYB-like (A0A178VZU9); AT1G33240 GTL1 (Q9C882-3); AT1G76880 DF1 (Q9C6K3); AT4G40060 AtHB16 (A0A178V0P3); AT5G15150 AtHB3 (B5RID5); AT3G01220 AtHB20 (Q8LAT0); AT5G39760 AtHB23 (A0A654G6H4); AT2G17840 ERD7 (A0A178VW80); AT1G20450 ERD10 (P42759-1).

### Bimolecular Fluorescence Complementation (BiFC)

Plasmid constructs. The primers used in this study are listed in the Supplementary Data. All plasmids reported in this work were provided by Trevor Nolan. The Arabidopsis thaliana CDSs of *RHD6, RSL4, HB13, HB16, HB20 and Myb-like* were amplified with PfuUltra II Phusion High-fidelity DNA polymerase (Agilent). All gene constructs were subcloned in pCR8/GW/TOPO vector (invitrogen), verified by sequencing and cloned into BiFC binary destiny vectors pVYNE and pVYCE, (Waadt et al., 2008) via LR reaction (Invitrogen). All vectors are provided with kanamycin selection marker for E. coli and A. tumefaciens. *Agrobacterium tumefaciens* cells (strain DB31.01) carrying the different constructs were grown in 5mL of lB supplemented with gentamicin, rifampicin and kanamycin overnight. Once the cultured reach OD 600= 0.8-1.0, the cultured cells were pelleted and resuspended in 1mL of 10mM MES/KOH pH5.6, 10mM MgCl_2_ and 150 uM acetosyringone and incubated for 3 hours at room temperature. Young leaves of 3-4 week-old *Nicotiana benthamiana* plants were co-infiltrated with a 0.5 OD culture of an equal mix of both Agrobacterium tumefaciens harboring the different BiFC constructs plus OD 600 = 0.3 of the p19 helper strain. After infiltration, all plants were kept in the greenhouse until the end of the analysis. After 3 days, the same leaves were infiltrated with 10 ug/mL of water dissolved DAPI, incubated in the dark for 20 minutes and observed in a confocal microscope. Images were acquired in an Olympus IX-81 confocal microscope with a 20X objective with digital zoom. For excitation, we used 465 and 488 lasers for DAPI and ALEXA 488/GFP, respectively. Emission filters were 430-470 nm for DAPI and 505-525 nm for Alexa488/GFP. Images were obtained in a sequential mode. All images were processed with ImageJ.

## Acknowledgements

We thank NASC (Ohio State University) for providing T-DNA lines seed lines, J.M.E., F.A. and R.L.C. are investigators of the National Research Council (CONICET) from Argentina. M.I. is supported by ANID FONDECYT POSTDOCTORADO [grant 3220138]. This work was supported by grants from ANPCyT (PICT2019-0015 and PICT2021-0514), by ANID – Programa Iniciativa Científica Milenio ICN17_022, NCN2021_010 and Fondo Nacional de Desarrollo Científico y Tecnológico [1200010] to J.M.E. J.M.A lab is supported by ANID FONDECYT 1210389, Programa Iniciativa Científica Milenio ICN17_022, and NSF Plant Genome Grant NSF-PGRP: IOS-1840761. We are grateful to the Research Computing Group at Harvard Medical School for access to the O2 High Performance Compute Cluster. A.K. was supported by the Postdoctoral Fellowship Program (Nurturing Next-generation Researchers) through the National Research Foundation of Korea (NRF) funded by the Ministry of Education (2021R1A6A3A14039622). N.P. is an investigator of Howard Hughes Medical Institute.

## Author Contribution

T.U.L., V.B.G., M.A.I. and H.S-G., performed most of the experiments, analyzed the data and helped in the writing process of the manuscript. J.P. and R.A. performed some experiments and helped in the writing process. G.N-L. and C.M. performed the RNA-seq analysis, T.M. and J.M.A. performed the gene network analysis. J.M.A also helped with the writing process. L.F. and F.A. carried out the ChIP experiment of RSL4 and helped on the writing process. A.K. and N.P. performed the AlphaFold-Multimer analysis. M.R. and K.S. carried out the root hair characterization in monocots. A.A.M. helped in the writing process. M.S. and K.S. provided seeds and helped in the writing process. F.P., V.N.M., F.P.S. carried out preliminary phenotypic assays and provided several HB mutants, transgenic, promoters for AtHB genes. T.N. provided multiple HBs mutants. R.L.C. provide most of the AtHBs lines and help in the writing process. J.M.E. designed research, analyzed the data, supervised the project, and wrote the paper. All authors commented on the results and the manuscript. This manuscript has not been published and is not under consideration for publication elsewhere. All the authors have read the manuscript and have approved this submission.

