## Supplementary Files for "Two antagonistic gene regulatory networks drive Arabidopsis root hair growth at low temperature"

Figures S1-S15
Supplementary Tables 1-8

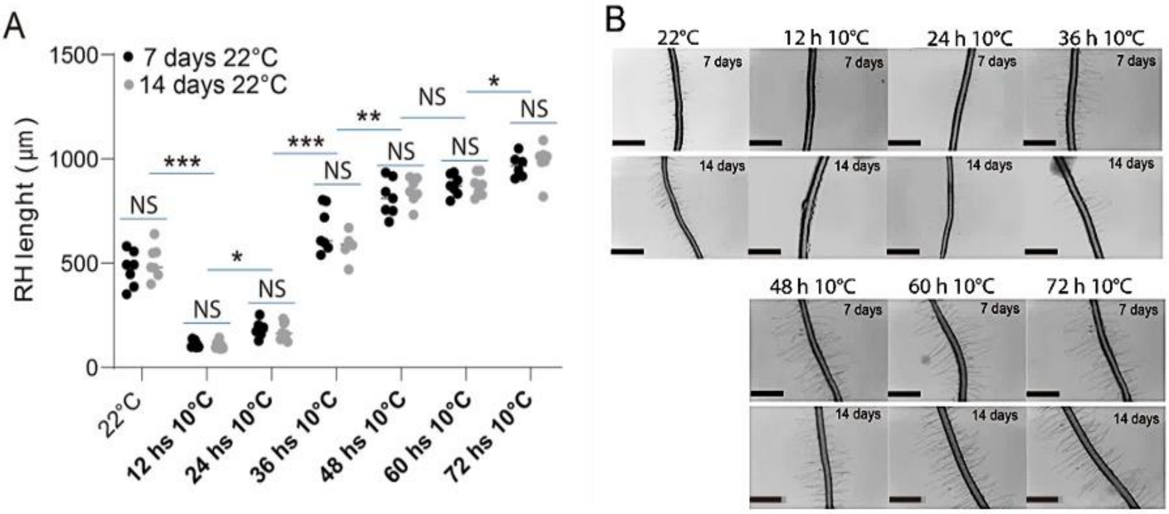


**Figure S1. Influence of room temperature pretreatment time length on low temperature RH growth in Arabidopsis thaliana**. (**A**) quantification of RH length at different treatment times at 22°C prior to cold exposure (7 and 14 days) and different cold exposure times (0, 12, 24, 36, 48, 60 and 72 hours) in both pretreatment conditions. (**B**) visualization by stereo microscopy of the phenotypic data represented. Data are the mean ± SD (N=20 roots), two-way ANOVA followed by a Tukey–Kramer test; (*) p <0.05, (**) p <0.01, (***) p <0.001, NS=non-significant. Results are representative of three independent experiments. Asterisks indicate significant differences between Col-0 and the corresponding genotype at the same temperature. Representative images of each genotype are shown below. Scale bars= 500 µm.

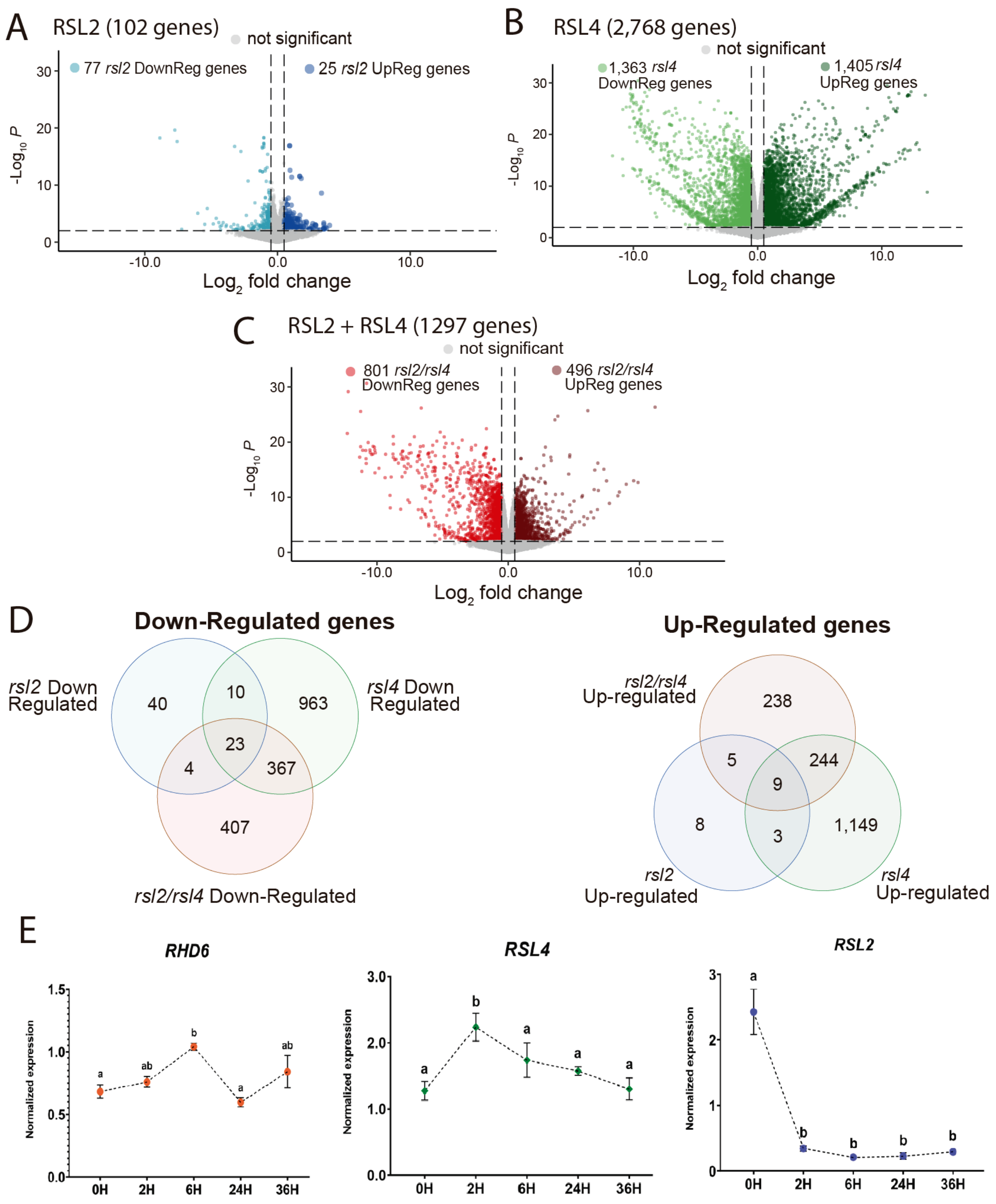


**Figure S2. Differential expression analysis between Col-0, *rsl2*, *rsl4* and rsl2 *rsl4* mutants under low temperature treatment and expression of RSLs under low temperature treatment.** (**A**) Volcano plot for differentially expressed genes comparing Col-0 and *rsl2* single mutant. Candidate genes were filtered with a *p-value* < 0.01. Down (light blue points) and up (dark blue points) regulated *rsl2* genes were selected with an FC<-0.5 and FC>0.5, respectively. (**B**) Volcano plot for differentially expressed genes comparing Col-0 and *rsl4* single mutant.

Candidate genes were filtered with a *p-value* < 0.01. Down (green points) and up (dark green points) regulated *rsl4* genes were selected with an FC<-0.5 and FC>0.5, respectively. (**C)** Volcano plot for differentially expressed genes comparing Col-0 and *rsl2rsl4* double mutant. Candidate genes were filtered with a *p-value* < 0.01. Down (red points) and up (dark red points) regulated *rsl2rsl4* genes were selected with an FC<-0.5 and FC > 0.5, respectively. (**D**) Venn diagram comparing the down (left side) and up (right side) regulated transcripts between *rsl2, rsl4* and *rsl2rsl4* candidate genes. (**E**) Expression of *RHD6*, *RSL4* and *RSL2* under low temperature. *RHD6, RSL2* and *RLS4* transcript levels were measured by RT-qPCR in roots of Wt Col-0 roots in parallel after 10 days at 22°C, followed by 6h, 24h and 36h at 10°C. Statistical differences between populations were assessed by one-way ANOVAs followed by Tukey post-tests. Significant differences were represented by letters. The average Cts for the technical duplicates of each transcript were normalized using the constant housekeeping transcript of *TAFII15*, which was evaluated parallelly in the same fashion. (a) p <0.05, (ab) p <0.005, (b) p <0.001.

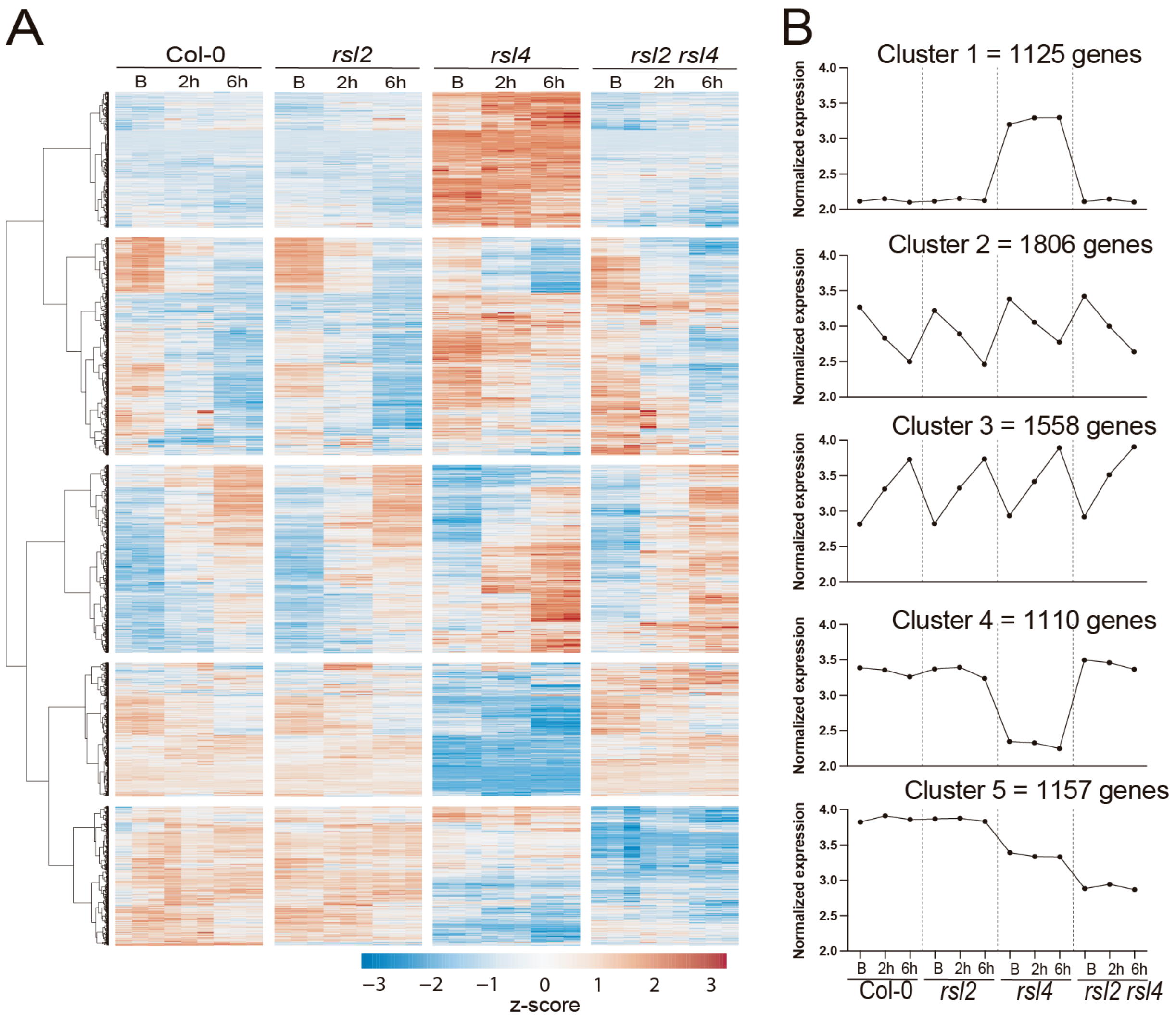


**Figure S3**. **DEG analysis between Col-0 and *rsl2, rsl4* and *rsl2 rsl4* mutants at low temperature at 2h and 6h.** (**A**) Root samples of *Arabidopsis thaliana* cv. Col-0 were compared with the mutant samples *rsl2*, *rsl4* and the double mutant *rsl2 rsl4* in a basal state at 22°C (B) and two cold treatments for 2 and 6 hours at 10°C (2h and 6h). 6,756 differentially expressed genes (DEG) in at least one condition were represented in a blue-red scale heatmap. Each column represents the expression of each replicate. Expression data was scaled considering the mean centered divided to standard deviation. (**B**) Clustering analysis of differential expressed genes was made using the Ward method. Continuous line represents each cluster tendency. See **Supplementary Table S1**.

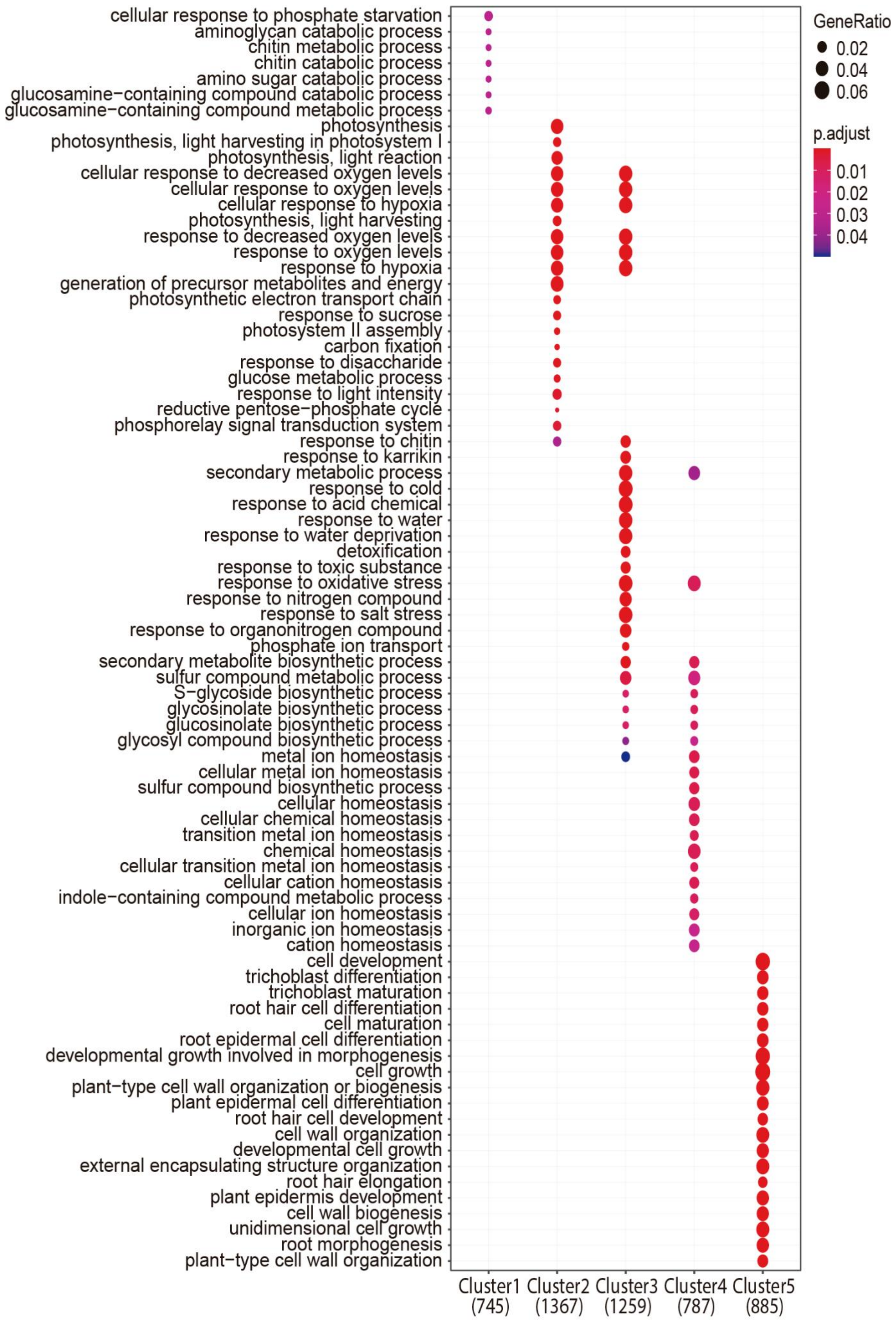


**Figure S4**. **GO analysis of the 5 clusters identified from the DEG analysis between Col-0 and *rsl2, rsl4* and *rsl2 rsl4* mutants at low temperature at 2h and 6h.** Gene ontology term enrichment analysis of genes belonging to each one of the identified 1-5 clusters. GO in cluster

5 showed several RH development processes. The blue-red scale color represents adjusted p-value and the point size represents gene ratio.

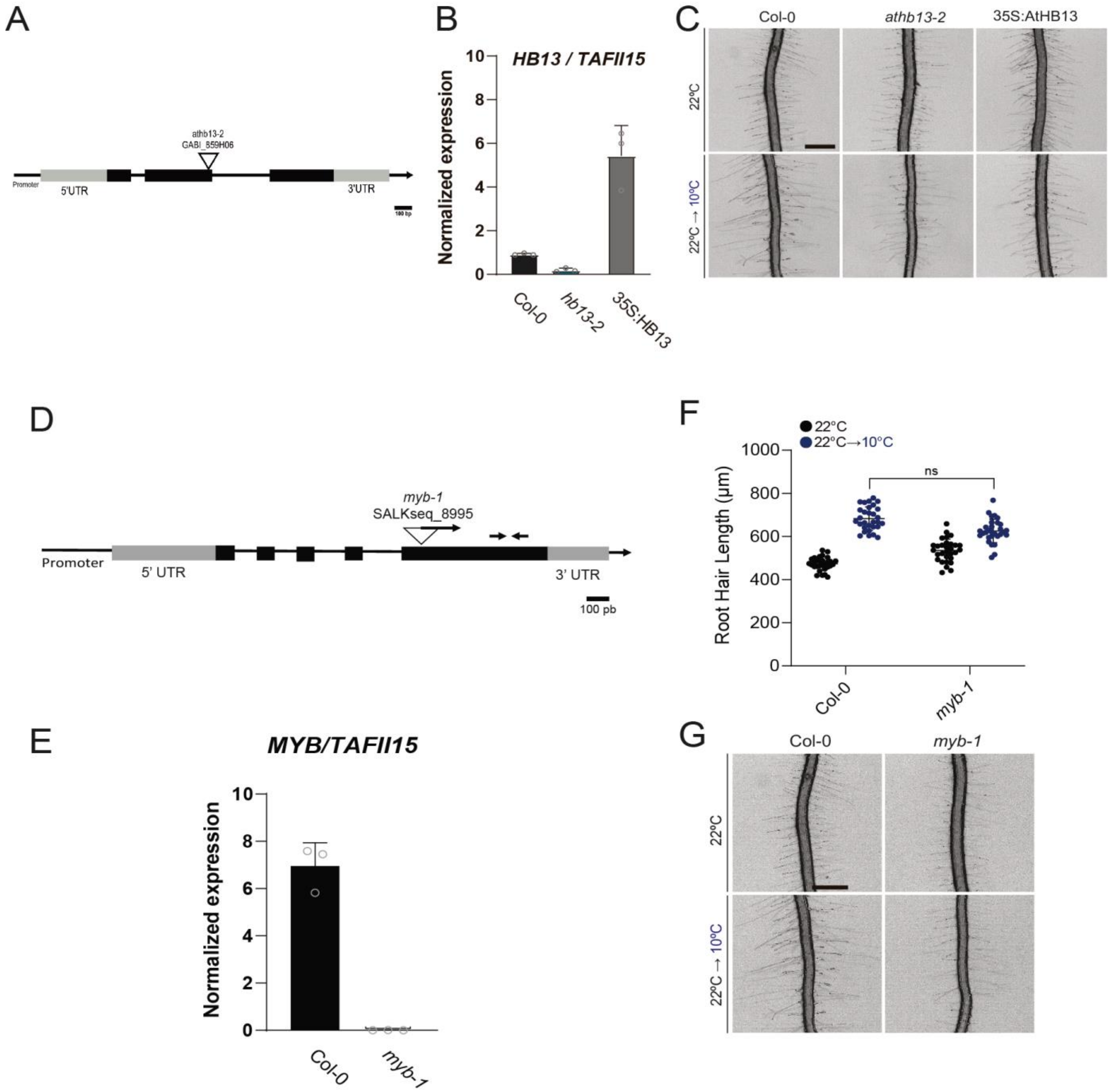


**Figure S5**. **Characterization of AtHB13 and MYB-like.** (**A**) AtHB13 gene structure and the insertion of the T-DNA mutants. Arrow indicates the position of the primers. (**B**) AtHB13 expression levels in Col-0, *athb13-2* and 35S:HB13 lines. (**C**) Representative images of each genotype are shown below. Scale bars= 500 µm. (**D**) MYB-like gene structure and the insertion of the T-DNA mutant. (**E**) MYB-like expression levels in Col-0, *myb-like-1* line. (**F**) Scatter-plot of RH length of Col-0, mutants grown at 22°C or at 10°C. Each point is the mean of the length of the 10 longest RHs identified in a single root. Data are the mean ± SD (N=20 roots), two-way ANOVA followed by a Tukey–Kramer test; ns=non-significant. Results are representative of three independent experiments. Asterisks indicate significant differences between Col-0 and the corresponding genotype at the same temperature. (**G**) Representative images of each genotype. Scale bars= 500 µm.

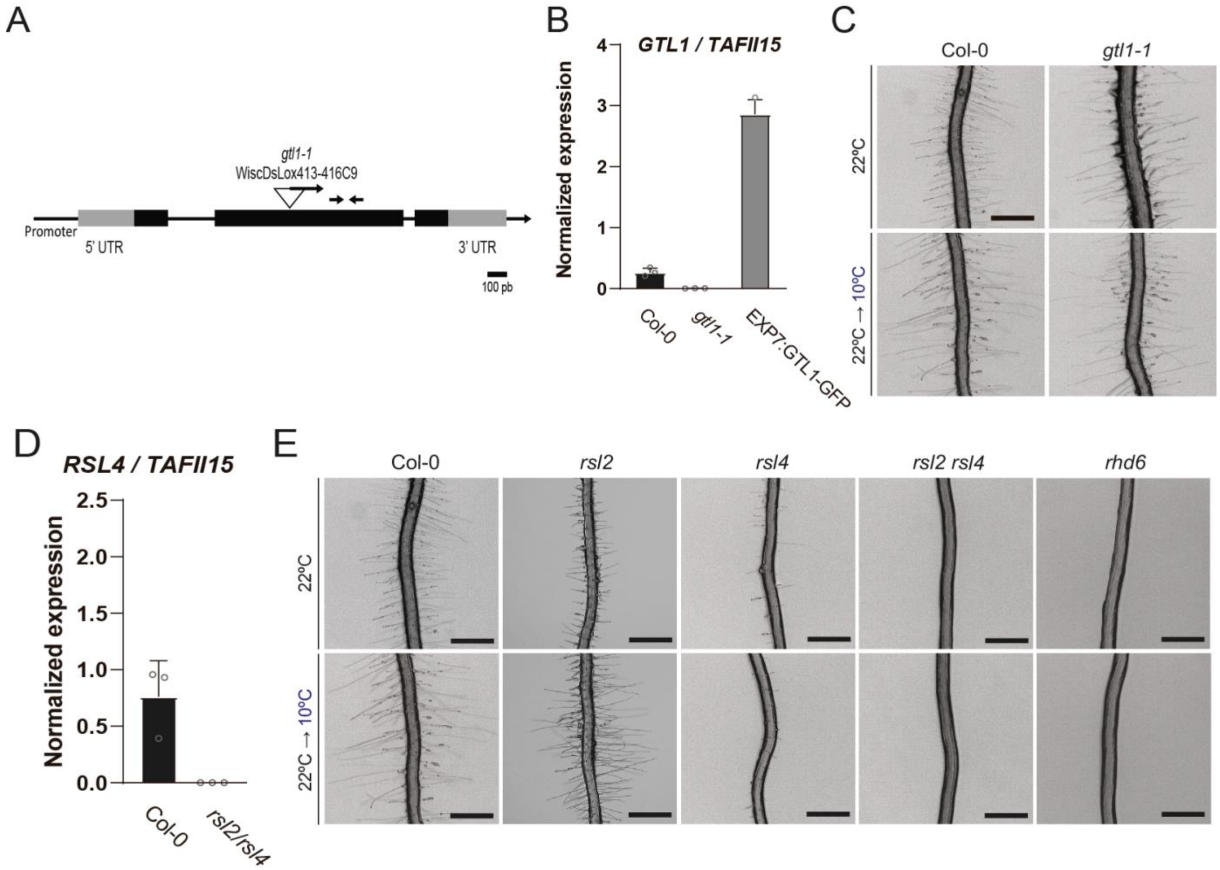


**Figure S6**. **Characterization of GTL1 and RSLs** (**A**) GTL1 gene structure and the insertion of the T-DNA mutant *gtl1-1*. (**B**) GTL1 expression levels in Col-0, *gtl1-1* and the overexpressor EXP7:GTL1-GFP lines. (**C**) Representative images of each genotype are shown below. Scale bars= 500 µm. (**D**) RSL4 expression levels in Col-0 and in the *rsl2/rsl4* line (**E**) Representative images of each genotype are shown below. Scale bars= 500 µm.

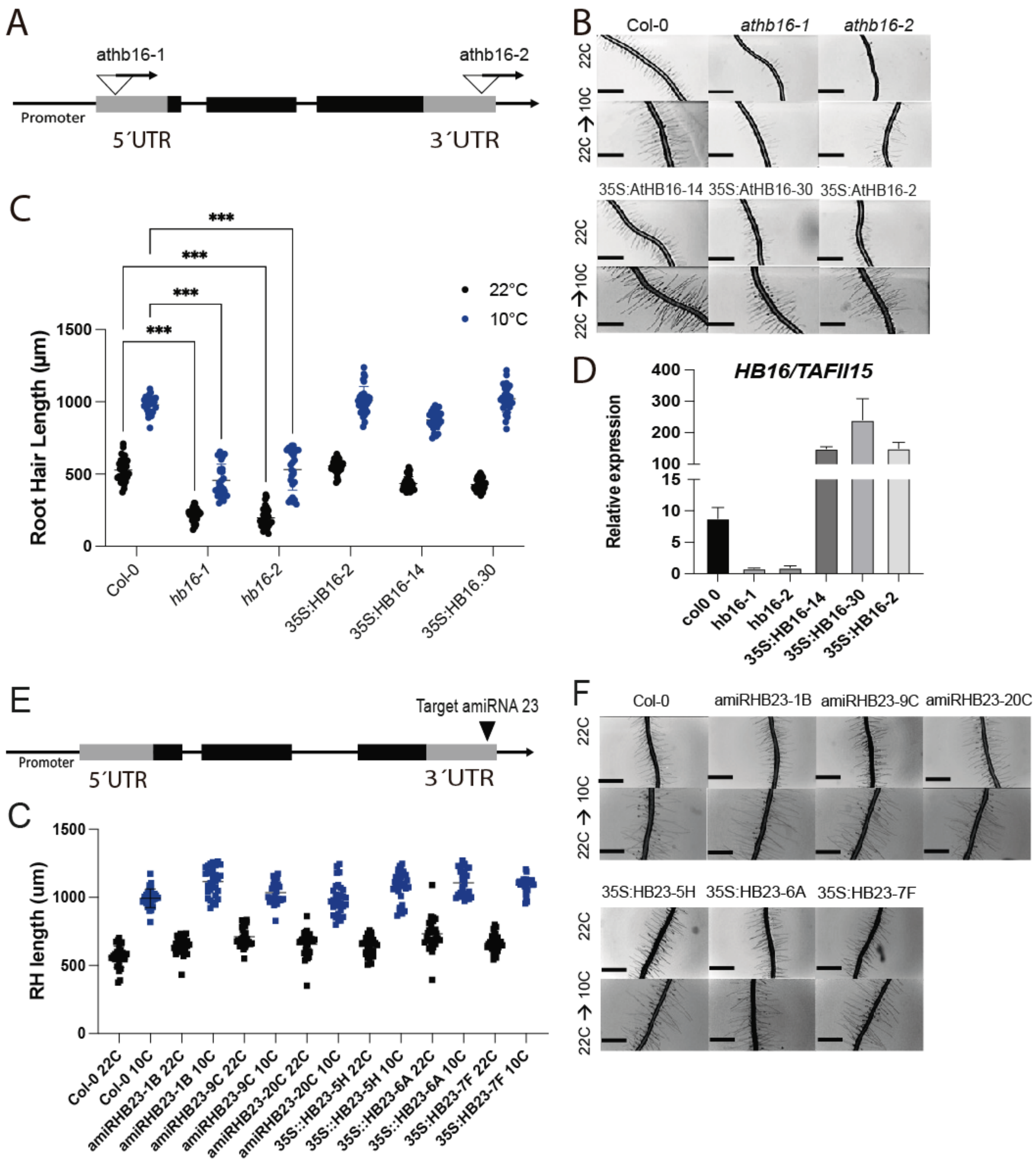


**Figure S7**. **Characterization of AtHB16 and AtHB23.** (**A**) AtHB16 gene structure and the insertion of the T-DNA mutants. (**B**) Scatter-plot of RH length of Col-0, *athb16-1* and *athb16-2* mutants and 35S:AtHB16 lines (2,14,30) grown at 22°C or at 10°C. Each point is the mean of the length of the 10 longest RHs identified in a single root. Data are the mean ± SD (N=20 roots), two-way ANOVA followed by a Tukey–Kramer test; (***) p <0.001, NS=non-significant. Results are representative of three independent experiments. Asterisks indicate significant differences between Col-0 and the corresponding genotype at the same temperature. (**C**) Representative images of each genotype are shown below. Scale bars= 500 µm. (**D**) AtHB13 expression levels in Col-0, *athb16-1* and *athb16-2* mutants and 35S:AtHB16 overexpression lines (2,14,30). **S10**. (**E**) AtHB23 gene structure and target of the amiRNA *athb23* silenced lines. (**F**) Scatter-plot of RH length of Col-0, amiRNA *athb23* (IB, 9C, 20C) lines and 35S:AtHB23 lines (5H, 6A,7F) grown at 22°C or at 10°C. Each point is the mean of the length of the 10 longest RHs identified in a single root. Data are the mean ± SD (N=20 roots), two-way ANOVA followed by a Tukey–Kramer test; (**) p <0.01, (***) p <0.001, NS=non-significant. Results are representative of three independent experiments. Asterisks indicate significant differences between Col-0 and the corresponding genotype at the same temperature. (**G**) Representative images of each genotype are shown below. Scale bars= 500 µm.

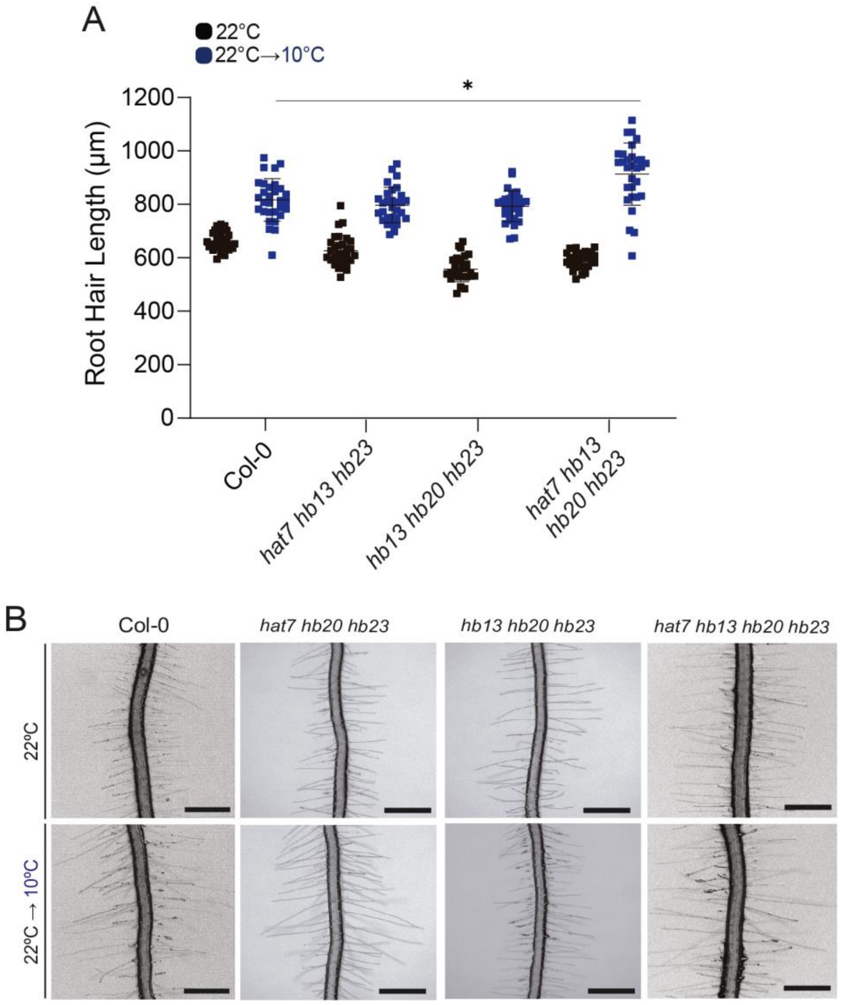


**Figure S8**. **Characterization of AtHB3, AtHB13, AtHB20, AtHB23.** (**A**) Scatter-plot of RH length of Col-0, the triple mutants *athb3 (hat7) athb13 athb23 athb13* and *athb13 athb20 athb23*, and the quadruple *athb3 (hat7) athb13 athb20 athb23* grown at 22°C or at 10°C. Each point is the mean of the length of the 10 longest RHs identified in a single root. Data are the mean ± SD (N=20 roots), two-way ANOVA followed by a Tukey–Kramer test; (*) $p < 0.05$. Results are representative of three independent experiments. Asterisks indicate significant differences between Col-0 and the corresponding genotype at the same temperature. (**B**) Representative images of each genotype are shown below. Scale bars= 500 µm.

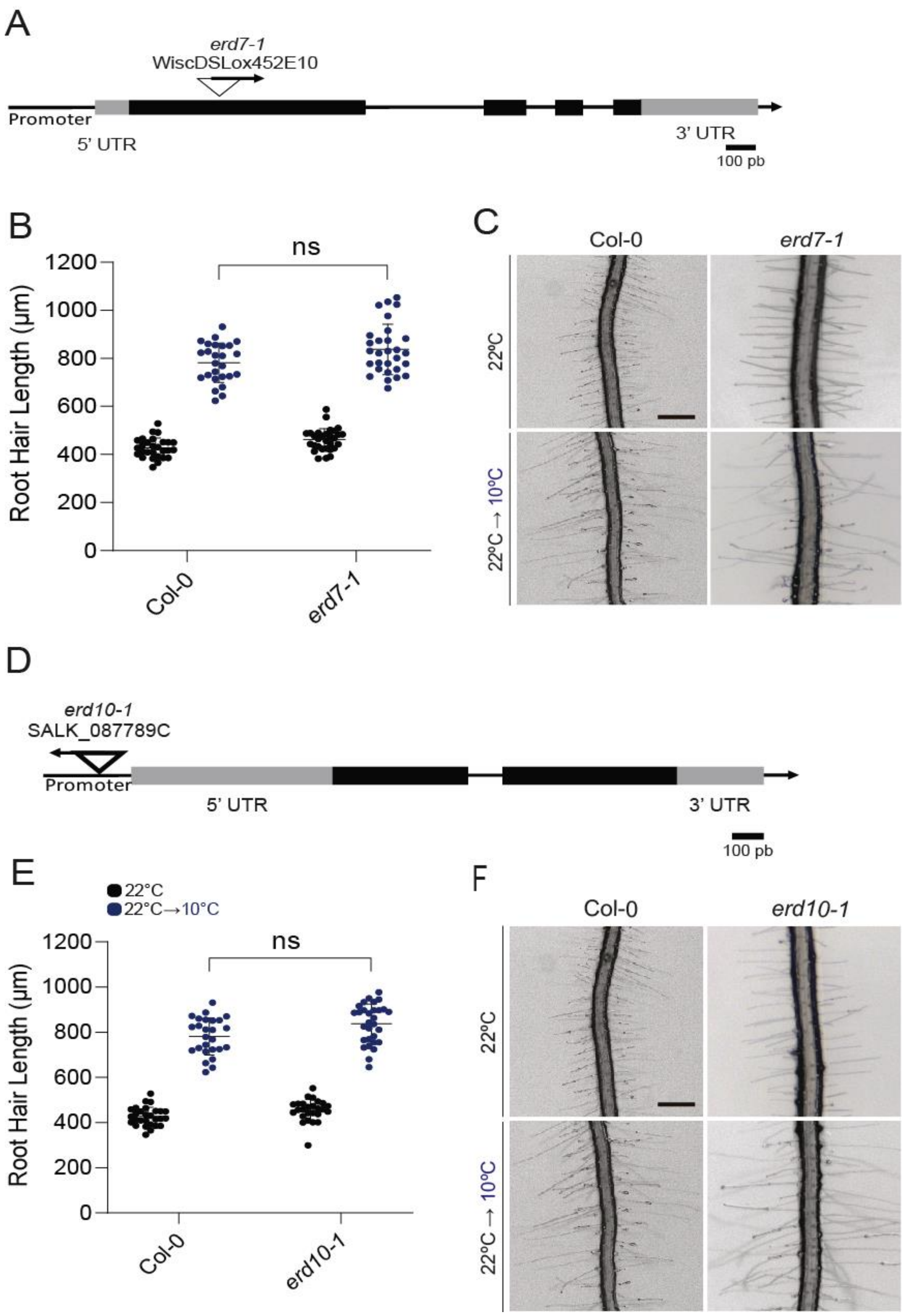


**Figure S9**. **Characterization of ERD7 and ERD10.** (**A**) ERD7 gene structure and the insertion of the T-DNA mutants. (**B**) Scatter-plot of RH length of Col-0, *erd7-1* mutant grown at 22°C or at 10°C. Each point is the mean of the length of the 10 longest RHs identified in a single root. Data are the mean ± SD (N=20 roots), two-way ANOVA followed by a Tukey–Kramer test; ns=non-significant. Results are representative of three independent experiments. Asterisks indicate significant differences between Col-0 and the corresponding genotype at the same temperature. (**C**) Representative images of each genotype are shown below. Scale bars= 500 µm. (**D**) ERD10 gene structure and the insertion of the T-DNA mutants. (**E**) Scatter-plot of RH length of Col-0 and *erd10-1* mutant grown at 22°C or at 10°C. Each point is the mean of the length of the 10 longest RHs identified in a single root. Data are the mean ± SD (N=20 roots), two-way ANOVA followed by a Tukey–Kramer test; ms=non-significant. Results are representative of three independent experiments. Asterisks indicate significant differences between Col-0 and the corresponding genotype at the same temperature. (**F**) Representative images of each genotype are shown below. Scale bars= 500 µm.

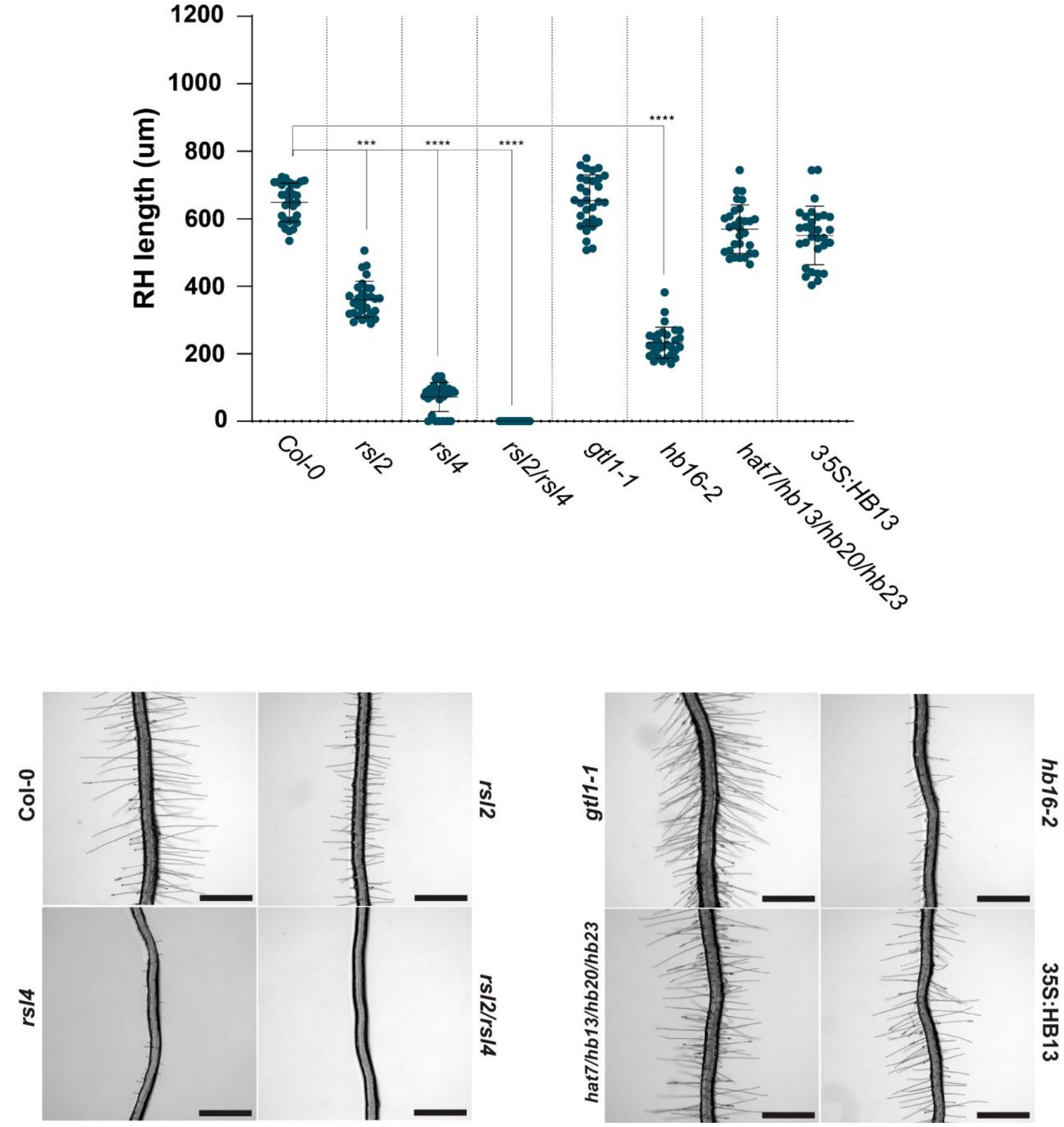


**Figure S10. Characterization of genotypes growth at low nutrient media (MS 0.1X).** Scatter-plot of RH length of Col-0, *rsl2*, *rsl4,* double mutant *rsl2 rsl4*, *athb16-2,* the quadruple mutant *athb3 (hat7) athb13 athb23 athb13* and AtHB13 grown at 22°C on MS 0.1X. Each point is the mean of the length of the 10 longest RHs identified in a single root. Data are the mean ± SD (N=20 roots), two-way ANOVA followed by a Tukey–Kramer test; ns=non-significant. Results are representative of three independent experiments. Asterisks indicate significant differences between Col-0 and the corresponding genotype at the same condition. (**B**) Representative images of each genotype are shown below. Scale bars= 500 µm.

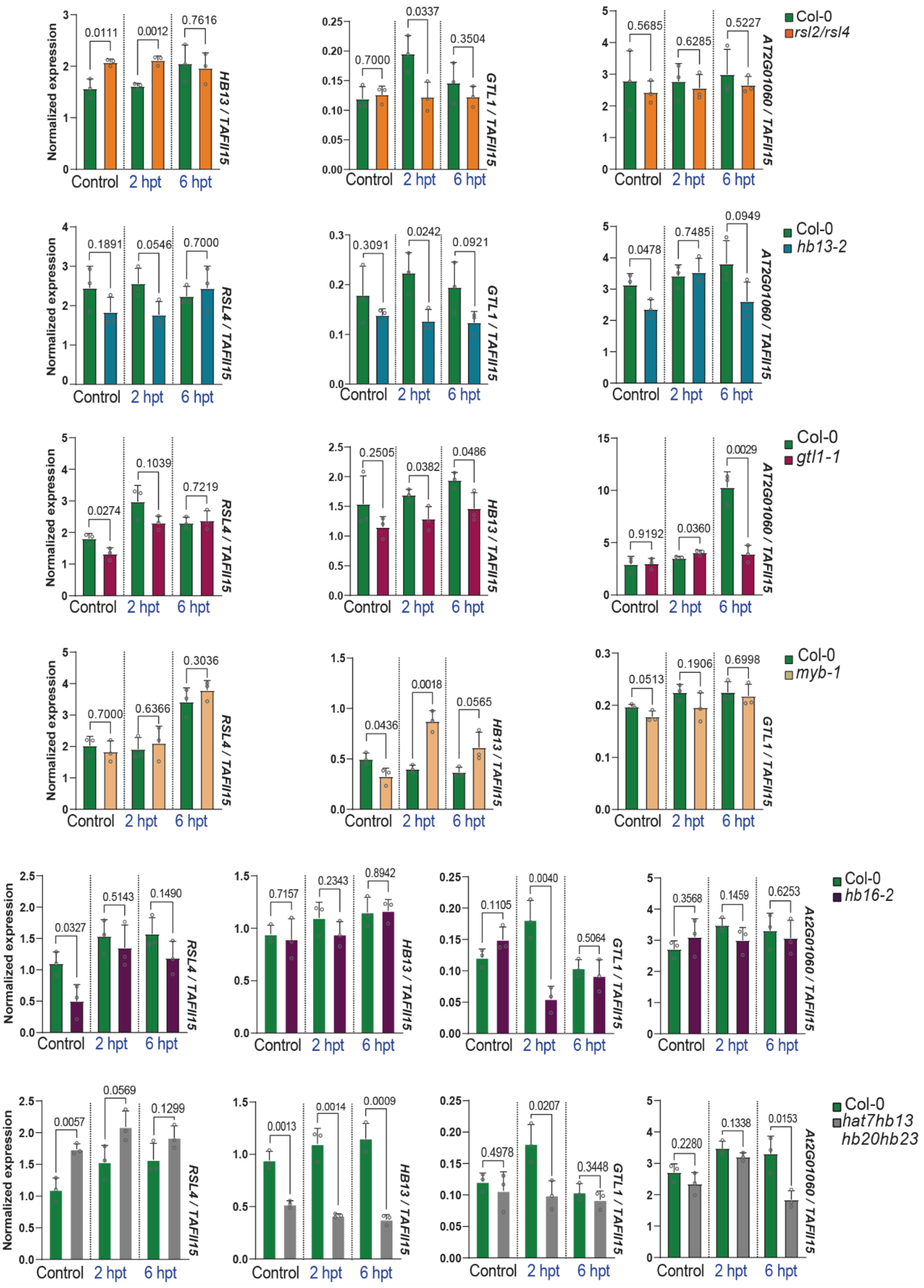


**Figure S11**. **Expression levels of the main RSL4-gene RH regulatory network nodes.** This includes RSL4, AtHB13 and GTL1, and MYB-like (AT2G01060) in Wt and in the mutants *rsl2/rsl4*, *athb13-2*, *gtl1-1*, *myb-like*, *athb16-2* and *athb3(hat7) athb13 athb20 athb23*. The mean and standard deviation is plotted, followed by parametric and non-parametric T-tests on populations paired treatment-wise.

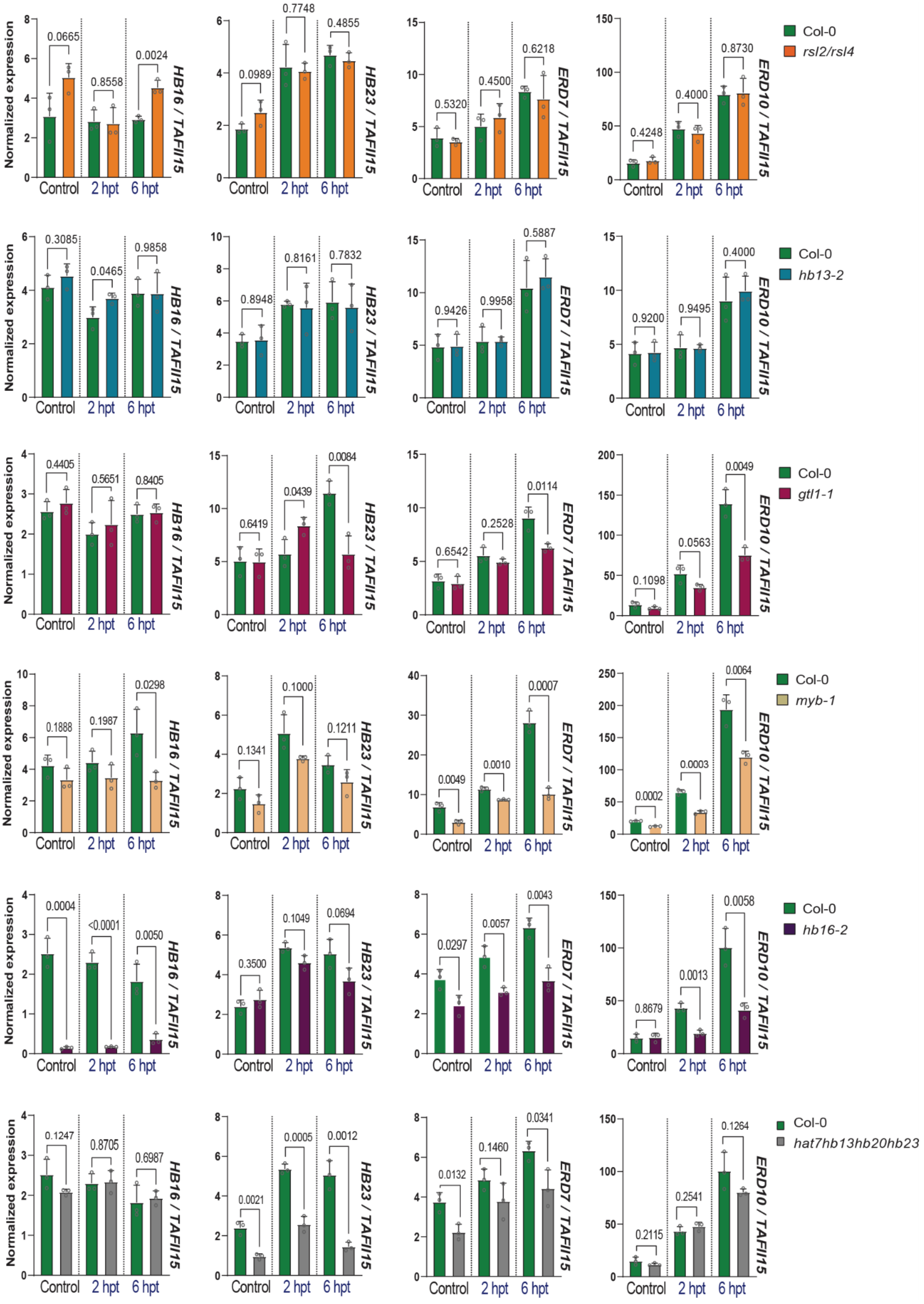


**Figure S12**. **Expression levels of the RSL4-downstream RH gene regulatory network nodes.** Expression levels of the RSL4-downstream genes ERD7/ERD10 and AtHB16/AtHB23 in Wt and in the mutants *rsl2/rsl4*, *athb13-2*, *gtl1-1* and *myb-like 1*, *athb16-2* and *athb3(hat7) athb13 athb20 athb23*. The mean and standard deviation is plotted, followed by parametric and non-parametric T-tests on populations paired treatment-wise.

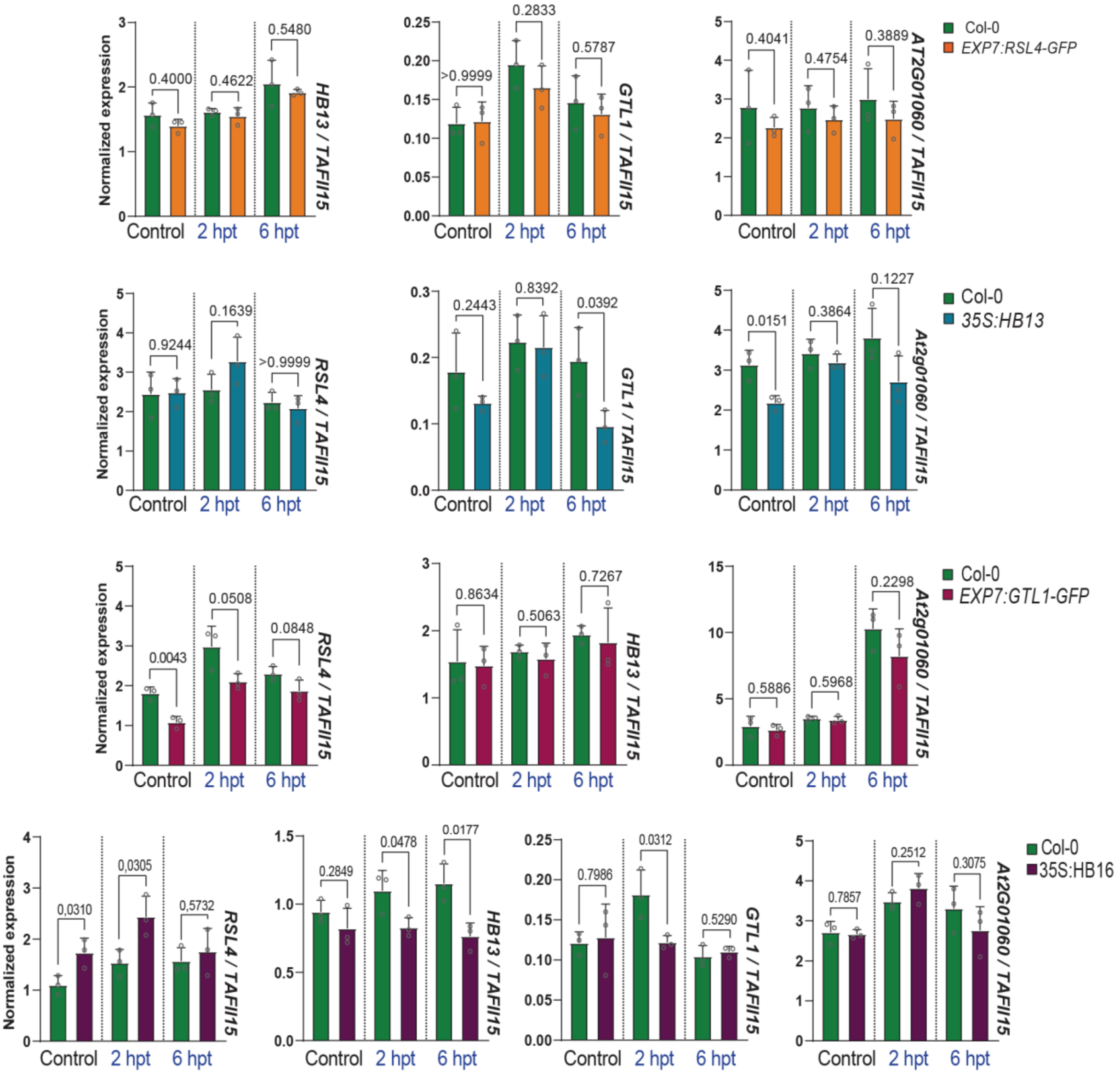


**Figure S13**. **Expression levels of the main RSL4-gene RH regulatory network nodes.** Expression levels of the RH main gene regulatory network nodes RSL4, AtHB13 and GTL1, and MYB-like (AT2G01060) in Wt and in the overexpression lines EXP7:RSL4-GFP, EXP7:GTL1-GFP, 35S:AtHB13 and 35S:AtHB16. The mean and standard deviation is plotted, followed by parametric and non-parametric T-tests on populations paired treatment-wise.

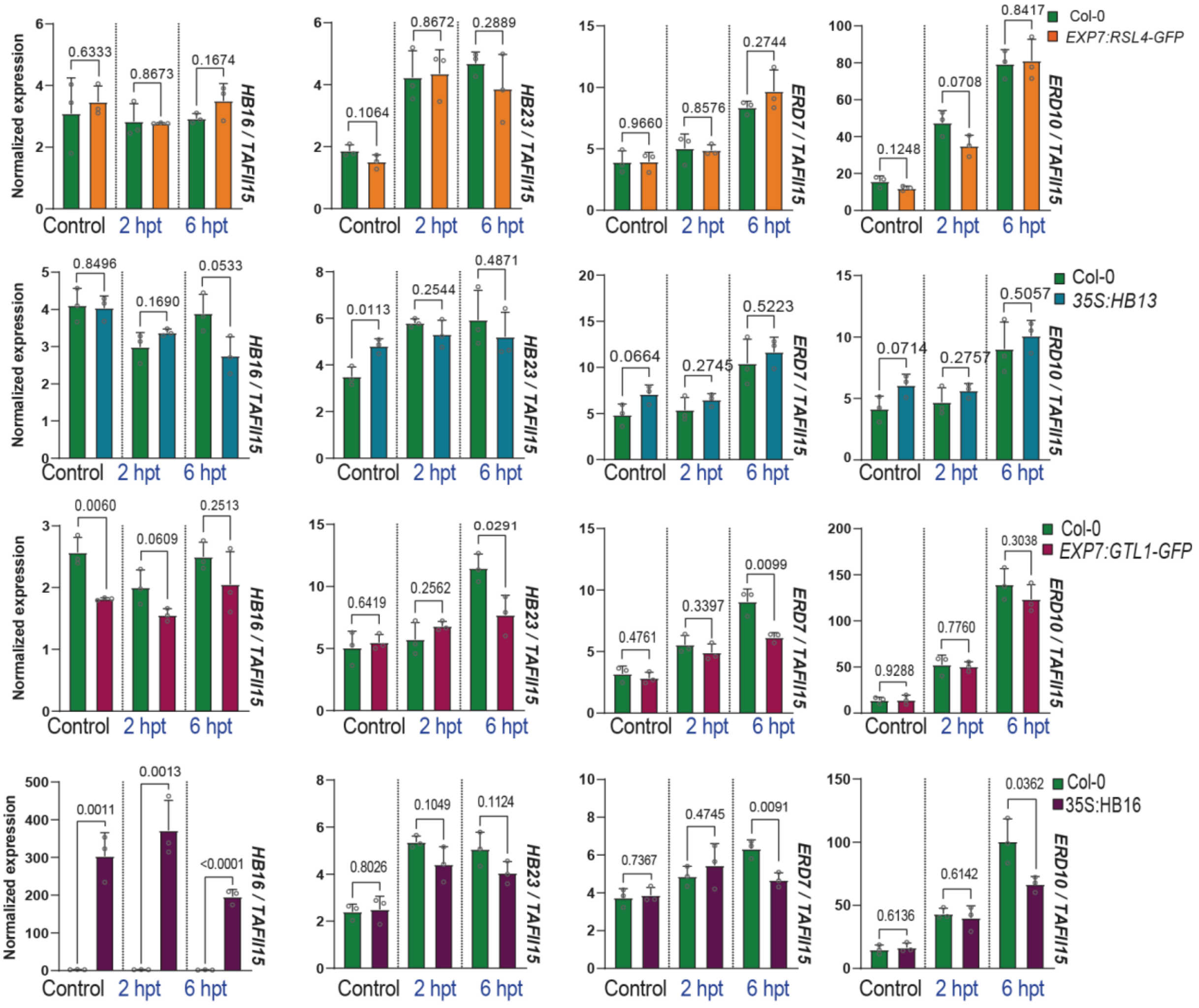


**Figure S14**. Expression levels of the RSL4-downstream genes ERD7/ERD10 and AtHB16/AtHB23 in Wt and in the overexpression lines EXP7:RSL4-GFP, EXP7:GTL1-GFP, 35S:AtHB13 and 35S:AtHB16. The mean and standard deviation is plotted, followed by parametric and non-parametric T-tests on populations paired treatment-wise.

**Supplementary Figure S14.** Predicted aligned error (PAE) maps of the best 35 positive protein-protein interactions (PPIs) identified via AFM predictions. The PAE indicates the expected positional error (in Å) of the alpha carbons (Cα) between two proteins when aligned (low is better). Protein labels (derived from UniProt ID), Local Interaction Score (LIS) and Local Interaction Area (LIA), and interface pTM (ipTM) metric derived from AFM scores for each prediction are shown. Positive PPIs were determined by optimal thresholds for best LIS/LIA and average LIS/LIA. The blue color in the PAE maps, especially in the interface regions (top right and bottom left), indicates areas with high confidence in the potential interaction between proteins. The red color areas indicate regions with no interaction. See the detail about the thresholds in the methods and all ranks for all prediction results, can be found in **Supplementary Table S8**.

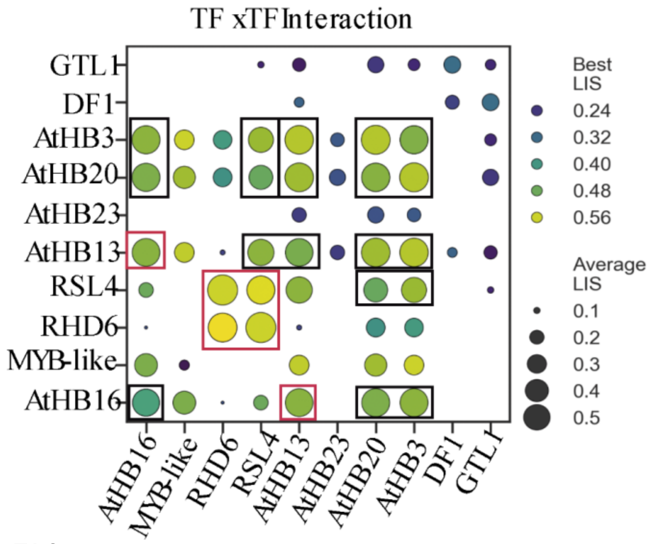


**Best 1 RHD6-RHD6**
**Best 2 RSL4-RSL4**
**Best 3 RHD6-RSL4**
**Best 4 MYB-like-HB3**
**Best 5 MYB-Like HB13**
**Best 6 HB13-HB3**
**Best 7 HB20-HB3**
**Best 8 MYB-like-HB20**
**Best 9 HB13-HB20**
**Best 10 RSL4-HB3**
**Best 11 HB16-HB3**
**Best 12 RSL4-HB13**
**Best 13 HB16-HB13**
**Best 14 HB20-HB20**
**Best 15 HB3-HB3**
**Best 16 HB16-HB20**
**Best 17 HB16-MYB-like**
**Best 18 HB13-HB13**
**Best 19 HB16-RSL4**
**Best 20 RSL4-HB20**
**Best 21 HB16-HB16**
**Best 22 RHD6-HB3**
**Best 23 RHD6-HB20**
**Best 24 DF1-GTL1**
**Best 25 HB13-DF1**
**Best 26 HB23-HB3**
**Best 27 HB23-HB20**
**Best 28 HB16-RHD6**
**Best 29 DF1-DF1**
**Best 30 RHD6-HB13**
**Best 31 HB13-HB23**
**Best 32 HB20-GTL1**
**Best 33 HB3-GTL1**
**Best 34 GTL1-GTL1**
**Best 35 RSL4-GTL1**

**Best 1 RHD6-RHD6**
Protein 1: A0A178W430_ARATH RHD6(green),
Protein 2: A0A178W430_ARATH RHD6 (magenta)
Green: LIA (A), Magenta: LIA (B)
LIS: 0.585
LIA: 7131
ipTM: 0.29

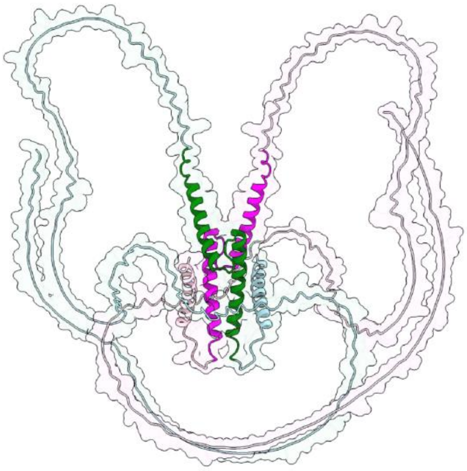

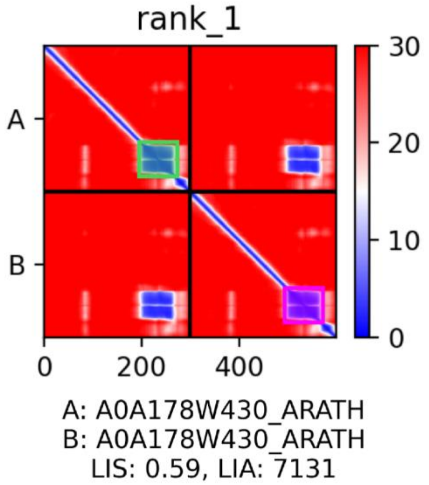

**Best 2 RSL4-RSL4**
Protein 1: A0A384L1Z0_ARATH RSL4 (green),
Protein 2: A0A384L1Z0_ARATH RSL4 (magenta)
Green: LIA (A), Magenta: LIA (B)
LIS: 0.571
LIA: 10791
ipTM: 0.37

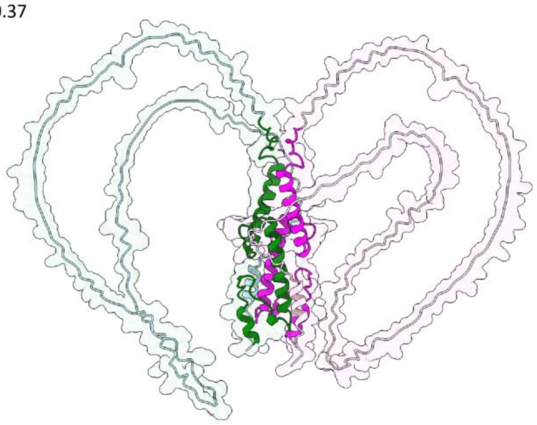

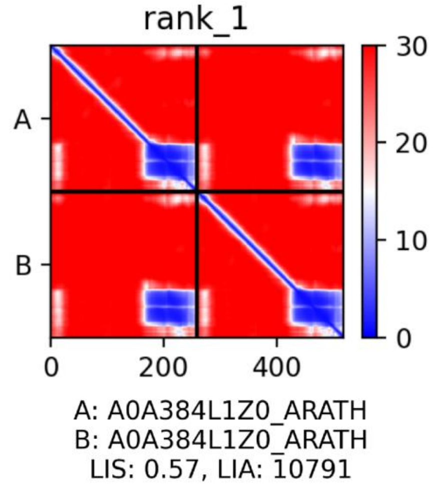


**Best 3 RHD6-RSL4**
Protein 1: A0A178W430_ARATH (green),
Protein 2: A0A384L1Z0_ARATH (magenta)
Green: LIA (A), Magenta: LIA (B)
LIS: 0.560
LIA: 9426
ipTM: 0.40

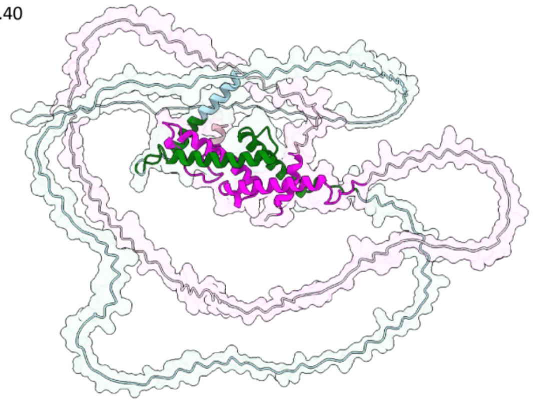

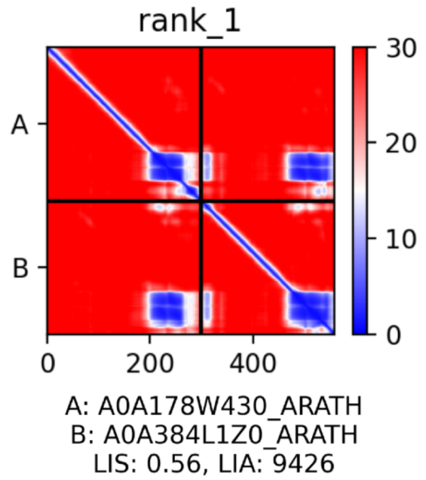


**Best 4 MYB-like-HB3**
Protein 1: A0A178VZU9_ARATH (green),
Protein 2: B5RID5_ARATH (magenta)
Green: LIA (A), Magenta: LIA (B)
LIS: 0.560
LIA: 10402
ipTM: 0.40

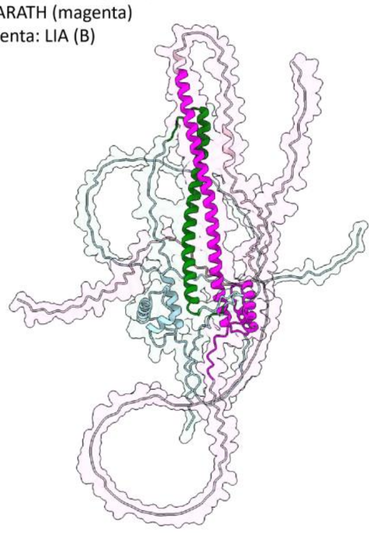

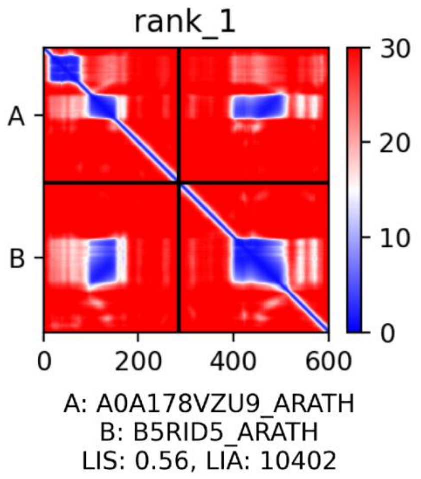

**Best 5 MYB-Like HB13**
Protein 1: A0A178VZU9_ARATH (green),
Protein 2: A0A654EP87_ARATH (magenta)
Green: LIA (A), Magenta: LIA (B)
LIS: 0.554
LIA: 10167
ipTM: 0.39

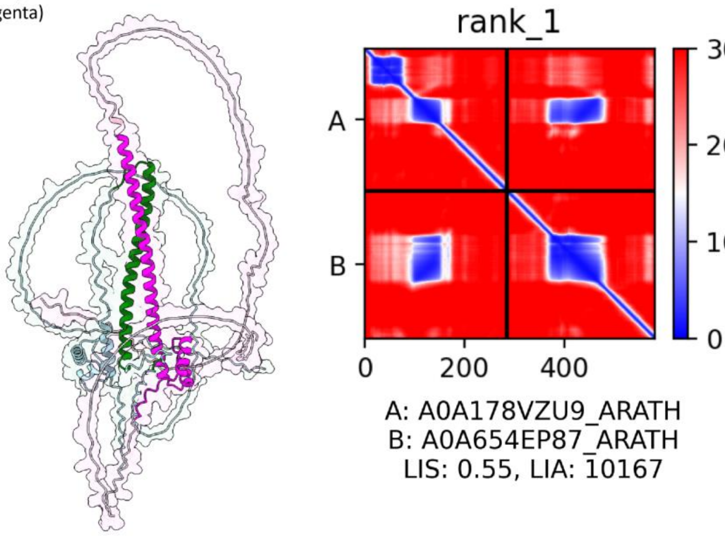


**Best 6 HB13-HB3**
Protein 1: A0A654EP87_ARATH HB13 (green),
Protein 2: B5RID5_ARATH HB3 (magenta)
Green: LIA (A), Magenta: LIA (B)
LIS: 0.547
LIA: 21622
ipTM: 0.44

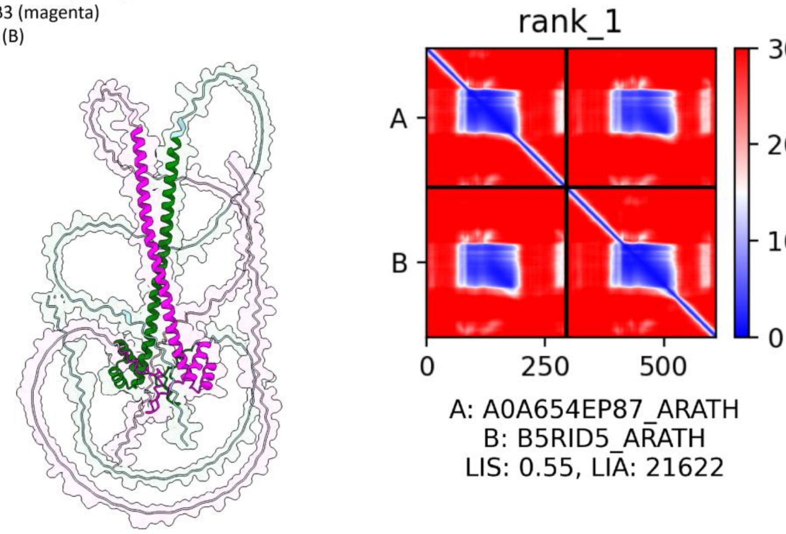


**Best 7 HB20-HB3**
Protein 1: ATB20_ARATH HB20 (green),
Protein 2: B5RID5_ARATH HB3 (magenta)
Green: LIA (A), Magenta: LIA (B)
LIS: 0.546
LIA: 22142
ipTM: 0.44

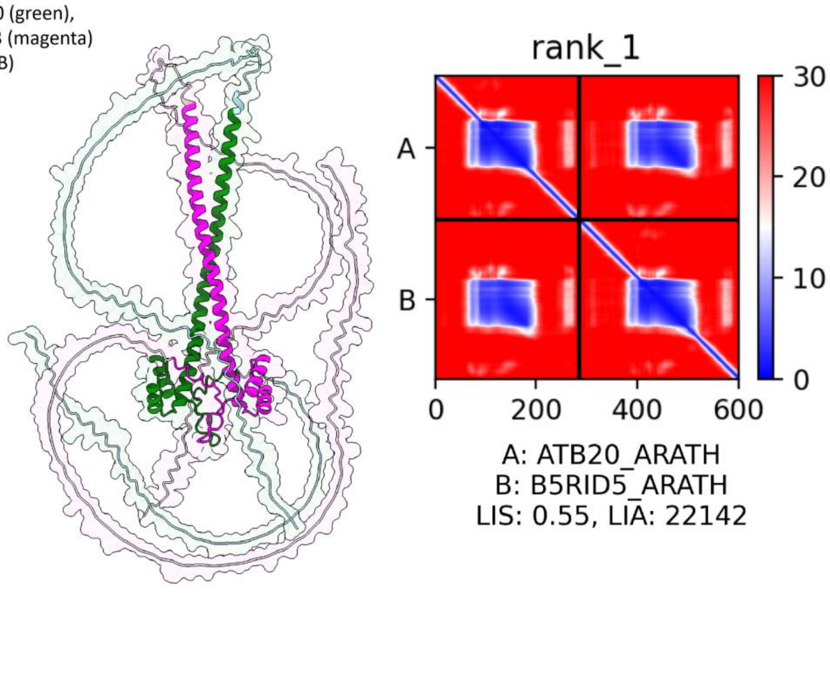

**Best 8 MYB-like-HB20**
Protein 1: A0A178VZU9_ARATH MYB-like (green),
Protein 2: ATB20_ARATH HB20 (magenta)
Green: LIA (A), Magenta: LIA (B)
LIS: 0.533
LIA: 10935
ipTM: 0.41

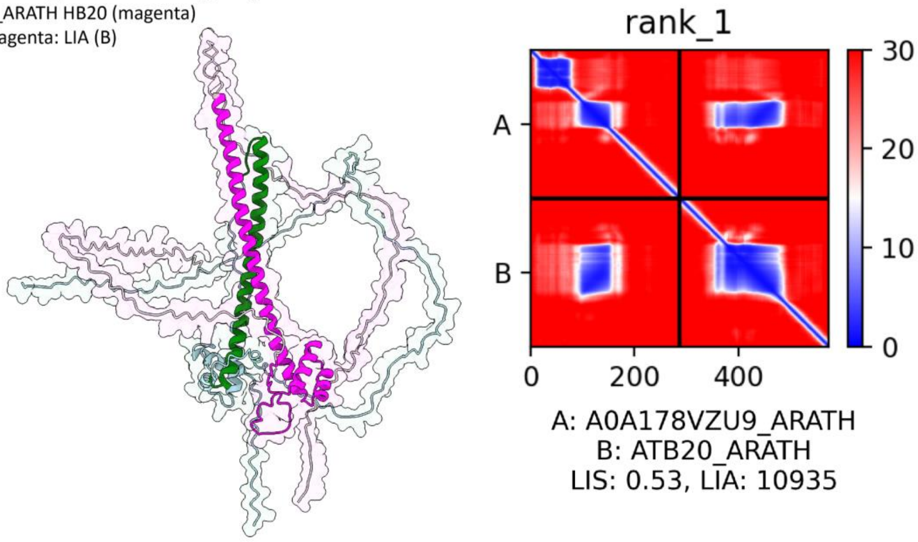


178

**Best 9 HB13-HB20**
Protein 1: A0A654EP87_ARATH HB13 (green),
Protein 2: ATB20_ARATH HB20 (magenta)
Green: LIA (A), Magenta: LIA (B)
LIS: 0.531
LIA: 22037
ipTM: 0.44

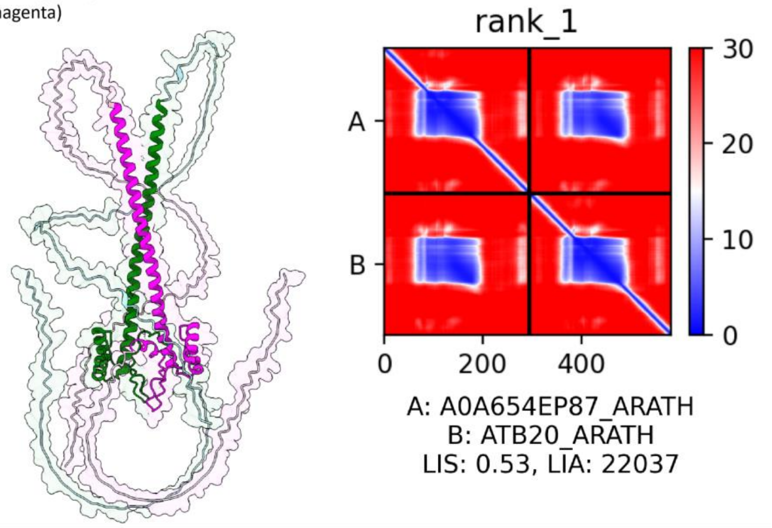


179

**Best 10 RSL4-HB3**
Protein 1: A0A384L1Z0_ARATH RSL4 (green),
Protein 2: B5RID5_ARATH HB3 (magenta)
Green: LIA (A), Magenta: LIA (B)
LIS: 0.527
LIA: 13586
ipTM: 0.40

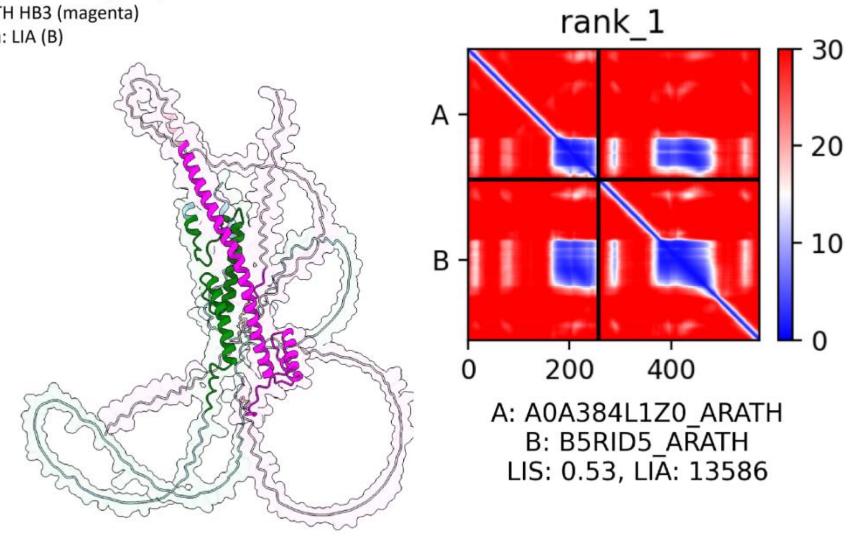


180

**Best 11 HB16-HB3**
Protein 1: A0A178V0P3_ARATH HB16 (green),
Protein 2: B5RID5_ARATH HB3 (magenta)
Green: LIA (A), Magenta: LIA (B)
LIS: 0.516
LIA: 21075
ipTM: 0.48

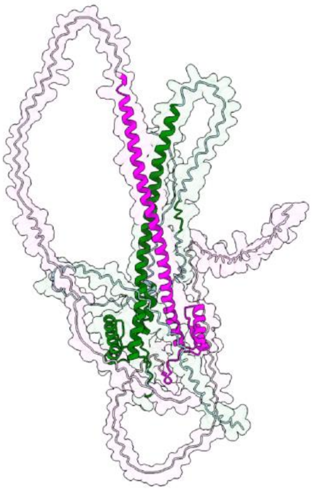

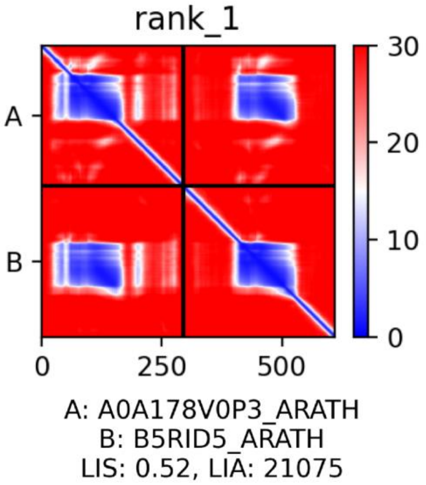


**Best 12 RSL4-HB13**
Protein 1: A0A384L1Z0_ARATH RSL4 (green),
Protein 2: A0A654EP87_ARATH HB13 (magenta)
Green: LIA (A), Magenta: LIA (B)
LIS: 0.515
LIA: 13876
ipTM: 0.38

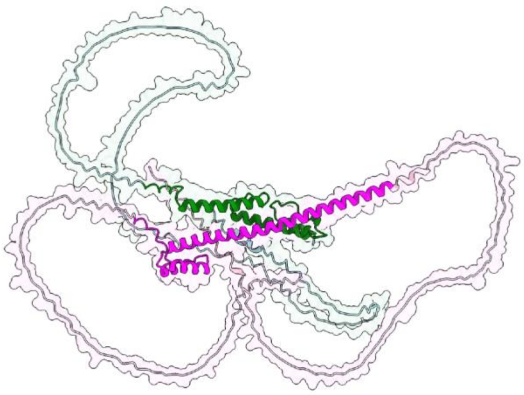

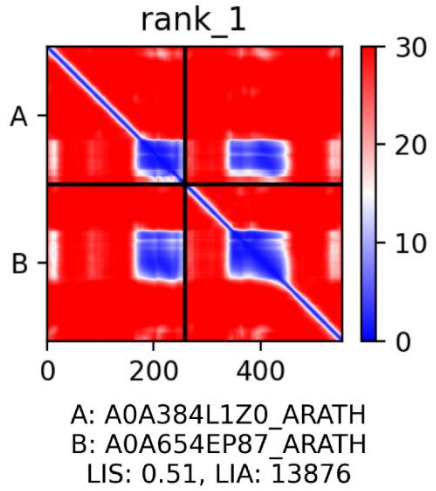


**Best 13 HB16-HB13**
Protein 1: A0A178V0P3_ARATH HB16 (green),
Protein 2: A0A654EP87_ARATH HB13 (magenta)
Green: LIA (A), Magenta: LIA (B)
LIS: 0.512
LIA: 21857
ipTM: 0.48

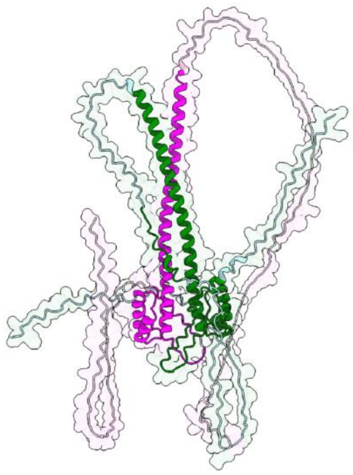

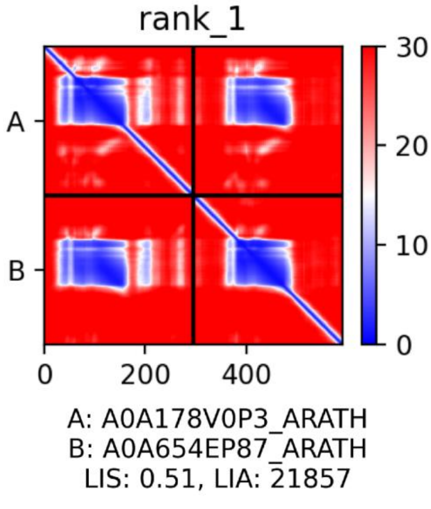

**Best 14 HB20-HB20**
Protein 1: ATB20_ARATH HB20 (green),
Protein 2: ATB20_ARATH HB20 (magenta)
Green: LIA (A), Magenta: LIA (B)
LIS: 0.510
LIA: 22487
ipTM: 0.42

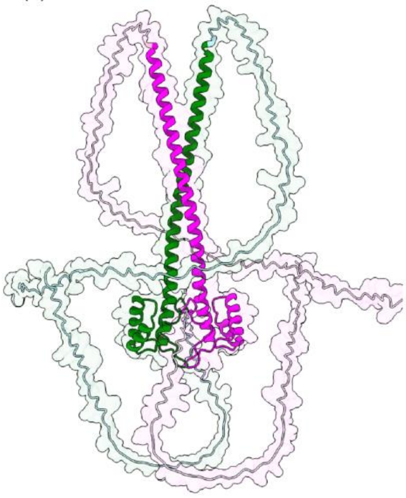

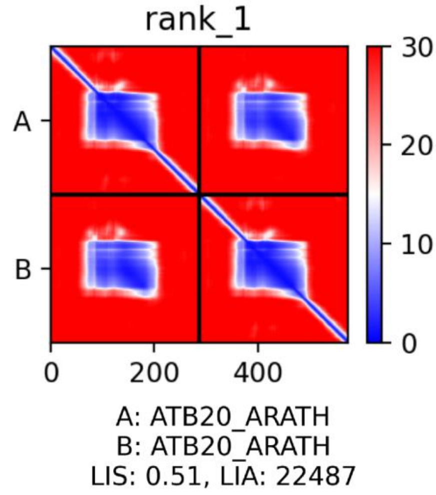


**Best 15 HB3-HB3**
Protein 1: B5RID5_ARATH HB3 (green),
Protein 2: B5RID5_ARATH HB3 (magenta)
Green: LIA (A), Magenta: LIA (B)
LIS: 0.504
LIA: 23276
ipTM: 0.42

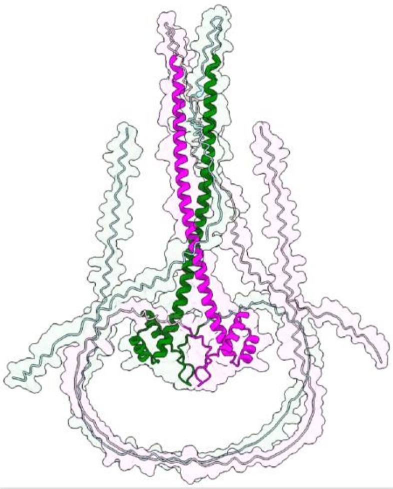

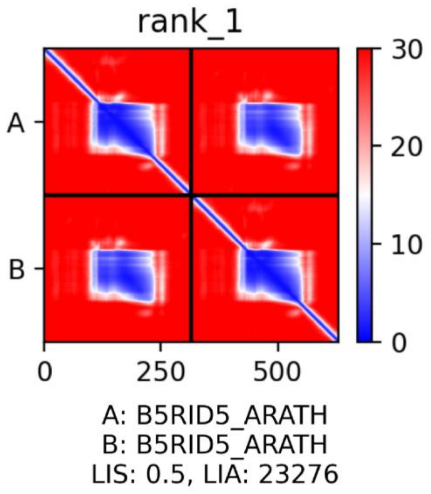


**Best 16 HB16-HB20**
Protein 1: A0A178V0P3_ARATH HB16 (green),
Protein 2: ATB20_ARATH HB20 (magenta)
Green: LIA (A), Magenta: LIA (B)
LIS: 0.501
LIA: 23164
ipTM: 0.49

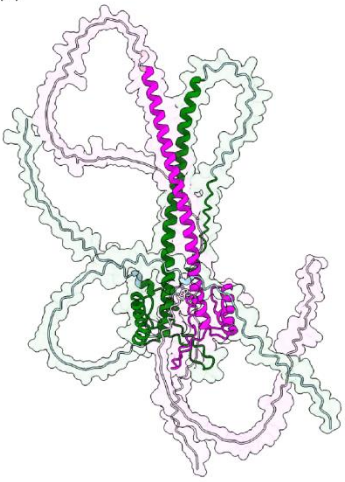

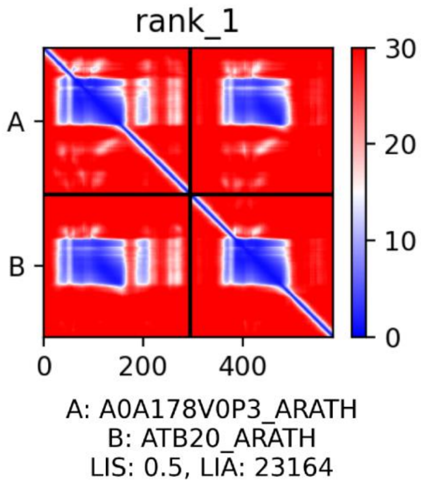

**Best 17 HB16-MYB-like**

Protein 1: A0A178V0P3_ARATH HB16 (green),
Protein 2: A0A178VZU9_ARATH MYB-like (magenta)
Green: LIA (A), Magenta: LIA (B)
LIS: 0.500
LIA: 11667
ipTM: 0.47

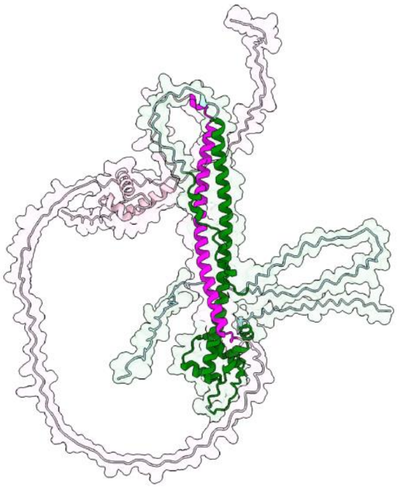

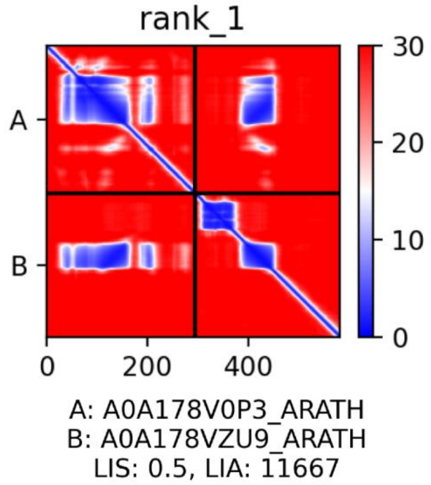


**Best 18 HB13-HB13**

Protein 1: A0A654EP87_ARATH HB13 (green),
Protein 2: A0A654EP87_ARATH HB13 (magenta)
Green: LIA (A), Magenta: LIA (B)
LIS: 0.496
LIA: 20702
ipTM: 0.43

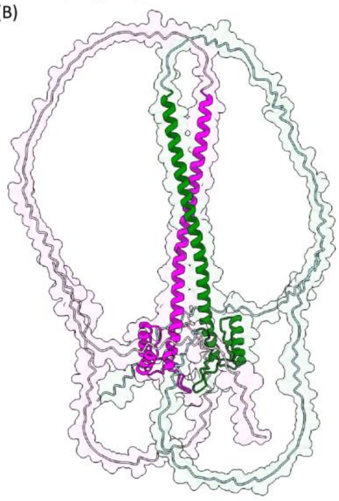

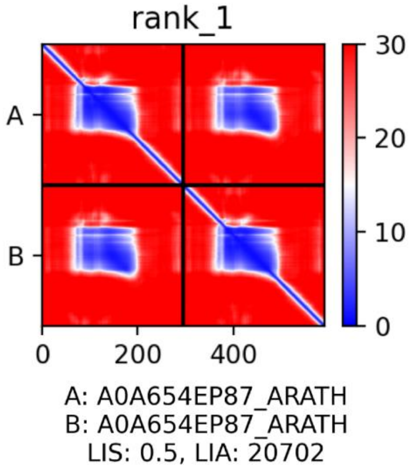


**Best 19 HB16-RSL4**

Protein 1: A0A178V0P3_ARATH HB16 (green),
Protein 2: A0A384L1Z0_ARATH RSL4 (magenta)
Green: LIA (A), Magenta: LIA (B)
LIS: 0.480
LIA: 15809
ipTM: 0.45

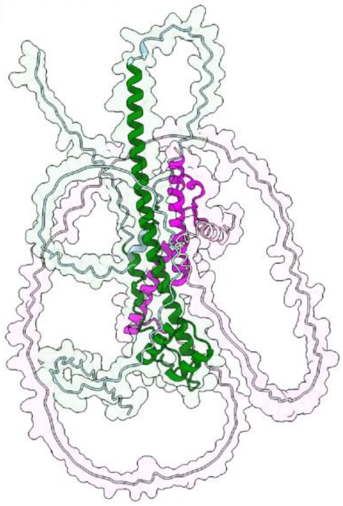

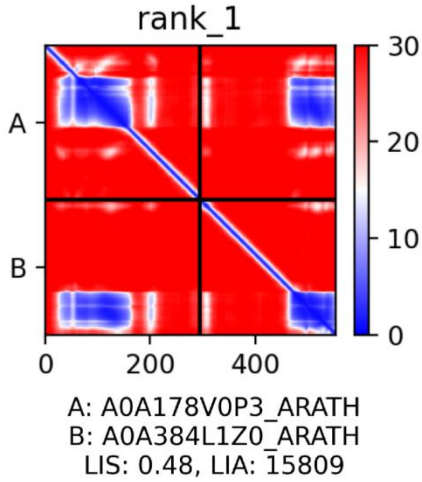

**Best 20 RSL4-HB20**
Protein 1: A0A384L1Z0_ARATH RSL4 (green),
Protein 2: ATB20_ARATH HB20 (magenta)
Green: LIA (A), Magenta: LIA (B)
LIS: 0.475
LIA: 13684
ipTM: 0.40

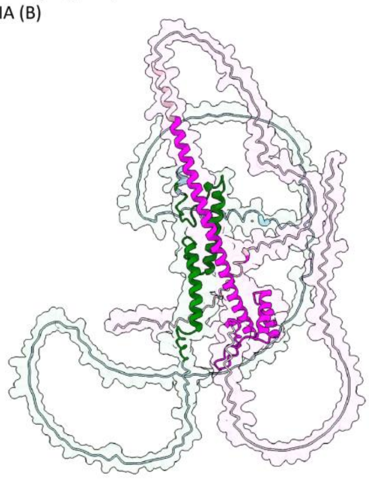

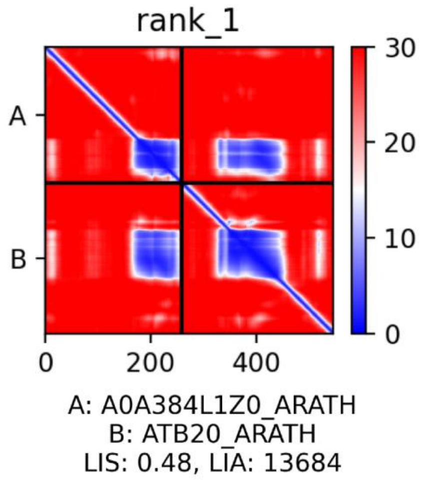


**Best 22 RHD6-HB3**
Protein 1: A0A178W430_ARATH RHD6 (green),
Protein 2: B5RID5_ARATH HB3 (magenta)
Green: LIA (A), Magenta: LIA (B)
LIS: 0.409
LIA: 10941
ipTM: 0.36

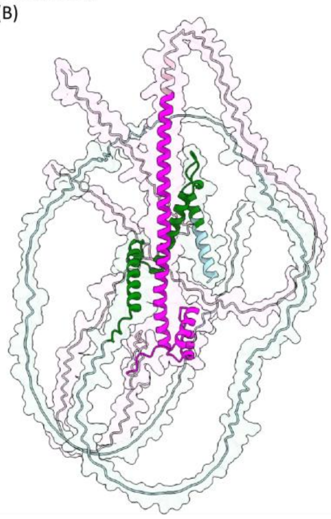

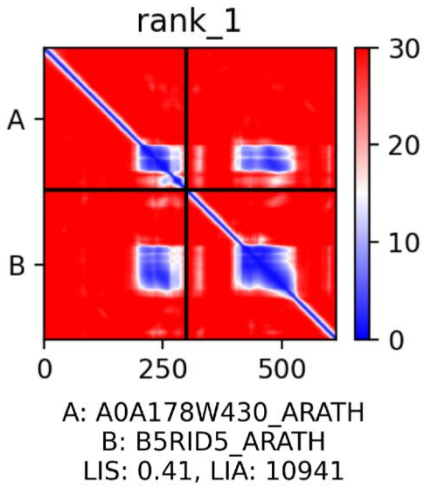


**Best 23 RHD6-HB20**
Protein 1: A0A178W430_ARATH RHD6 (green),
Protein 2: ATB20_ARATH HB20 (magenta)
Green: LIA (A), Magenta: LIA (B)
LIS: 0.392
LIA: 10939
ipTM: 0.38

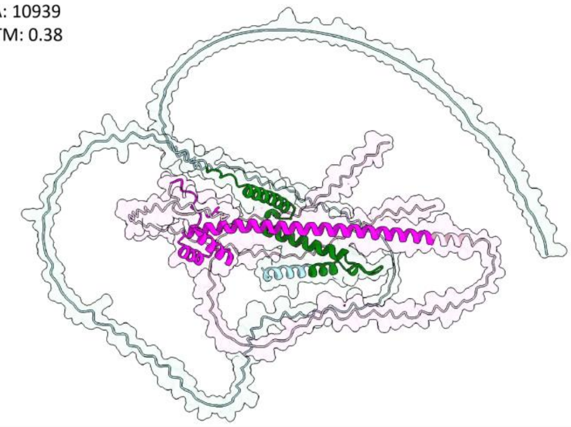

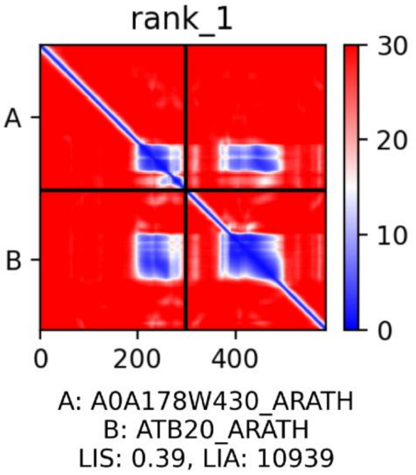

**Best 24 DF1-GTL1**
Protein 1: DF1_ARATH (green),
Protein 2: GTL1_ARATH (magenta)
Green: LIA (A), Magenta: LIA (B)
LIS: 0.328
LIA: 10473
ipTM: 0.25

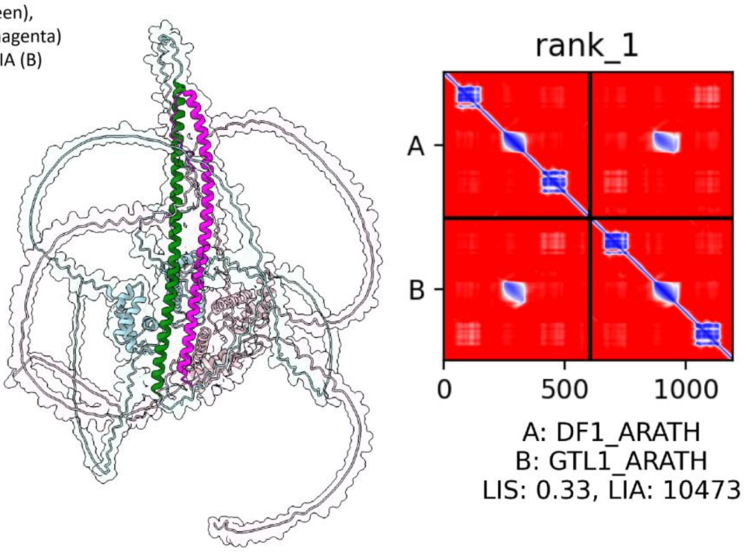


**Best 25 HB13-DF1**
Protein 1: A0A654EP87_ARATH HB13 (green),
Protein 2: DF1_ARATH (magenta)
Green: LIA (A), Magenta: LIA (B)
LIS: 0.308
LIA: 10108
ipTM: 0.34

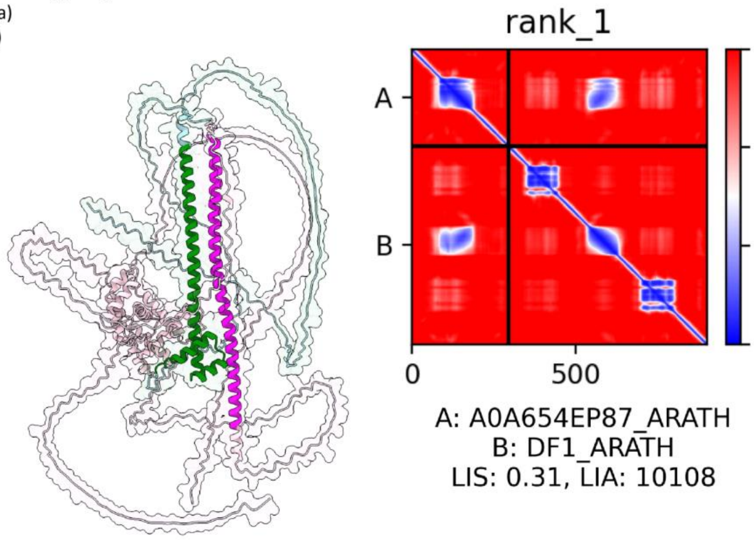


**Best 26 HB23-HB3**
Protein 1: A0A654G6H4_ARATH HB23 (green),
Protein 2: B5RID5_ARATH HB3 (magenta)
Green: LIA (A), Magenta: LIA (B)
LIS: 0.293
LIA: 9287
ipTM: 0.28

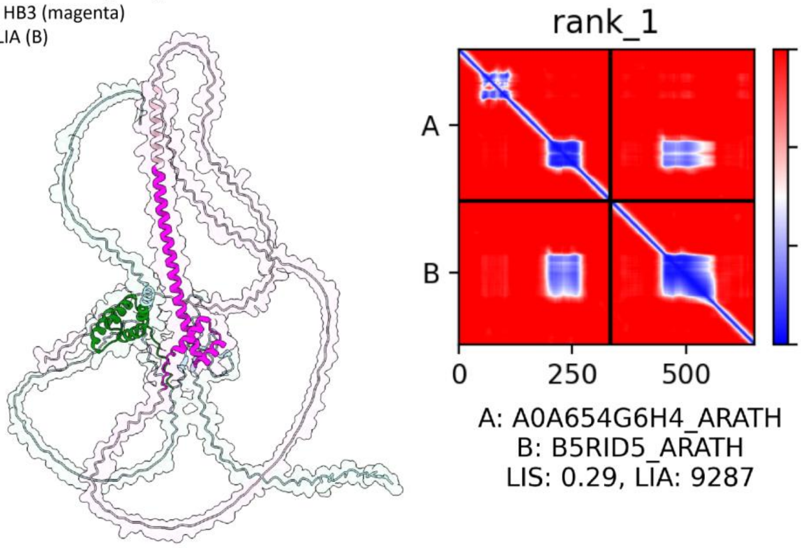

**Best 27 HB23-HB20**
Protein 1: A0A654G6H4_ARATH HB23 (green),
Protein 2: ATB20_ARATH HB20 (magenta)
Green: LIA (A), Magenta: LIA (B)
LIS: 0.283
LIA: 9035
ipTM: 0.29

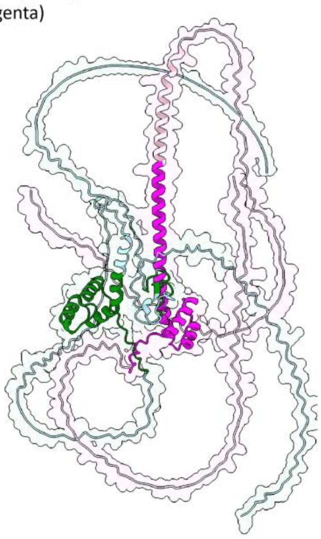

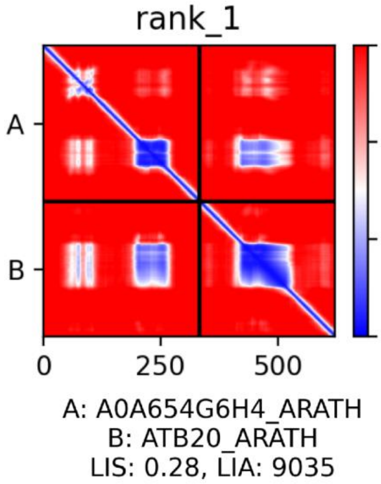


196

**Best 28 HB16-RHD6**
Protein 1: A0A178V0P3_ARATH HB16- (green),
Protein 2: A0A178W430_ARATH RHD6 (magenta)
Green: LIA (A), Magenta: LIA (B)
LIS: 0.283
LIA: 4363
ipTM: 0.34

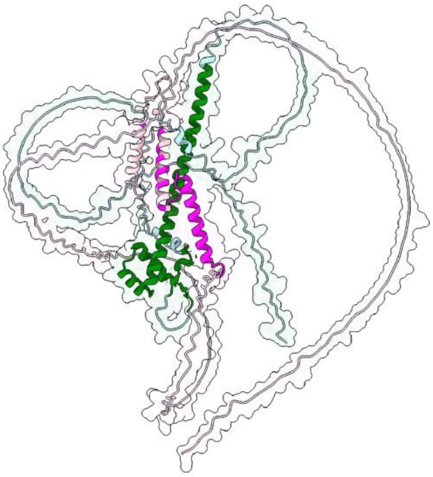

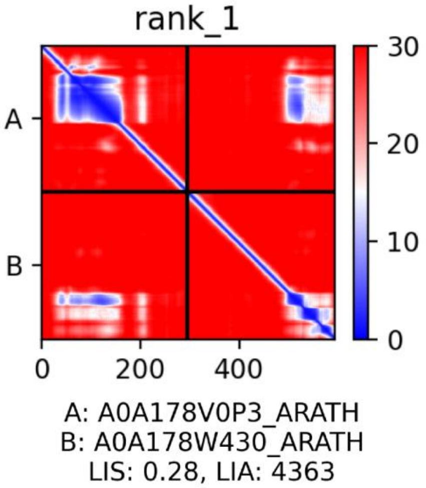


197

**Best 29 DF1-DF1**
Protein 1: DF1_ARATH (green),
Protein 2: DF1_ARATH (magenta)
Green: LIA (A), Magenta: LIA (B)
LIS: 0.258
LIA: 9908
ipTM: 0.26

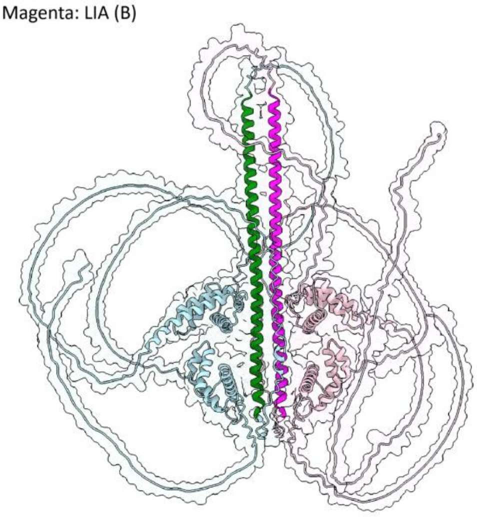

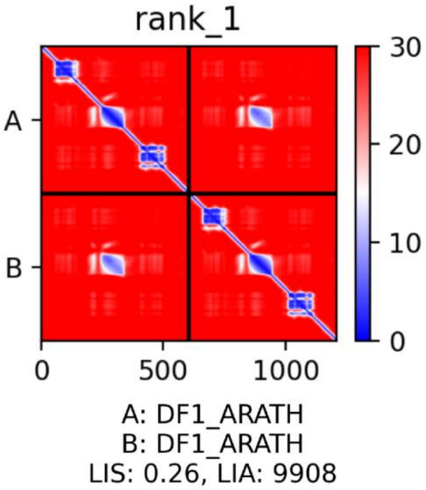


198

**Best 30 RHD6-HB13**
Protein 1: A0A178W430_ARATH RHD6 (green),
Protein 2: A0A654EP87_ARATH  HB13 (magenta)
Green: LIA (A), Magenta: LIA (B)
LIS: 0.258
LIA: 7414
ipTM: 0.30

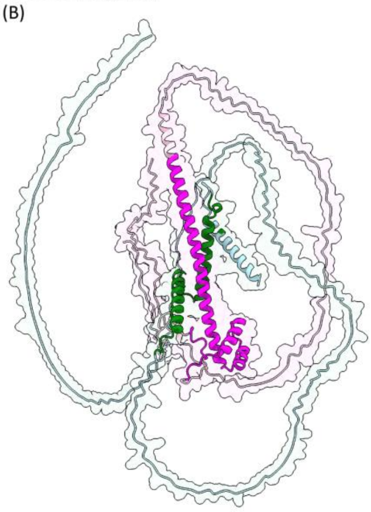

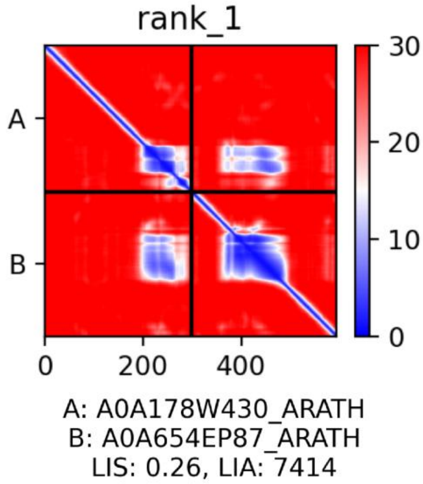


199

**Best 31 HB13-HB23**
Protein 1: A0A654EP87_ARATH HB13 (green),
Protein 2: A0A654G6H4_ARATH HB23 (magenta)
Green: LIA (A), Magenta: LIA (B)
LIS: 0.249
LIA: 7943
ipTM: 0.27

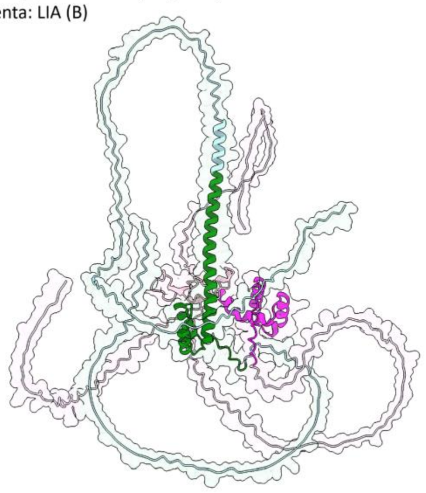

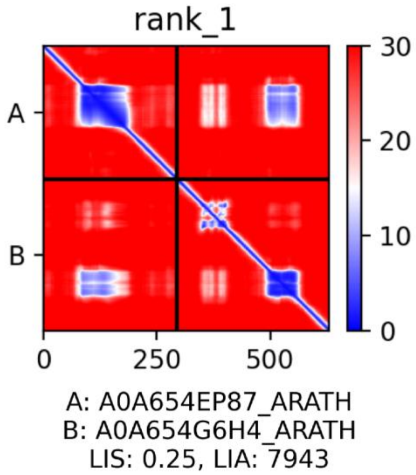


200

**Best 32 HB20-GTL1**
Protein 1: ATB20_ARATH (green),
Protein 2: GTL1_ARATH (magenta)
Green: LIA (A), Magenta: LIA (B)
LIS: 0.242
LIA: 11892
ipTM: 0.31

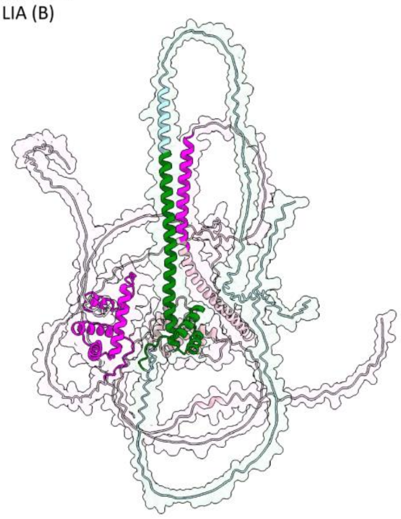

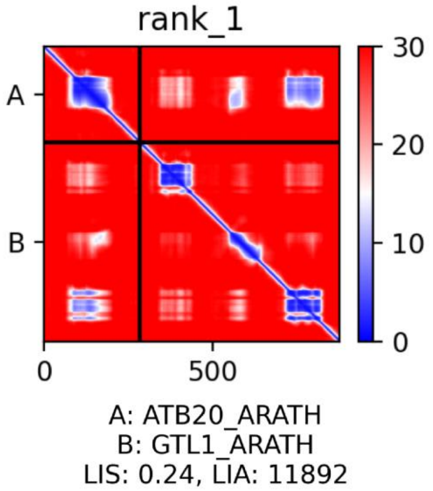


201

**Best 33 HB3-GTL1**
Protein 1: B5RID5_ARATH HB3 (green),
Protein 2: GTL1_ARATH (magenta)
Green: LIA (A), Magenta: LIA (B)
LIS: 0.221
LIA: 11922
ipTM: 0.31

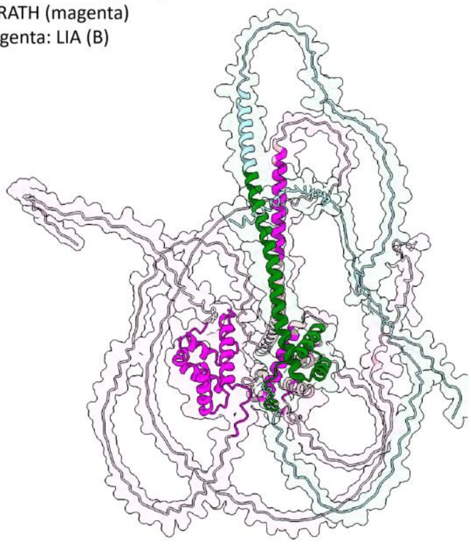

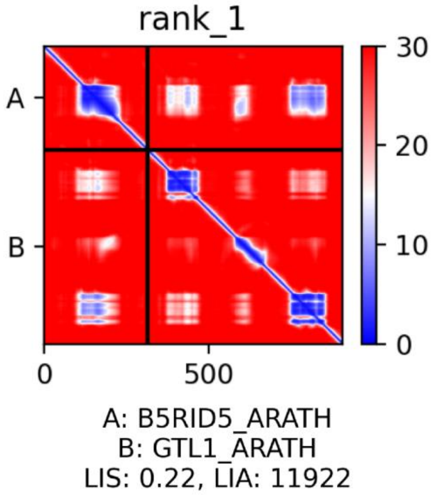


202

**Best 34 GTL1-GTL1**
Protein 1: GTL1_ARATH (green),
Protein 2: GTL1_ARATH (magenta)
Green: LIA (A), Magenta: LIA (B)
LIS: 0.220
LIA: 10050
ipTM: 0.24

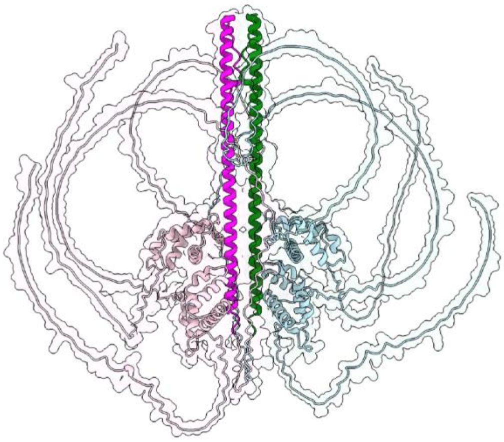

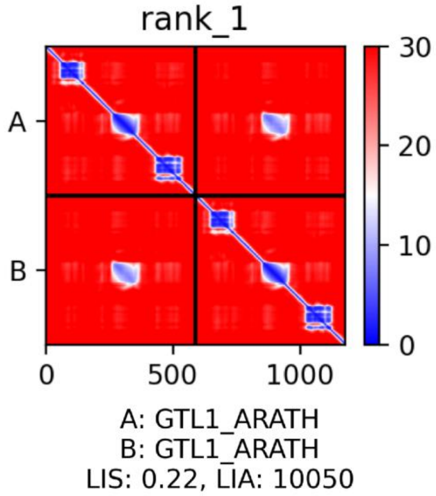


203

**Best 35 RSL4-GTL1**
Protein 1: A0A384L1Z0_ARATH RSL4 (green),
Protein 2: GTL1_ARATH (magenta)
Green: LIA (A), Magenta: LIA (B)
LIS: 0.218
LIA: 3710
ipTM: 0.33

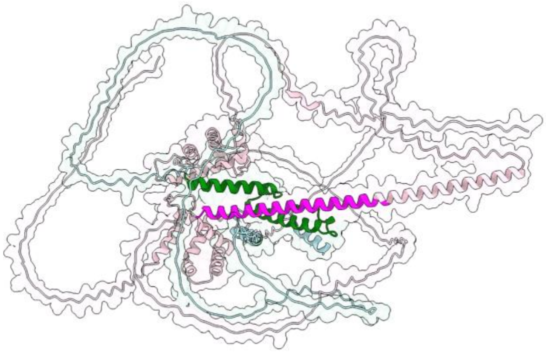

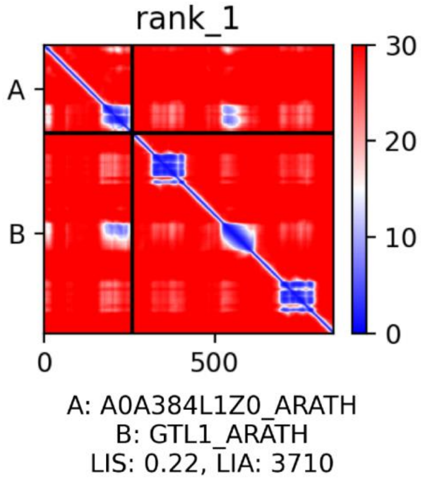


204

**Supplementary Table S6**. Mutants and transgenic lines used and generated in this study.

| Mutants and transgenic lines | | | | | |
|---|---|---|---|---|---|
| **Locus** | **Gene name** | **Gene expression effect** | **Line name** | **ABRC name/stock number** | **Reference** |
| **AT4G33880** | ***RSL2*** | mutant | *rsl2* | | Yi et al., 2010 |
| **AT1G27740** | ***RSL4*** | mutant | *rsl4* | | Yi et al., 2010 |
| | | mutant | *rsl2 rsl4* | | Yi et al., 2010 |
| | | O.E. | EXP7:RSL4 | | Yi et al., 2010 |
| **AT1G69780** | ***AtHB13*** | mutant | *athb13-1* | SAIL_893_G05 | Ribone et al., 2015 |
| | | mutant | *athb13-2* | GABI_859H06 | Ribone et al., 2015 |
| | | O.E. | 35S:AtHB13 | | Cabello et al., 2012 |
| | | reporter GUS | HB13:GUS | | Ribone et al., 2015 |
| **AT2G01060** | ***MYB-like*** | mutant | *myb-like 1* | SALKseq_8995 | this study |
| **AT1G33240** | ***GTL1*** | complemented | GTL1:GTL1-GFP/*gtl1-1* | | Shibata et al., 2018 |
| | | O.E. | EXP7:GTL1-GFP/Col-0 | | Shibata et al., 2018 |
| | | mutant | *gtl1-1* | WiscDsLox413-416C9 | Shibata et al., 2018 |
| | | mutant | *gtl1-1 df1-1* | | Shibata et al., 2018 |
| **AT1G76880** | ***DF1*** | complemented | DF1:DF1-GFP/*df1-1* | | Shibata et al., 2018 |
| | | O.E. | EXP7:DF1-GFP/Col-0 | | Shibata et al., 2018 |
| | | mutant | *df1-1* | SALK_106258 | Shibata et al., 2018 |
| **AT4G40060** | ***AtHB16*** | mutant | *athb16-1* | | This study |
| | | mutant | *athb16-2* | | This study |
| | | O.E. | 35S:AtHB16-2 | | This study |
| | | O.E. | 35S:AtHB16-14 | | This study |
| | | O.E. | 35S:AtHB16 30 | | This study |
| | | reporter GUS | HB16:GUS | | This study |
| **AT5G39760** | ***AtHB23*** | O.E. | 35S:AtHB23-7F | | Perotti et al. 2021, 2022 |
| | | O.E. | 35S:AtHB23-5H | | Perotti et al. 2021, 2022 |
| | | O.E. | 35S:AtHB23-6A | | Perotti et al. 2021, 2022 |
| | | silenced | amiR-AtHB23-1B | | Perotti et al. 2021, 2022 |
| | | silenced | amiR-AtHB23-20C | | Perotti et al. 2021, 2022 |

| | | | | | |
|---|---|---|---|---|---|
| | | silenced | amiR-AtHB23-9C | | Perotti et al. 2021, 2022 |
| | | reporter GUS | HB23:GUS | | Perotti et al. 2021 |
| **AT5G15150**<br>**AT1G69780**<br>**AT3G01220**<br>**AT5G39760** | ***AtHB3***<br>***AtHB13***<br>***AtHB20***<br>***AtHB23*** | CRISPR mutant<br>CRISPR mutant<br>CRISPR mutant<br>CRISPR mutant | *Athb3 athb13 athb20 athb23*<br>*athb3 athb20 athb23*<br>*athb13 athb20 athb23* | | Nolan et al. 2023 |
| **AT2G17840** | ***ERD7*** | mutant | *erd7-1* | WISCDSLOX452E10 | this work |
| **AT1G20450** | ***ERD10*** | mutant | *erd10-1* | SALK_087789C | this work |

207

**Supplementary Table S7**. Oligonucleotide list for RT-qPCR analysis. Primer sequences are shown next to their corresponding targeted genes. F= Forward and R= Reverse primers.

| RT-qPCR primer list | | |
|---|---|---|
| **Gene** | **Primer probe** | **Sequence (5' - 3')** |
| ***AT2G01060*** | F primer | TCGATTGGTGAGAGGTGCAG |
| | R primer | AAAGGCGAGGACATGGTCTG |
| ***ERD7*** | F primer | AAGGGCTCGATGCAGCAGGATA |
| | R primer | GGTTTTTGCAAGCGTCGAAGGC |
| ***ERD10*** | F primer | ACCGATTGCTGACATCCCGGAG |
| | R primer | TCTCCAGTGGTCTTGGCGTGAT |
| ***GTL1*** | F primer | ATGGAATTGTTTGAAGGTTTGG |
| | R primer | GACATGACCTCGTGTTCTCG |
| ***AtHB13*** | F primer | TCTCAACTGCGCCGCCATCAAA |
| | R primer | GCGACGGTGGGAAGAAGTGTCG |
| ***RHD6*** | F primer | TGATTTGGTGACAATGCTTGA |
| | R primer | GGAGAGAATGGCATCAATGG |
| ***RSL2*** | F primer | CCCCAATGGAACAAAGGTC |
| | R primer | TCTCGGTGAGCTGAGACCAA |
| ***RSL4*** | F primer | GTGCCAAACGGGACAAAAGT |
| | R primer | TTGTGATGGAACCCCATGTC |
| ***TAFII15*** | F primer | GAATCACGGCCAACAATC |
| | R primer | ACTCTTAGCCAAGTAGTGCTCC |

| Genotyping primer list | | |
|---|---|---|
| **Gene** | **Primer probe** | **Sequence (5' - 3')** |
| ***myb-1*** **(SALKseq_8995.3)** | F primer | CCATGAGAGGGAAAGCCTATC |
| | R primer | AGGACATGGTCTGACTGATGG |
| ***myb-2*** **(SALKseq_034222)** | F primer | AGGAGCAATTTGAACTCCCTC |
| | R primer | TTGGAAACCTGGATTGTTGAC |
| ***erd7-1*** **(WISCDSLOX452E10)** | F primer | TTTAATGCAATTTCCGAGACG |
| | R primer | CAATCTCTTTCCTTTCCCCAG |
| ***erd10-1*** **(SALK_087789C)** | F primer | CTCACCGTCTTCACCTTCTTC |
| | R primer | TGCATTCTTCTCCAATTCTCC |
| **ChIP-PCR primer list** | | |
| **Gene** | **Primer probe** | **Sequence (5' - 3')** |
| ***EXP7*** | F primer | GAAACAAATCCCAATCGGTAC |
| | R primer | ATGAAGTAATCGAATTTGAGACAATC |
| ***RSL4.1*** | F primer | GTGTCGTGAAGGAACAAAGCC |
| | R primer | CATTTGTGTCGTGAACAAGTCGT |
| ***RSL4.2*** | F primer | CGACACATTCTTTGAACCCGTG |
| | R primer | AGAGGACGGGTTACGGTACAT |
| ***GTL1*** | F primer | GACGTCACGTTTGTTTATTTATGGG |
| | R primer | ACAAGTGATTCTTGTATGTTGGCAT |
| ***AtHB13.1*** | F primer | TCCACAGGTCTAGGTCAATCCA |

| | | |
|---|---|---|
| | R primer | AGGGACGAAGCTGGATAGTGA |
| ***AtHB13.1*** | F primer | CTCTGCATGTCTCTCCCAAGG |
| | R primer | CAAGGGGGTTTAGCTTGCGT |
| ***AtHB16.1*** | F primer | GCTCGGGCATCATCATCCAA |
| | R primer | ACGTTCCCGTACACGTTTCAT |
| ***AtHB16.2*** | F primer | ACGTATCTCTCTACTATCCGCTGT |
| | R primer | AGAACAAGTGGTGGAGTCTGTG |
| ***AtHB23.1*** | F primer | CACACAAAAAAGAACAACTCGTGG |
| | R primer | GGCCCAACTACTGTATAAATTGGTC |
| ***AtHB23.2*** | F primer | ATTCCATTGACTTGAACACTTATGC |
| | R primer | TTTCTTTTAACAGCATGCCCTACA |
| ***MYB-like.1*** | F primer | ATCACCATCTTGCTTCCCCC |
| | R primer | TTTGGTGTTCCTGGGTGTGA |
| ***MYB-like.2*** | F primer | GGAAGGCGTCAGGAAGGAC |
| | R primer | CTGAGCACGTTGGCATCATTG |
| ***ERD7*** | F primer | GACCGACCGACATGAGAACAA |
| | R primer | GTCAGATTTGGGAAGACGTGGAT |
| ***ERD10.1*** | F primer | AACTTTCCGGTTCCTTACCCG |
| | R primer | TCGTACACGTAACGGCGAC |
| ***ERD10.2*** | F primer | ATCTGTGGGCTCCTGAAACG |
| | R primer | AGCCATAGCCATCATGCCAC |
| **BIFC assay** | | |

| Gene | Primer probe | Sequence (5' - 3') |
|---|---|---|
| **HB13** | F primer | CACCATGTCTTGTAATAATGGAATGTCT |
| | R primer | TCAATTGTACTGTTGCTGATCAAGCCA |
| **HB16** | F primer | CACCATGAAGAGACTAAGCAGCTCAGAT |
| | R primer | TCAAGTCCAATGATCTGAAGCAGAGTA |
| **HB20** | F primer | CACCATGTATGTGTTTGATCCAACAAC |
| | R primer | TCAGTTGAACTGGTGGTGGTGGTG |
| **Myb-like** | F primer | ATGGAAGCAGACAACGGAGGC |
| | R primer | TCAAAGATTATCTCCTGAAACCTTATCCA |
| **RHD6** | F primer | ATGGTCTCATTTCGTTGGGAT |
| | R primer | TTAATTGGTGATCAGATTCGAATTCCTG |
| **RSL4** | F primer | AAGGTCTCTCACCATGGACGTTTTTGTTGATGGTGAA |
| | R primer | TCACATAAGCCGAGACAAAAGGTTGTG |
